# Site-specific phosphorylation affects the structure and interactions of the Ycf1p R region

**DOI:** 10.64898/2026.01.07.698183

**Authors:** Sarah E.S. Quail, Sarah C. Bickers, Agatha Tymczak, Voula Kanelis

## Abstract

Many ATP binding cassette (ABC) proteins function in active transport of solutes across biological membranes. At minimum, ABC proteins are composed of two repeats, each consisting of a transmembrane domain (TMD) and a nucleotide binding domain (NBD). In many ABC proteins, the TMD-NBD halves are connected by an intrinsically disordered linker that regulates the activity of the ABC protein through phosphorylation. These regulatory (R) regions are often invisible or at low-resolution in electron cryo-microscopy maps, and thus information about how R region phosphorylation controls ABC transporter activity is missing. Here we employ nuclear magnetic resonance (NMR) spectroscopy to discern the structural features and interactions of the R region from the yeast cadmium factor 1 protein (Ycf1p), a member of the C subfamily of ABC proteins that is homologous to human multidrug resistance protein 1. Our data show that the entire R region possesses residual secondary structure that changes with phosphorylation, including for often-invisible R region segments. The data demonstrate R region interactions with NBD1 and, for the first time, NBD2. NBD/R region interactions depend on the phosphorylation state of the R region and the nucleotide-bound and oligomeric states of the NBDs, indicating how R region interactions change in the transport cycle. Complementary biochemical studies show that R region phosphorylation affects the ATPase activity of the NBDs. The molecular-level information provided by the solution NMR studies presented allow for a more complete model of how R region phosphorylation modulates the activities of Ycf1p and related ABC proteins.

**Significance:** NMR data on the intrinsically disordered R region from Ycf1p show that the disordered R region possesses residual structure that is altered by phosphorylation. Phosphorylation of the R region also modulates its interactions with the NBDs, as does the nucleotide-bound and oligomeric states of the NBDs themselves. The molecular-level information regarding the structural features of the R region and its complexes with the NBDs provide sought-after information regarding R region states throughout the transport cycle. The data presented here provide a more complete description of how phosphorylation of the often-invisible R region controls ABC protein activity.

## Introduction

The ATP-binding cassette (ABC) superfamily of membrane proteins is ubiquitous in nature (1–3). ABC proteins are comprised, at minimum, of two half-transporters that each consist of a transmembrane domain (TMD) and a nucleotide binding domain (NBD) (Fig. 1A). Binding and hydrolysis of ATP at the NBDs enables ABC proteins to actively transport solutes across biological membranes, function as channels, or regulate activities of other proteins. Many ABC proteins in which the TMDs and NBDs are fused in a single polypeptide also possess a large intrinsically disordered linker that connects the two halves and that regulates the activity of ABC proteins (4, 5). Thus, the arrangement of the ABC core of many single polypeptide ABC proteins follows the sequence TMD1-NBD1-R-TMD2-NBD2, where R denotes the disordered regulatory region (Fig. 1A and Supplementary Fig. 1).

**Figure 1.**
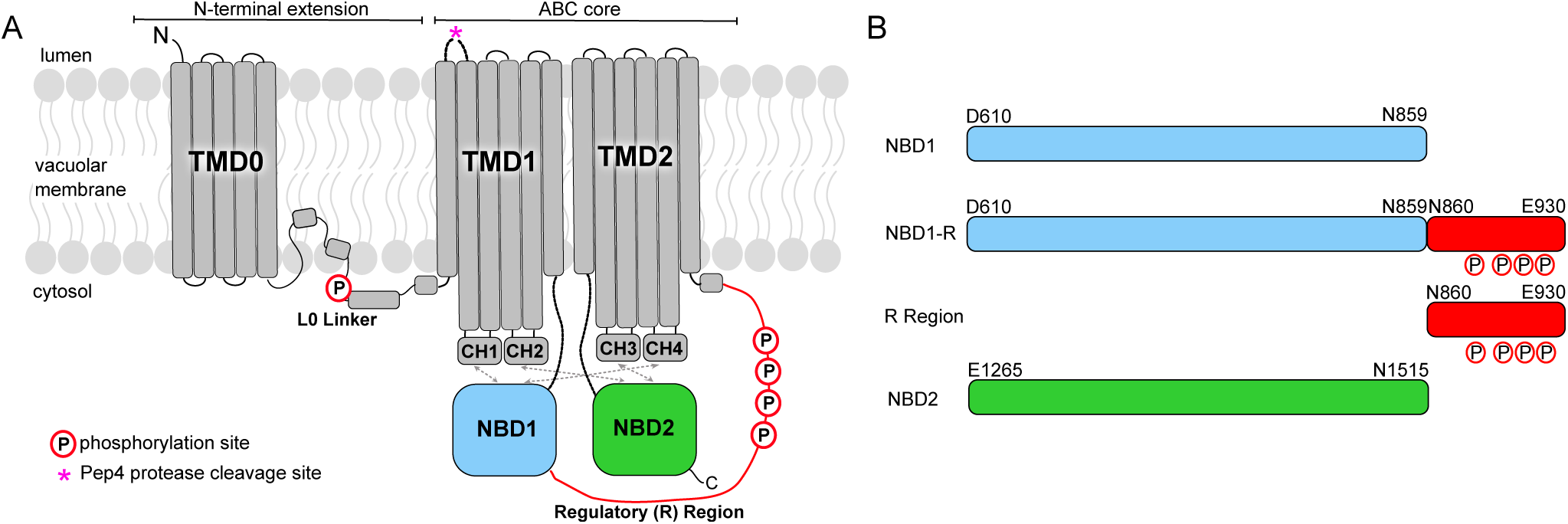
Yeast cadmium factor 1 protein (Ycf1p) is a C-subfamily ABC (ABCC) transporter. (A) Schematic diagram of Ycf1p showing the transmembrane domains (TMD0, TMD1, TMD2) and L0 linker in grey, nucleotide binding domain 1 (NBD1) in blue, NBD2 in green, and the intrinsically disordered regulatory (R) region in red. Coupling helices (CH1-CH4) link the cytoplasmic extensions of transmembrane helices in TMD1 and TMD2 and interact with the NBDs. The two half transporters of the ABC core in Ycf1p are comprised of TMD1-NBD1 and TMD2-NBD2. Phosphorylation sites in the L0 linker (S251) (13) and in the R region (S903, S908, T911, and S914) (14, 20) are indicated with a ‘P’ circled in red. The Pep4 protease cleavage site is denoted by a pink asterisk (*). (B) Schematic representation of proteins used in the present study. As in other studies, NBD1 consists of residues D610-N859 (16, 20), whereas NBD2 is comprised of residues E1265-N1515 (35). NBD1-R consists of residues D610-E930, and includes NBD1 and the R region as a single polypeptide (16, 20), leaving the R region to be defined as N860-E930. Residues S903, S908, T911, and S914 are highlighted in Supplementary Fig. 1.

ABC proteins in eukaryotes are divided into seven subfamilies (ABCA-ABCG) based on the sequences of the TMDs and NBDs (1, 3). The C subfamily of ABC proteins (ABCC), which comprises the most functionally diverse group of ABC proteins, contains an ion channel known as the cystic fibrosis transmembrane conductance regulator (CFTR), the sulfonylurea receptors (SURs) that form the regulatory subunit in ATP-sensitive K+ channels, and several *bona fide* transporters that function as multidrug resistance proteins (MRPs). ABCC proteins also possess an N-terminal extension that consists either of an additional TMD (TMD0) that is linked to the ABC core by the L0 linker or just an L0 tail (Fig. 1A) (1, 2, 6).

The ABCC transporter yeast cadmium factor 1 protein (Ycf1p) is located in the vacuolar membrane of *Saccharomyces cerevisiae* and is responsible for transporting glutathione-conjugated heavy metals (*e.g.,* Cd[II], Hg[II], and As[III]) from the cytosol into the vacuole (7–11). Ycf1p is a homologue of human MRP1 (8, 12), and therefore, contains a TMD0 and L0 linker in addition to the ABC core (Fig. 1A). Ycf1p activity is regulated by multiple mechanisms. Phosphorylation at one site in the L0 linker (S251) decreases Ycf1p activity (13), whereas phosphorylation of up to four sites in the R region (S903, S908, T911, S914) increases activity (14–18) (Fig. 1). Pep4 protease-mediated cleavage of the loop between the first two transmembrane helices of TMD1 of Ycf1p yields the mature state of the protein and affects substrate specificity (19). Pep4-mediated proteolysis also stabilizes the formation of a Ycf1p homodimer, which has lower levels of R region phosphorylation compared to the cleaved Ycf1p monomer (16).

Available cryo-EM structures of Ycf1p have yielded some insights into the molecular basis by which R region controls Ycf1p activity. There are multiple structures of Ycf1p in absence of nucleotide and substrate, and in which the transporter is in the inward-facing conformation with the substrate cavity exposed to the cytoplasm and the NBDs separated (15, 16, 18, 20) (Supplementary Fig. 2A). These structures show phosphorylated R region bound to the peripheral side of NBD1, opposite the ATP binding site and NBD2 dimerization interface. However, density is missing for at least part of the R region, including the phosphorylated residues in some Ycf1p structures, and the structures vary in which R region residues interact with NBD1 (15, 16, 18, 20). As a disordered protein (21, 22), the R region is expected to sample multiple conformations that rapidly interconvert and that can engage in different interactions, leading to the missing cryo-EM density and different NBD1/R region contacts. There is also a structure of dephosphorylated Ycf1p in the inward-facing conformation with part of the R region embedded in the substrate binding cavity (17) (Supplementary Fig. 2B). Because this structure buries the R region phosphorylation sites, there must be a mechanism by which the R region is dislodged from this auto-inhibitory position to enable phosphorylation-dependent activation of Ycf1p. Also not understood are whether structural features of the R region ensemble change when it dissociates from the substrate cavity and/or whether other R region interactions occur.

Structures of other ABCC transporters in the inward-facing conformation also highlight interactions of the non-phosphorylated R region with the substrate-binding cavity (23–25), NBD1 (26–29), and NBD2 (23, 27). However, the R region is often at low-resolution and thus not all of the R region is visible, as seen for auto-inhibited Ycf1p (17). Additionally, R region residues become invisible in structures where NBD1 and NBD2 are dimerized and the ABCC protein is in an outward-facing conformation, regardless of the phosphorylation state of the R region (23, 24). At least for the CFTR Cl^−^ channel and the regulatory SUR protein, nuclear magnetic resonance (NMR) studies have identified additional R region interactions and interactions of other disordered regions, respectively (26, 27, 30).

To uncover the missing molecular features of the differentially phosphorylated forms of the Ycf1p R region, we conducted solution NMR experiments. Our NMR studies show that R region possesses significant residual secondary structure across the entire sequence, including for residues missing in the Ycf1p cryo-EM maps, and that the residual R region structure changes with phosphorylation. Phosphorylation of the R region also affects its interactions with the NBDs, as do the nucleotide-bound and oligomeric states of the NBDs. Together with complementary ATPase assays, the solution NMR structural data enhance our understanding of how R region phosphorylation affects the Ycf1p transport cycle, which can be applied to other ABCC proteins. The work also highlights the synergy between cryo-EM structures of membrane proteins and in-solution structural data of their intrinsically disordered regions, which are found in a significant fraction of ABC transporters (4, 5) and other eukaryotic membrane proteins (22, 31–34).

## Results

### Isolated Ycf1p NBDs and R region can be used in NMR-based studies

Different proteins comprising the NBDs and R region were used in our NMR studies (Fig. 1B). As in previous work (16, 20), NBD1 consists of residues D610-N859 and NBD1-R, in which NBD1 is connected to the R region in a single polypeptide as in full length Ycf1p, is comprised by residues D610-E930. Thus, the isolated R region is defined as N860-E930. We also used isolated NBD2 in this study, which consists of residues E1265-N1515 according to structural data (20) and biochemical information regarding NBD2 sample stability (35).

As expected on the basis of its disordered character, most R region resonances are centered about 8.2 ppm in the ^1^H dimension and possess sharp line-shapes in two-dimensional ^1^H-^15^N correlation spectra (Fig. 2A and Supplementary Fig. 3A). The limited ^1^H^N^ dispersion of R region resonances and their high signal intensities are consist with the view of a disordered R region that samples multiple, rapidly inter-converting conformations so that on average all backbone nuclei are exposed to the solvent and have low relaxation (or signal decay) rates (36–39). Further, many resonances in the spectra of isolated R region overlay with resonances in spectra of NBD1-R (Supplementary Fig. 3A), indicating that R region is disordered in the absence and presence of the neighbouring NBD1. The overlap of most R region resonances between the spectra also suggests that R region residues adopt similar conformations in the absence and presence NBD1. Spectra of NBD1-R also possess resonances with ^1^H^N^ chemical shifts ranging from 7.0-10.0 ppm, which is typical for structured proteins, that overlap with resonances in spectra of isolated NBD1 (Supplementary Fig. 3B). The few differences in NBD1 peaks between the NBD1 and NBD1-R spectra may be derived from residues near the C terminus of NBD1 where R region is attached. Additional differences in NBD1 peaks may be due to transient interactions of the R region with the folded NBD1. These data indicate that we can use isolated NBD1, NBD1-R, or R region proteins in our studies. Note that NMR resonance assignments could only be obtained for the R region (see below). Unfortunately, concentrated NBD1 samples suffer from precipitation over time at the high temperatures (30°C) required to obtain high-quality NMR spectra needed for resonance assignments. In contrast, high-quality ^15^N-^1^H^N^ transverse relaxation optimized spectroscopy (TROSY)-HSQC spectra of NBD1 and NBD1-R can be obtained (Supplementary Fig. 3).

**Figure 2.**
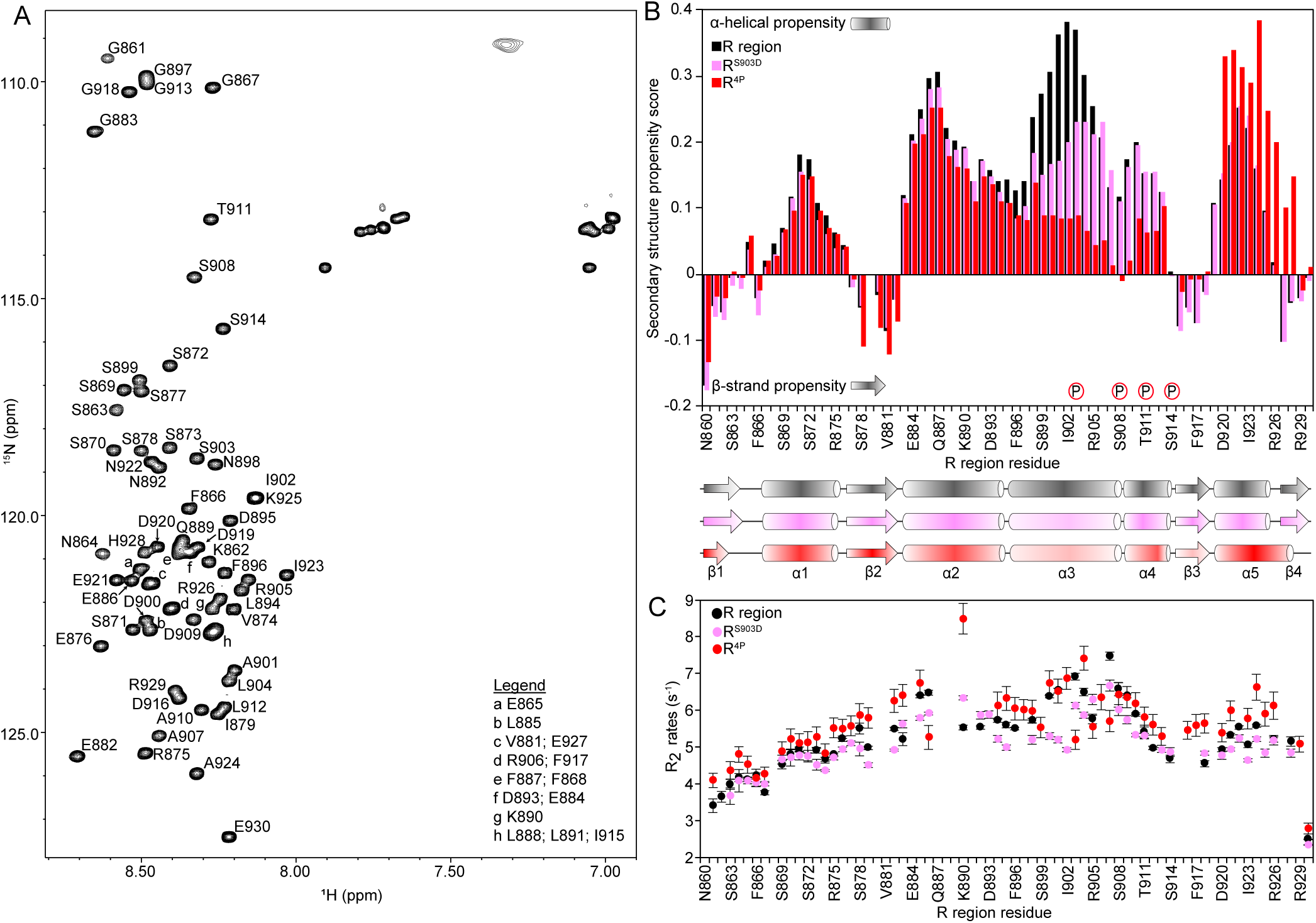
R region possesses residual secondary structure that changes with phosphorylation. (A) ^1^H-^15^N HSQC spectrum of non-phosphorylated R region (257 µM) recorded at 4 °C labeled with resonance assignments. (B) Secondary structure propensity (SSP) scores for the R region (black), R^S903D^ (pink) and R^4P^ (red), which were calculated using ^13^Cα and ^13^Cβ and shown as a function of R region residue number. The phosphorylation sites are indicated by a P circled in red. The absence of a bar indicates that data could not be analyzed for those residues either because the residue is a Pro or is next to a Pro. Schematic representation of the secondary structure, determined by SSP values, for each R region protein are shown below the plot. The colour of the schematic corresponds to the colour used in the SSP plot for each R region protein, whereas the shading represents the magnitude of the SSP value. Cylinders represent consecutive residues with α-helical propensity, whereas arrows denote consecutive residues with β-sheet propensity. (C) ^15^N R_2_ relaxation rates for R region (257 µM, black), R^S903D^ (260 µM, pink), and R^4P^ (190 µM, red). Residues with overlapped resonances or low signal were excluded from the analysis. Selected resonances and decay curves are shown in Supplementary Fig. 7.

The use of *E. coli* for protein expression allows for obtaining samples of R region (and NBD1-R) in the non-phosphorylated state (Supplementary Table 1). While phosphorylated forms of the proteins could be obtained by *in vitro* phosphorylation, this method has some challenges. First, the R region is phosphorylated primarily at four sites (16, 20), with overlapping consensus sequences for different kinases (15, 17), making the generation of fully phosphorylated R region samples difficult (Supplementary Table 2). Samples of R region and NBD1-R with non-uniform phosphorylation would complicate our NMR experiments. Given the flexibility of the disordered R region, even small populations of non-phosphorylated R region or R region that is differentially phosphorylated would be visible in NMR spectra. Non-uniform phosphorylation would lead to multiple resonances from the R region phosphorylation sites and residues nearby. Similar complications would occur for non-uniformly phosphorylated NBD1-R. To circumvent these problems, we employed phosphomimetic mutations to study the effect of phosphorylation in Ycf1p. Two different phosphomimetic R region mutants were used in the work. The R^S903D/S908D/T911E/S914D^ mutant, herein denoted as R^4P^, represents the fully phosphorylated form of the R region in monomeric Ycf1p (16, 18, 20). The R^S903D^ variant is used to study the mono-phosphorylated R region seen in dimeric Ycf1p (16). Previous cell survival assays and biochemical investigations have successfully utilized phosphomimetic mutants to probe the effects of R region phosphorylation (13, 14, 16, 18, 20). Notably, NMR spectra of PKA-treated R region, which results in phosphorylation of S908 and T911 (references (8, 14) and Supplementary Tables 1 and 2), and the R^S908D/T911E^ phosphomimetic mutant display similar chemical shift differences from spectra of non-phosphorylated R region (Supplementary Fig. 4). Thus, phosphomimetic R region mutants can be employed to study phosphorylation-dependent changes in structure and interactions of the R region.

### The Ycf1p R region possesses residual structure that changes with phosphorylation

Obtaining residue-specific information on R region and its interactions with the Ycf1p NBDs requires NMR resonance assignment of the R region (Fig. 2A). In the case of non-phosphorylated R region, we obtained resonance assignments for 98.6% of ^1^H^N^, 97.2% of backbone ^15^N, 100% of ^13^C carbonyl (^13^C’), 98.6% of ^13^Cα, and 100% of ^13^Cβ nuclei. While spectra of non-phosphorylated R region, R^S903D^ and R^4P^ have multiple peaks that overlap, there are significant chemical shift perturbations of many resonances that preclude the straight-forward transfer of assignments between the different spectra (Supplementary Fig. 5A and B). Thus, we also obtained resonance assignments for R^S903D^ (98.6% ^1^H^N^, 97.2% backbone ^15^N, 98.6% ^13^C’, 98.6% ^13^Cα, 100% ^13^Cβ) and for R^4P^ (95.8% ^1^H^N^, 94.4% backbone ^15^N, 95.8% ^13^CO, 97.2% ^13^Cα, 95.4% ^13^Cβ). Note that resonance assignment R^4P^ was achieved by using data on both the isolated R^4P^ and NBD1-R^4P^. Interactions of R^4P^ with NBD1 in the NBD1-R^4P^ protein (20) likely alleviated some of the overlap of the R^4P^ resonances and the ambiguity in some of the resonance connectivities, enabling more resonance assignments of R^4P^ to be obtained.

The available ^13^Cα and ^13^Cβ chemical shifts were used to obtain 2° structure propensity (SSP) scores for R region, R^S903D^, and R^4P^ (Fig. 2B). SSP scores, which are determined from the deviation of the observed chemical shifts of a specific amino acid from its random coil values, provide a measure of the likelihood of an amino acid to adopt an α-helical or β-strand conformation in the protein (40). SSP scores of 1 and −1 reflect stable α-helical and β-strand conformations, respectively, as is observed in folded proteins. For disordered proteins like the R region, the magnitude of SSP scores between −1 and 1 reflect the percentage of conformers that have β-strand or α-helical structure at that position. As evident from the SSP scores, the R region does not adopt a random coil conformation and instead possesses significant residual 2° structure that consists of four β-strands and five α-helices (Fig. 2B, *black bars*). Residues throughout R region possess residual 2° structure, including those invisible in the cryo-EM map of dephosphorylated Ycf1p (17), Notably, part of α3-helix, which has the highest α-helical propensity, as well as the α4- and α5-helices denoted by the SSP analysis align with α-helices modelled in the dephosphorylated Ycf1p structure (Supplementary Fig. 2A and Supplementary Fig. 6) (17), further highlighting that NMR studies of isolated R region can provide insights into its structural features in the full transporter.

We also determined the backbone ^15^N R_2_ relaxation rates of isolated R region (Fig. 2C and Supplementary Fig. 7) (41). R_2_ rates, which measure the rate at which the NMR signal in transverse plane decays, are sensitive to motions on a ns-ps timescale and slow conformational changes on the μs-ms timescale (42). High R_2_ rates indicate slower changes in conformational states or restricted mobility, as is typical for structured proteins. Low R_2_ rates reflect rapid motion, such as those found in disordered proteins in which motions of individual amino acids are generally not affected by neighbouring residues and only by their locations in the polypeptide chain. ^15^N R_2_ rates of disordered proteins typically range from 3-5 s^-1^, with reduced rates at the N- and C-termini (21, 36–38, 43). However, residues in disordered proteins with propensity for 2° structure or that form residual 3° contacts will exhibit elevated ^15^N R_2_ rates for the associated nuclei. Such is the case for the R region, where the ^15^N R_2_ rates vary between 2.4 – 8.5 s^-1^ (Fig. 2C, *black filled circles*). While higher R_2_ rates generally correspond to residues with α-helical propensity, the SSP value and R_2_ rate do not always match, possibly because of 3° contacts within the R region such as those involved in the helix-strand hairpin structure of dephosphorylated R region bound in the Ycf1p substrate-binding cavity (17).

Phosphorylation affects residues throughout the R region. As expected, mono-phosphorylation of the R region, as probed by the R^S903D^ phosphomimetic mutation, causes chemical shift changes in resonances from residues in the α3-helix where S903 is located (Supplementary Fig. 5A). Significant chemical shift changes are also observed for resonances from the α2-helix (Q887-L891) and the C-terminal end of the R region (H928-E930). Note that chemical shift changes in H928-E920 may be from long-range effects of S903 phosphorylation or from slight differences in pH of the samples that could affect the charge state of H928, the only His in R region. Local structural changes with the S903D phosphomimetic mutation result in decreased α-helical propensity of the α3-helix (Fig. 2B, *pink bars*) and lower ^15^N R_2_ relaxation rates for the associated nuclei that indicates their increased mobility (Fig. 2C, *pink filled circles*). Similar structural changes are observed with phosphorylation of other intrinsically disordered regions in ABC proteins (26, 30) or other disordered proteins (44). Long-range structural changes from S903D are exemplified by lower SSP values in α1- and α2-helices and a general decrease of ^15^N R_2_ rates throughout the protein (Fig. 2B and C, *black vs. pink bars and circles, respectively*).

Structural changes are also observed in R^4P^, but in this case the effect is more pronounced. There is a marked decrease in residual α-helical structure of the α3- and α4-helices, which are where the four phosphorylation sites are located (Fig. 2B, *red bars*) and which is consistent with some Ycf1p structures that show extended structure for phosphorylated residues in the R region (15, 18). R^4P^ also exhibits a decrease in the 2° structure propensity of the α1- and α2-helices and the β3-strand. Additionally, R^4P^ possesses an increase in both the α-helical content and length of the α5-helix that includes residues with residual β-strand or extended conformations in non-phosphorylated R region and R^S903D^. In fact, SSP scores for the α5-helix in R^4P^ are comparable to those of the α3-helix in non-phosphorylated R region. Notably, the α5-helix observed in R^4P^ is consistent with the α-helical structure of these residues in some cryo-EM structures of phosphorylated Ycf1p (Supplementary Fig. 6) (15, 18, 45). As with R^S903D^, the far-reaching effects of the phosphomimetic mutations in R^4P^ are also demonstrated by chemical shift differences between non-phosphorylated R region and R^4P^ that extend to α2-helix and α5-helix residues (Supplementary Fig. 5B), and by changes in ^15^N R_2_ rates (Fig. 2C, *filled red circles*). However, rather than a general overall decrease, as seen for R^S903D^, ^15^N R_2_ rates in R^4P^ generally increase compared to non-phosphorylated R region. Increased ^15^N R_2_ rates for residues comprising the α5-helix are likely due, at least partly, to the increased α-helical propensity for these resides in R^4P^. Increased rates for the remaining residues suggest tertiary interactions within R^4P^, given that SSP values for other R^4P^ residues are similar or lower than R and R^S903D^ SSP values. Although Ycf1p structures show interactions of phosphorylated R region with the peripheral side of NBD1 (15, 16, 18, 20, 45), there is evidence that phosphorylated R region can adopt different structures and engage in different interactions in full-length Ycf1p. First, not all of the R region is observed in phosphorylated Ycf1p cryo-EM maps, including the phosphorylated residues (20), and thus intra-R region interactions are possible. Further, there is variability in which R region residues bind NBD1 in the different structures (15, 18, 20, 45), indicating that different NBD1/R region complexes are possible. Together, the cryo-EM structures (Supplementary Figs. 2, 6) (15–18, 20, 45) and NMR data (Fig. 2 and Supplementary Figs. 4-7) highlight the dynamic nature of the R region that would allow this disordered element to adopt different structures in different phosphorylated states.

### R region/NBD1 interactions change when nucleotide binds NBD1 and R region is phosphorylated

Previous work probing changes in fluorescence of Trp residues in NBD1 showed that both non-phosphorylated and phosphorylated R region bind NBD1, and NBD1/R region interactions change with MgATP binding at NBD1 and with R region phosphorylation (16, 20). Because Trp residues are only found in NBD1, these studies probed NBD1/R interactions from the perspective of NBD1. Here we determined which R region residues bind NBD1 with NMR studies in which the R region is visible.

We chose to probe R region/NBD1 interactions primarily using NBD1-R proteins in absence and presence of phosphomimetic mutations. While studies of isolated R region (Fig. 2 and Supplementary Figs. 4-7) provide insights into this disordered region in full-length Ycf1p, the physical link between NBD1 and the R region in Ycf1p (Fig. 1) likely affects the dynamics and interactions of the R region. 2D ^15^N-^1^H-HSQC NMR spectra of 200 µM NBD1-R proteins in the absence and presence of 2 mM MgATP were recorded at 4° C (Fig. 3A and Supplementary Fig. 8). Note that ^15^N-^1^H HSQC spectra of NBD1-R recorded at low temperatures (4 °C) essentially contain R region peaks only, with all (or most) NBD1 peaks being invisible (Supplementary Fig. 9). Notably and for the most part, any NBD1 peaks present in NBD1-R spectra do not overlap with R region peaks. In contrast, ^15^N-^1^H TROSY-HSQC spectra of NBD1-R at higher temperatures (30 °C) contain NBD1 and R region resonances (Supplementary Fig. 3). We therefore include a ‘glow’ around R in the NBD1-R schematics in Fig. 3 to demonstrate that virtually only R region resonances are observed in ^15^N-^1^H HSQC spectra of NBD1-R at 4 °C.

**Figure 3.**
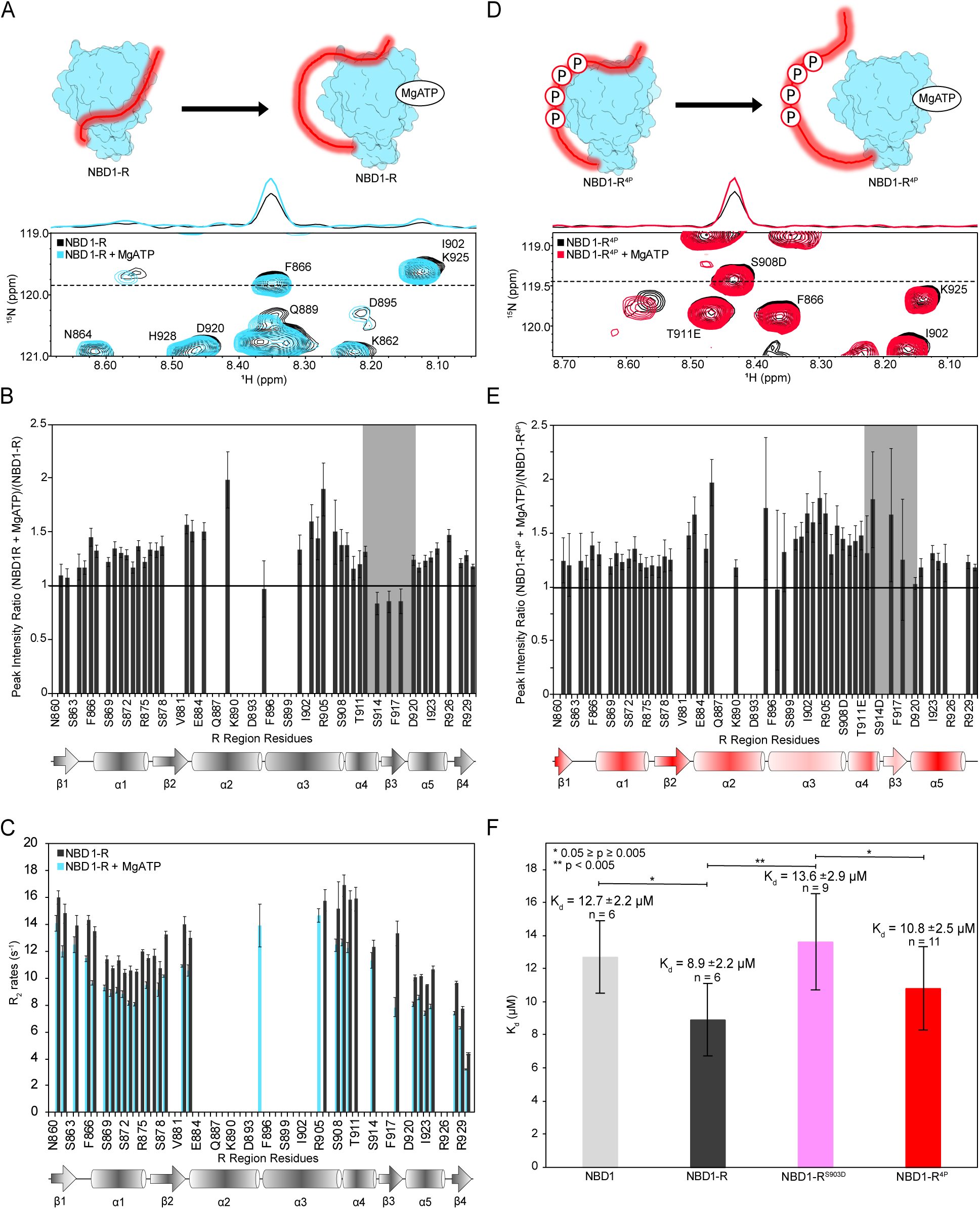
R region interactions with NBD1 change upon nucleotide binding at NBD1 and phosphorylation of the R region. (A) Schematic representation of the NBD1-R samples used in the NMR interaction experiments (top). NMR experiments are performed such that NBD1 signals are silenced and only R region peaks are visible, represented by the red glow around R. A section of the ^1^H-^15^N HSQC spectrum of NBD1-R (200 μM), recorded at 4 °C, in the absence and presence of 2 mM MgATP (bottom). The NBD1-R spectrum in absence of MgATP is shown in black and in the background, whereas the NBD1-R spectrum with MgATP is in light blue and in the foreground. The 1D traces at the top of the spectrum correspond to the dotted line and show the intensities and lineshapes of the F866 resonance. An overlay of the full spectra comparison is shown in Supplementary Figure 9. (B) Plot of the peak intensities ratios for NBD1-R with and without MgATP as a function of R region residues. Peak intensity ratios above 1 indicate increased intensities of the corresponding resonances upon addition of MgATP, while ratios below 1 indicate decreased NMR intensities. Schematic representation of the secondary structure of R region from Fig. 2 is shown below the plot. Light grey shading highlights the lower mobility of residues comprising the residual β3-strand. (C) ^15^N R_2_ relaxation rates for 200 µM NBD1-R in the absence (dark gray) and presence (light blue) of 2 mM MgATP plotted as a function of R region residue. Residues with overlapped resonances or low signal intensities were excluded from the analysis. (D) Schematic representation of the NBD1-R^4P^ samples used in the NMR interaction experiments (top). A section of the ^1^H-^15^N HSQC spectrum of NBD1-R^4P^ (200 μM), recorded at 4 °C, in the absence (black) and presence (red) of 2 mM MgATP (bottom) is shown. The 1D traces show the intensities and line shapes of the resonance corresponding to the S908 phosphomimetic mutation (*i.e.,* S908D) in the absence and presence of MgATP. The full spectra comparison can be found in Supplementary Fig. 14. The grey shading in the plot in this panel and in panel B significant differences in R region residues between MgATP-bound NBD1-R and NBD1-R^4P^. (E) A plot of the peak intensity ratios for NBD1-R^4P^ with and without MgATP. A schematic representation of the secondary structure of R^4P^ is shown below the plot. Light grey shading highlights the increased mobility of residues comprising the residual β3-strand of NBD1-R^4P^ upon binding MgATP, compared to those in NBD1-R in panel B. The data shown in panels A - E are representative of interaction experiments conducted on two separate preparations of NBD1-R or NBD1-R^4P^. (F) Bar graph showing dissociation constants (K_d_ values) for MgATP binding to NBD1, NBD1-R, NBD1-R^S903D^, NBD1-R^4P^. Representative binding curves are shown in Supplementary Fig. 16. K_d_ values were determined from fluorescence binding experiments conducted on protein samples from at least 2 different preparations. The total number of trials (n) indicated above each bar.

Even though MgATP binds NBD1 (Fig. 3A), there are changes to R region resonances (Fig. 3B, and Supplementary Fig.8). Upon addition of MgATP to NBD1-R, R region resonances exhibit chemical shift perturbations (*e.g.,* E886, Q889, R905, S908, I915; Supplementary Fig.8) and/or changes in intensities (*e.g.,* Q889, S903-R905, D920-A924; Fig. 3B). Chemical shift changes reflect the different environments R region nuclei experience, such as would be the case if R region was bound to NBD1 and these NBD1/R region interactions were altered when NBD1 binds MgATP. Changes in the intensities of R region peaks reflect differences in R region mobility in the apo and MgATP-bound states of NBD1-R, as the line shape of an NMR resonance depends on the motion of the associated bond vector (*i.e.,* the ^1^H^N^-^15^N bond), with sharper resonances indicating greater mobility. The fact that most R region resonances increase in intensity upon addition of MgATP suggests that multiple R region residues bind NBD1 in absence of nucleotide and that these NBD1/R region interactions are disrupted when NBD1 binds MgATP. However, not all R region resonances are equally affected. Some peak intensity ratios are close to 1 (*e.g.,* G861, K862, D895, A910), suggesting that some R region residues are not affected by MgATP-binding to NBD1. Peak intensity ratios of less than 1 (*e.g.,* S914, D916, G918 near or in the β3-strand) indicate that some R region residues make greater interactions with MgATP-loaded NBD1 (Fig. 3A, *right* and 3B), consistent with Trp quenching studies of NBD1 and NBD1-R (20). Note that peak intensity ratios could not be determined for resonances that suffer from peak overlap, such as those from some residues in the α2- and α3-helices. Fortunately, reliable peak intensity ratios can be determined for the phosphorylation sites. Data on NBD1-R in the absence and presence of MgATP are consistent with NMR interaction experiments in which NBD1 and R region are separate polypeptides (Supplementary Fig. 10). Note that data shown in Fig. 3 are representative of two experiments repeated on two separate preparations of NBD1-R. Further, data from all other NMR interaction experiments presented in this paper are similarly representative of multiple experiments using separate preparations of protein.

The overall increased mobility of R region resonances in MgATP-loaded NBD1-R is also exemplified by ^15^N R_2_ relaxation experiments (Fig. 3C). Note that, non-TROSY-based ^15^N R_1_ and R_1ρ_ experiments were recorded so that R region peaks are visible and NBD1 peaks are essentially invisible (Supplementary Fig. 11). In general, the ^15^N R_2_ rates of R region peaks are lower when NBD1-R is bound to MgATP compared to apo NBD1-R, consistent with the fact that R region peaks have increased mobility upon addition of MgATP to NBD1-R. Further, and for the most part, R region ^15^N R_2_ rates in NBD1-R are higher than those observed for R region alone and are not uniform across the R region sequence, regardless of the MgATP-loaded state of the protein (Figs. 2C, 3C). As expected, resonances from residues at the N-terminal end of the R region, which is physically connected to NBD1, have high ^15^N R_2_ rates likely because of reduced mobility of these residues compared with the isolated R region. However, elevated R_2_ rates are also seen for resonances from residues throughout R region, such as those between the β2-strand and α2-helix and within in the α3- and α4-helices. Thus, while MgATP binding to NBD1 leads to fewer NBD1/R region interactions, the R region is not completely displaced from NBD1. Instead, multiple R region sites may engage with NBD1, in a dynamic ‘fuzzy’ complex such as that observed for interactions of other IDPs or IDRs with their folded binding partners(46–49), including in other ABC proteins.(26, 27, 30)

With the exception of resonances for N-terminal R region residues, the trend of ^15^N R_2_ rates for R region resonances in NBD1-R (Fig. 3C) match those of R region resonances in the isolated protein (Fig. 2C). One possible explanation of these similarities is that the structural features determined for the isolated R region (*e.g.,* location of residual 2° structures) are conserved in NBD1-R. Binding of other IDRs to their folded interaction partners involves residues in the IDR with fractional 2° structure, with binding enhancing the residual 2° structure. Such situations have been observed for interactions of the R region and NBDs in CFTR (26, 27) and for the L1 linker and NBD1 in SUR2 (30).

We also recorded spectra of NBD1-R^S903D^ and NBD1-R^4P^ in the absence and presence of MgATP (Fig. 3D-E, Supplementary Fig. 12-14). MgATP binding to NBD1-R^4P^ results in increased intensities of R^4P^ region resonances, such as those from residues in the α3-, α4-, and α5-helices and including D916 and G918 that display decreased intensities in MgATP-bound NBD1-R (Fig. 3E and Supplementary Fig. 13). We do note that the error bars in the NBD1-R^4P^ *vs*. MgATP-bound NBD1-R^4P^ intensity ratio plot are higher than those observed for the NBD1-R data, as some resonances are broadened in the NBD1-R^4P^ spectra, possibly because of the higher ^15^N R_2_ rates of R^4P^ (Fig. 2C). Nonetheless, the data indicate a general displacement of R^4P^ from MgATP-bound NBD1. The overall, but slightly increased mobility of R^4P^ residues in the α3-, α4-, and α5-helices in MgATP-bound NBD1-R^4P^ is also seen in the ^15^N R_2_ data (Supplementary 14A). While cryo-EM data show that the tetra-phosphorylated R region binds the peripheral side of NBD1 lacking bound nucleotide (15, 16, 18, 20, 45), it is possible that conformational changes in NBD1 resulting from nucleotide binding affect NBD1/R region interactions. Addition of MgATP to NBD1-R^S903D^ induces similar behaviour of R^S903D^ to that of non-phosphorylated R region when MgATP binds NBD1-R (Fig. 3A, Supplementary Fig. 12 and 14B). Chemical shift changes are observed for some R^S903D^ resonances. Further, most R^S903D^ resonances increase in intensity in the presence of nucleotide and have lower ^15^N R_2_ rates, except those from residues in the α4-helix and β3-strand, reflecting the general increased mobility of R^S903D^ in MgATP-bound NBD1-R^S903D^.

We wanted to determine if differences in how the differentially phosphorylated R region interacts with NBD1 impact the affinity of NBD1 for ATP. The A-loop of NBD1 contains a Trp residue that forms π-π stacking interactions with the adenosine moiety of ATP (Supplementary Fig. 1) (2). Addition of ATP to samples of NBD1 and NBD1-R proteins results in concentration-dependent quenching of the fluorescence of NBD1, which can be used to determine ATP-binding affinities for the proteins (Fig. 3F and Supplementary Fig. 15). Although the ATP-binding affinities of the different NBD1 proteins are similar (K_d_ values from 8.9 – 13.6 µM), there are some statistically significant differences that may be because the different phosphorylated forms of R region mediate distinct interactions with NBD1 (Fig. 3, Supplementary Figs. 8,12-14 and references (15, 16, 18, 20, 45)). NBD1-R has a slightly higher affinity for MgATP compared to NBD1, suggesting that some R region interactions stabilize the nucleotide-bound state of NBD1. The MgATP-binding affinity of NBD1-R is also greater than that of NBD1-R^S903D^ but not NBD1-R^4P^. Because R region phosphorylation increases the ATPase (15–17) and transport activities (17, 18) of Ycf1p, one might expect for NBD1-R^4P^ to have the highest MgATP-binding affinity, which is not the case. It is likely, then, that differentially phosphorylated forms of the R region not only affect MgATP binding at NBD1 but also impact NBD1/NBD2 dimerization and/or ATP hydrolysis. MgATP-binding studies were repeated using at least two separate preparations of protein.

### Differential interactions of non-phosphorylated and phosphorylated R region with NBD2

We wanted to determine if changes in interactions of the R region upon single- and multi-site phosphorylation extend to NBD2. In one structural study, in which Ycf1p was treated *in vitro* with PKA in order to obtain R region with high levels of phosphorylation, low-resolution cryo-EM density attributed to N-terminal residues of R region is observed close to NBD2 (18). However, in all other cryo-EM maps of Ycf1p, with varying levels of phosphorylation, the N-terminal residues of R region are invisible (15, 16, 18, 20). These data further highlight the dynamic nature of R region and suggest that differentially phosphorylated forms of R region mediate different interactions with NBD2. Interactions of non-phosphorylated (and phosphorylated) R region and NBD2 have also been observed for CFTR (27), as have interactions between non-phosphorylated R region and ATP-bound NBD2 in a structure of MRP2 in which the NBDs are separated (23).

In order to obtain additional information on R region/NBD2 interactions, we performed NMR binding experiments using ^15^N-labeled R region proteins and unlabeled NBD2 (Fig. 4A, Supplementary Fig. 16). Note that these interaction studies are done in absence of nucleotide. Addition of NBD2 to R region results in decreased intensities for many R region resonances (Fig. 4B), as observed in spectra of other disordered proteins upon addition of a large and folded binding protein (26, 27, 30, 50), indicating binding of R region to NBD2. As indicated above, NMR resonance line shapes are sensitive to motion of the associated bond vector and are dependent on the R_2_ relaxation rate. Binding to NBD2 causes the interacting residues of R region to tumble more slowly, leading to broadening of the corresponding R region peaks, with the amount of broadening affected by the affinity of the interaction. However, broadening may also be observed for R region residues that do not directly contact NBD2. For example, overall tumbling of an R region residue can be affected if that R region residue interacts with R region residues that bind NBD2. Additionally, changes in intramolecular R region interactions and exchange between free and NBD2-bound conformations would also affect the line shape. These data indicate that multiple R region residues bind NBD2, likely forming a dynamic and “fuzzy” complex (46, 47, 49), and that the strongest NBD2-interacting residues comprise residues in the α3- and α4-helices that include the S903, S908, and T911 phosphorylation sites (Fig. 4B). Because we are measuring intensities of the backbone ^1^H^N^-^15^N resonances, these data suggests a possible conformation for non-phosphorylated R region in Ycf1p that is amenable to phosphorylation. Notably, one of the PKA-binding sites in CFTR is at NBD2 (51), indicating that phosphorylation of NBD2-interacting R region segments, at least by PKA, is plausible. There are little to no interactions of the isolated R^S903D^ or R^4P^ with NBD2, including at the phosphorylation sites (Supplementary Fig. 17), as expected from cryo-EM structures of Ycf1p in which phosphorylated R region binds the peripheral side of NBD1, opposite the NBD2 dimerization interface (Supplementary Fig. 2B and references (15, 16, 18, 20)).

**Figure 4.**
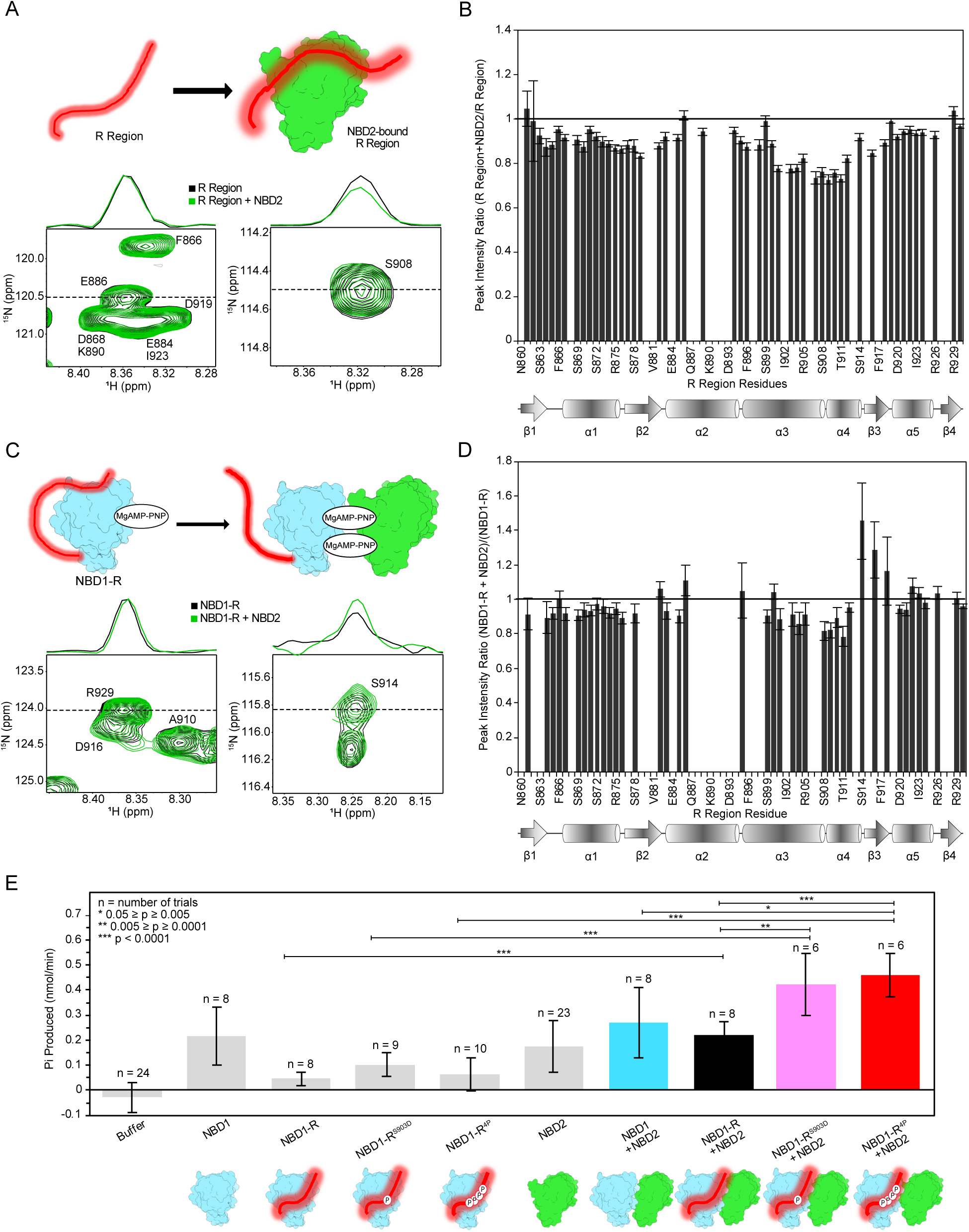
Interactions of non-phosphorylated R region and NBD2 change with NBD1-NBD2 dimerization. (A) Schematic representation of samples used in the R region/NBD2 interaction experiments (top). The R region is ^15^N-labeled, whereas NBD2 is unlabeled. Selected regions of ^1^H-^15^N HSQC spectra of R region (40 µM) recorded at 4 °C and in the absence and presence of NBD2 (160 µM) (bottom). The spectrum of R region is shown in black in the background, while the spectrum of R region in the presence of NBD2 is shown in green and in the foreground. The 1D traces for the E886 and S908 resonances are shown. The overlay of the full spectra is in Supplementary Figure 17. (B) Plot of peak intensity ratios for R region in the absence and presence of NBD2. Peak intensity ratios below 1 indicate decreased intensities of R region resonances upon addition of NBD2, whereas peak intensity ratios above 1 depict increases in R region signals in the presence of NBD2. Schematic diagram of the secondary structure of the R region is below the plot. The data shown are representative experiments repeated on two separate preparations of R region and NBD2. (C) Schematic representation of the samples used in the NBD1-R/NBD2 binding experiments (top). Selected regions of the ^1^H-^15^N HSQC spectra, recorded at 4 °C, of NBD1-R (40 µM) in the presence of 2 mM MgAMP-PNP and without and with NBD2 (160 µM) (top). 1D traces of the R929 and S914 resonances are shown. The overlay of the full spectra is in Supplementary Fig. 19. (D) Plot of the peak intensity ratios for NBD1-R bound to MgAMP-PNP in the absence and presence of NBD2. Schematic representation of the secondary structure of R region is below the plot. The data shown are representative experiments repeated on two separate preparations of NBD1-R and NBD2 proteins. (E) Plot of the rates of inorganic phosphate (P*i*) production as determined by the EnzChek assay (see Methods). A schematic of the protein(s), when included, in each sample is located below the corresponding bar. Experiments were repeated from at least 2 protein preparations with the total number of trials (n) indicated above the bar for each protein sample. Note that the data for NBD1 alone and NBD1/NBD2 were published previously (35), and are shown here for clarity. Data for isolated NBD2 were also shown previously. However, the NBD2 alone data here include additional trials needed as controls for ATPase experiments with NBD2 and NBD1-R proteins. ATPase rate values are shown in Supplementary Fig. 21E.

### NBD1-NBD2 dimerization changes R region interactions and changes with phosphorylation

We extended our R region binding studies to include experiments that probe changes in R region interactions in the context of NBD dimerization. Previous work established that isolated Ycf1p NBD1 and NBD2 proteins associate in a functional dimer in presence of nucleotide (35). Here, we recorded 2D ^1^H-^15^N HSQC spectra of ^15^N-labeled NBD1-R in the absence and presence of unlabeled NBD2 (Fig. 4C). Additionally, samples contained saturating concentrations of MgAMP-PNP, a slowly hydrolyzable analogue of MgATP, to promote stable NBD1/NBD2 dimerization.

Addition of NBD2 to nucleotide-bound NBD1-R results in changes in intensities for multiple R region resonances, particularly those corresponding to residues at and near the phosphorylation sites (Fig. 4C, D and Supplementary Figure 18). The intensity changes indicate that R region residues in the α3- and α4-helices, which include three of the four phosphorylation sites have slightly lower mobilities when NBD1-R binds NBD2, whereas residues in the β3-strand are more mobile. In contrast to that observed for NBD1-R, binding of NBD2 to NBD1-R containing different levels of R region phosphorylation (*i.e.,* as NBD1-R^S903D^, NBD1-R^4P^) does not alter R region interactions (Supplementary Fig. 19), possibly because interactions between NBD1 and the phosphorylated forms of the R region were already disrupted upon nucleotide-binding to NBD1 (Fig. 2D and E; Supplementary Figs. 12-14).

Note that by using MgAMP-PNP in the NMR samples, we are probing the effects of phosphorylation on R region interactions with NBD1 and NBD2 at the beginning of ATPase cycle. Given that conformational changes occur during ATP hydrolysis and subsequent release of ADP and Pi (2), changes in interactions of R region with the NBDs are expected during these reaction steps. Thus, to complement the NMR studies, we assessed the ATPase activities of samples containing our NBD1-R and NBD2 preparations (Fig. 4E and Supplementary Fig. 20) using a colorimetric assay (52). These studies were done using at least two preparations of NBD1, all NBD-R, and NBD2 proteins.

As expected and shown previously for NBD1 (35), the ATPase activities of the individual NBD proteins are relatively low. Further, the ATPase activity level of the NBD1-R/NBD2 complexes are equal to the sum of the ATPase activities of the individual proteins, also seen for Ycf1p NBD1/NBD2 (35) and MRP1 NBD1/NBD2 (53) heterodimers. In contrast, complexes between NBD2 and NBD1-R proteins with phosphomimetic mutations have higher ATPase activities that are also greater than the sum of the ATPase activities for the individual proteins. Notably, the S903D substitution, which is a proxy for mono-phosphorylation of the R region at S903, is sufficient to increase the ATPase activity of NBD1-R/NBD2. These data are consistent with increased ATPase activities of full-length Ycf1p with an S903E phosphomimetic mutation and with yeast viability data showing that removing the S903 phosphorylation site reduces the ability of yeast to grow on high concentrations of cadmium (17). The greater ATPase activities of complexes between NBD1-R^S903D^ or NBD1-R^4P^ with NBD2 may result, in part, because phosphorylation of the R region disrupts its inhibitory interactions with NBD1 and/or NBD2. However, such a model in which non-phosphorylated R region competes with productive NBD1/NBD2 interactions suggests that the NBD1/NBD2 complex should have the highest ATPase activity, which is not the case. Instead, the ATPase hydrolysis rate of the NBD1/NBD2 complex is very similar to that of NBD1-R/NBD2. One possible explanation is that phosphorylated R region stabilizes the NBD1/NBD2 dimer during some steps of the ATPase reaction cycle and/or that phosphorylated R region promotes dissociation of the NBD1/NBD2 dimer after ATP hydrolysis. Together, the NMR and ATPase data further highlight the significant contribution of the R region to Ycf1p function.

## Discussion

In this paper, we present NMR spectroscopy studies that provide residue-specific information on the structural features of the intrinsically disordered R region and its complexes with the NBDs in Ycf1p (Supplementary Fig. 21). The NMR data highlight structural features and interactions of the R region that are consistent with Ycf1p cryo-EM structures, while also providing unique information. Using information from NMR chemical shifts, we show that the entire R region possesses residual 2° structure that changes with phosphorylation, with greater changes elicited by greater levels of phosphorylation and/or inclusion of phosphomimetic mutations. The changes are not just limited to the α3- and α4-helices where the phosphorylation sites are located, but extend to other secondary structure elements, indicating 3° contacts within R region. Where comparisons can be made, the 2° structure propensity of non-phosphorylated R region obtained from NMR studies is consistent with Ycf1p cryo-EM structures (17). The NMR data also show that both non-phosphorylated and phosphorylated R region bind NBD1, and that these interactions change when NBD1 binds MgATP. Further, we show that only non-phosphorylated R region binds NBD2, a result that is consistent with cryo-EM structures of Ycf1p that show binding of phosphorylated R region to the peripheral side of NBD1 opposite the NBD2 dimerization interface (15, 16, 18, 20). As with other NBD complexes involving disordered regions (26, 27, 30), residues in R region with α-helical propensity bind NBD1 and NBD2, particularly residues in the α3- and α4-helices. Our biochemical data show that R phosphorylation leads to greater ATPase activity of complexes between NBD2 and NBD1-R, consistent with functional data using full-length Ycf1p (14, 18).

The data presented here add to previous work (17) to define a model linking R region phosphorylation to Ycf1p activity (Fig. 5). While non-phosphorylated R region forms a helix-strand hairpin motif that occupies the substrate binding cavity (17) (Fig. 5A), interactions of the non-phosphorylated R region and the NBDs also occur. For example, the unobserved R region residues in dephosphorylated Ycf1p may interact with NBD1 and/or NBD2 (Fig. 5A and B, *top*). For phosphorylation to occur, the R region must disengage from the substrate cavity, which may occur stochastically (Fig. 5B, *top*), given the flexibility of the R region and its transient interactions. Alternatively, binding of substrate could out compete the R region from its auto-inhibitory position, given that binding of the R region significantly reduces the volume of the substrate cavity and residues in the substrate cavity interact with both R region and GSSG (17, 45) (Fig. 5B, *bottom*). Each mechanism has been observed in other ABCC proteins (23–25, 54), with the latter mechanism ultimately resulting in basal levels of transport. Non-phosphorylated R region that has dislodged from the substrate-binding site would be free to interact with both NBD1 and NBD2, and to adopt a conformation that exposes R region phosphorylation sites to the cytosol, making them available to kinases (Fig. 5B). Such a scenario is consistent with structural data on CFTR showing PKA binding at NBD2 (51). Once phosphorylated, R region relocates to the periphery of NBD1, opposite the ATP and NBD2 binding sites (Fig. 5C). The disordered nature of the R region enables different R region residues to interact with multiple Ycf1p sites. Two possible R region interaction schemes, based on cryo-EM structures and NMR data, are shown. Binding of phosphorylated R region to the peripheral side of NBD1 allows for ATP binding at the NBDs, NBD dimerization, and subsequent ATP hydrolysis that is required for substrate transport (Fig. 5D and E). Our NMR data also show that R region transiently interacts with NBD1 upon ATP binding and NBD dimerization, allowing for increased ATP hydrolysis upon phosphorylation leading to enhanced substrate transport. Release of ADP and Pi resets Ycf1p to the inward open conformation to allow for transport of another substrate molecule, as for other ABC transporters (2).

**Figure 5.**
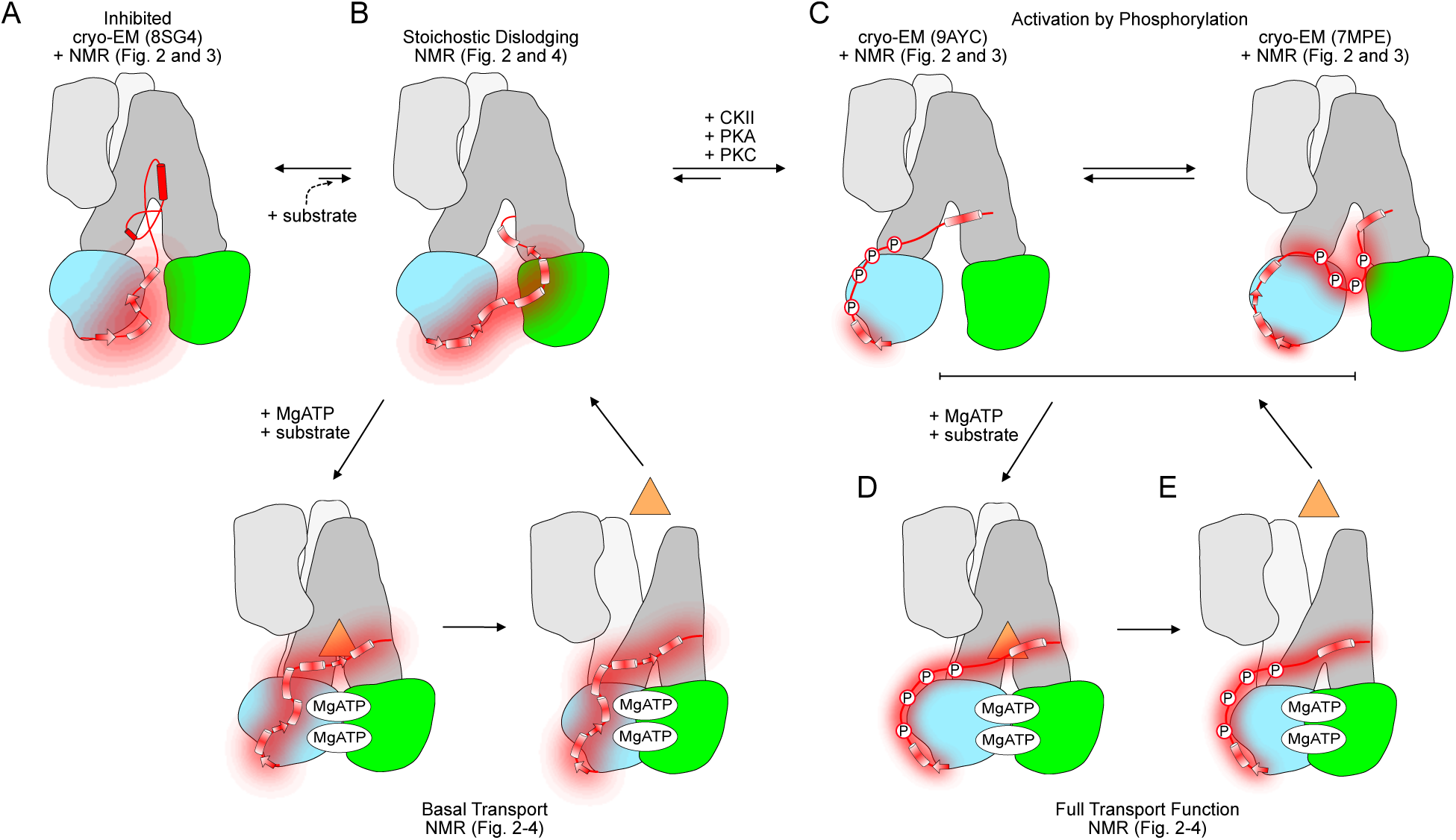
Model of R region interactions throughout Ycf1p transport cycle. (A) In the dephosphorylated Ycf1p cryo-EM structure, R region occupies the substrate binding cavity in a helix-strand hairpin conformation (17). In this state, the phosphorylation sites are in the substrate binding cavity and would need to be dislodged in order for phosphorylation to occur. Trp fluorescence quenching data show that non-phosphorylated R region can also interact with NBD1 near the ATP binding site (16, 20), consistent with NMR data presented here. Thus, segments of the R region not observed in the cryo-EM structure of the auto-inhibited Ycf1p are shown with their possible NBD1 interactions and residual 2° structures. (B) NMR data in this paper also show that R region residues comprising α3- and α4-helices interact with NBD2 and can represent a state to allow for R region phosphorylation. Dislodging of the R region from the substrate cavity may occur either stochastically or be induced by substrate binding, as in other ABCC proteins (23–25, 54). The latter case may result in basal levels of transport, provided that the non-phosphorylated R region adopts a conformation that allows for nucleotide binding and dimerization of NBD1 and NBD2, as seen in our NMR data. The ‘fuzziness’ around the entire R region represents the limited interactions observed for the R region with the nucleotide-bound and dimerized NBDs from the NMR data. (C) Upon phosphorylation, the R region relocates to the periphery of NBD1 opposite the ATP and NBD2 binding sites. Different cryo-EM structures show that different residues of phosphorylated R region can interact with NBD1 (15, 16, 18, 20), and is depicted with the two models. Further, parts of the R region are also missing in these structures. Thus, the phosphorylated R region can engage in different interactions with NBD1, leading to a fuzzy NBD1/R region complex. Potential interactions of non-observed R region residues (at the N-terminal end and comprising the phosphorylation sites) are shown as fuzzy elements. (D) Regardless, movement of the R region to the peripheral side of NBD1 allows for substrate binding and formation of an NBD1/NBD2 MgATP sandwich dimer. (E) Formation of the outward facing conformation follows, leading to transport. Note that all of the phosphorylated R region is shown as a fuzzy curve to highlight the fact that NBD1/R region interactions are disrupted when NBD1 binds MgATP and dimerizes with NBD2. As with other ABC transporters (2), MgATP hydrolysis at the NBD2 composite site resets that transporter for another transport cycle. The model presented here shows possible interactions of the R region at different stages of the ATP hydrolysis and transport cycles. Note that due to the dynamic nature of the disordered R region, interactions of R region with the substrate-binding cavity and with the NBDs are likely transient.

The proposed model by which R region regulates Ycf1p activity is applicable to other ABCC proteins, given that phosphorylation is a common regulatory mechanism for ABCC proteins (55) and that there are phosphorylation sites in the R region of many ABCC proteins (4, 5, 24, 25, 56–59). Cryo-EM structures of ABCC proteins throughout the transport cycle share similarities in interactions of their R regions with those observed in Ycf1p. For example, cryo-EM structural studies of CFTR show phosphorylation-dependent interactions of R region with other parts of the protein (28, 29, 58, 59). NMR binding data show that interactions of CFTR R region with NBD1 and NBD2 are largely disrupted when R region is phosphorylated (26, 27). Cryo-EM structures of CFTR also show that segments of non-phosphorylated R region can insert between the two halves of the protein when CFTR is in the inward-facing conformation, inhibiting NBD1/NBD2 dimerization (28, 29). Phosphorylated R region is repositioned from its inhibitory location to the peripheral side of NBD1 in CFTR in which NBD1 and NBD2 are bound to nucleotide and the NBDs are dimerized (58, 59). The low-resolution density for most of the R region, however, precludes knowing which R region residues are involved binding NBD1 from the cryo-EM structures alone.

Structures of rMRP2 and hMRP2 also show R region in the substrate cavity of the protein (23–25). Addition of substrate can dislodge the R region from this auto-inhibitory position. Further, phosphorylation of specific Ser residues in the R region leads to increased activity of MRP2 (25), as seen for Ycf1p (14–18). Notably, MRP2 preparations used to obtain the auto-inhibited structures are phosphorylated at Ser residues that are not observed in the structures and are presumably located outside the substrate cavity (24, 25), indicating that specific R region sites need to be phosphorylated for full activation of the protein. The three auto-inhibited MRP2 structures reveal different segments of the R region, possibly because the R region is flexible within the substate binding cavity. Such flexibility would enable the R region to dissociate from its auto-inhibitory location, leading to ATPase and transport activity even without additional phosphorylation. Notably, one of the MRP2 structures shows part of the R region contacting the bound ATP and Walker A sequence of NBD2 (23), indicating that the Ycf1p R region/NBD2 contacts we observed by NMR occur in other ABCC proteins. Another such ABCC protein is MRP1 considering that it is also regulated by R region phosphorylation (56).

Inhibition of ABCC proteins can involve other intrinsically disordered regions in the protein. Structural studies of MRP5, which does not contain a TMD0, show that auto-inhibition of the protein is mediated by its cytoplasmic N-terminal extension (60). Note, however, that the absence of a TMD0 is not the determining factor in which intrinsically disordered region will provide the auto-inhibitory interactions, given that CFTR lacks a TMD0. As an additional difference to Ycf1p, MRP2 and CFTR, a phosphorylated Ser residue (S60) in MRP5 makes electrostatic interactions with multiple Arg residues in the substrate cavity, indicating that phosphorylation of this Ser enhances auto-inhibition. A structure of a catalytically-inactive MRP5 in which the NBDs form an ATP-sandwich dimer and the transporter is in the outward-facing conformation is lacking density for the inhibitory portion of N-terminal extension. Thus, as with the R region in CFTR (28, 29, 58, 59) and MRP2 (23–25), the N-terminal extension can dislodge from the substrate cavity.

It is possible that different auto-inhibitory interactions also occur in other ABCC proteins and in ABC transporters in different subfamilies. Our work highlighting the phosphorylation-dependent structural features and interactions of the Ycf1p R region, including for residues invisible in Ycf1p cryo-EM maps, and the effect of the R region on ATPase activity lay the foundation for studies on other ABC proteins. Additionally, the correlation of our NMR data on the NBDs and R region with structures of full length Ycf1p exemplifies the complete molecular picture that can be obtained by combining cryo-EM studies of membrane proteins and in-solution studies of their disordered regulatory regions.

## Materials and Methods

### Materials

*E. coli* LOBSTR-BL21 (DE3) RIL was purchased from Kerafast Inc. ^15^N-NH_4_Cl and ^13^C-glucose were purchased from Cambridge Isotope Laboratories Inc. Other growth media components were obtained from BioShop, as were reagents used in the purification buffers. During purification, the adenosine-5-triphosphate (ATP) used was a disodium salt at 98.5% purity that was purchased from BioShop (Cat # ATP007). For NMR spectroscopy and other assays, samples contained ATP at >99% purity purchased from Thermo Fisher Scientific (Cat # R0441) or containing AMP-PNP purchased from Roche (Cat # 10102547001). The stock ATP solution was diluted accordingly, while AMP-PNP stocks were obtained by dissolving the salts in water to 150 mM and adjusting the pH to 7.4. Pierce™ BCA Protein Assay was purchased from Thermo Scientific (Cat # A55864). Enzchek™ Phosphate Assay Kit was purchased from Invitrogen (Cat # E6646).

### Protein Expression

Ycf1p NBD1, NBD1 linked to the regulatory (R) region (NBD1-R), and NBD2 were expressed as fusion proteins with an N-terminal His_6_-SUMO tag, as described previously (16, 20, 35). The isolated R region was also expressed as a His_6_-SUMO fusion protein. Note that NBD1-R and R region proteins were expressed in their wild-type forms or with the following phosphomimetic mutations: S903D, S908D/T911E, or S903D/S908D/T911E/S914D. NBD1-R and R region proteins containing the four phosphomimetic mutations are referred to as NBD1-R^4P^ and R^4P^, respectively. Briefly, the His_6_-SUMO Ycf1p proteins were expressed in *E. coli* LOBSTR-BL21 (DE3) RIL (Kerafast Inc) using a pET26b-derived expression vector (61). Cell cultures were grown in 1 L of 95% (v/v) M9 minimal media containing ^15^N-NH_4_Cl (Cambridge Isotope Laboratories Inc) and 5% (v/v) lysogeny broth (LB) media, or in 1 L M9 minimal media containing ^15^N-NH_4_Cl and ^13^C-glucose (Cambridge Isotope Laboratories Inc). Cell cultures were grown at 37 °C until they reached an OD_600_ of 0.4 – 0.5, at which point the temperature was gradually lowered so that the temperature was 16 – 18 °C when the OD_600_ of the cell culture was 0.8. The cell cultures were then incubated at 16-18 °C for 30 min before the addition of isopropyl-β-D-thiogalactoside, at a final concentration of 750 µM, to induce expression of the His_6_-SUMO Ycf1p fusion proteins. Cells were harvested 18-20 h post-induction by centrifugation at 4 °C and 2,150 g for 30 minutes and the resulting cell pellets were stored at −20 °C until purification.

### Purification of NBD1, NBD1-R, NBD1-R^S903D^, NBD1-R^4P^

Purification of NBD1, NBD1-R, NBD1-R^S903D^, and NBD1-R^4P^ proteins were achieved using previously-described methods (16, 20) which are described here. All steps were conducted at 4 °C. Cell pellets from 1 L of cell culture were resuspended in 15 mL of lysis buffer (20 mM Tris HCl, 10% [v/v] glycerol, 150 mM NaCl, 5 mM imidazole, 100 mM arginine, 0.2% [v/v] Triton X-100, 1 mg/mL lysozyme, 2 mg/mL deoxycholic acid, 5 mM benzamidine, 5 mM ε-amino n-caproic acid, and 1 mM PMSF, pH 7.6) that also contained 2 mM β-mercaptoethanol, 15 mM MgCl_2_, and 15 mM ATP, and incubated on ice for 15-20 min. A small amount of DNase I was added and the solution was further incubated for another 15 min. The cells were then lysed by sonication (Misonix Moicroson Ultrasonic Cell Disruptor), and the soluble and insoluble fractions were separated by centrifugation at 15,000 g for 30 min. The soluble cell lysate was set aside while the insoluble cell pellet was resuspended in 15 mL of lysis buffer, incubated for 15-20 min, then subjected to sonication and centrifugation. The soluble cell lysates from the first and second sonication steps were combined, filtered using 0.45 µm syringe filter (Pall), and applied to a 5 mL Ni^2+^-NTA affinity column (GE Healthcare) that was pre-equilibrated with 20 mM Tris HCl, 10% (v/v) glycerol, 150 mM NaCl, 5 mM imidazole, pH 7.6. The column was then washed with 5 column volumes of Ni^2+^-column equilibration buffer that also contained 5 mM MgCl_2_ and 5 mM ATP. The His_6_-SUMO-NBD1 (or NBD1-R) fusion proteins were subsequently eluted in five 5-mL fractions with elution buffer (20 mM Tris HCl, 10% [v/v] glycerol, 150 mM NaCl, 500 mM imidazole, 1 mM ε-amino n-caproic acid, pH 7.6) that also contained 15 mM MgCl_2_ and 15 mM ATP. Each 5-mL elution fraction was immediately diluted threefold with dilution buffer (20 mM Tris HCl, 10% [v/v] glycerol, 1 mM benzamidine, 1 mM ε-amino n-caproic acid, pH 7.6) that also contained 15 mM MgCl_2_, 15 mM ATP, and 2 mM β-mercaptoethanol. Elution fractions containing the His_6_-SUMO-NBD1 (or NBD1-R) protein, as confirmed by SDS-PAGE, were dialyzed overnight against a buffer containing 20 mM Tris HCl, 10% (v/v) glycerol, 150 mM NaCl, 2 mM β-mercaptoethanol, 15 mM MgCl_2_, 15 mM ATP, 1 mM benzamidine, 1 mM ε-amino n-caproic acid, pH 7.6. The dialysis bag also included 3.2 µg of His_6_-Ulp1 protease per 1 mL of protein solution to remove the His_6_-SUMO tag from the NBD1 or NBD1-R protein. After dialysis, the NBD1 (or NBD1-R) was removed from the His_6_-SUMO tag and His_6_-Ulp1 protease using a 3-mL reverse Co^2+^-agarose affinity column (Thermo Scientific) that was pre-equilibrated with Co^2+^-column wash buffer (20 mM Tris HCl, 10% glycerol [v/v], 150 mM NaCl, 1 mM benzamidine, 1 mM ε-amino n-caproic acid, pH 7.6) that also contained 10 mM MgCl_2_ and 10 mM ATP. The column was then washed with 5 column volumes of the Co^2+^-column wash buffer. The flowthrough and wash fractions were concentrated to 5-6 mL (so that the protein concentration was 200 – 250 µM) and applied to a size exclusion column (Superdex 75, GE Healthcare) in the NBD1 NMR buffer (20 mM Na_3_PO_4_, 150 mM NaCl, 2% [v/v] glycerol, 2 mM β-mercaptoethanol, 100 µM benzamidine, and 100 µM ε-amino n-caproic acid, pH 7.3) in the absence or presence of 2 mM MgCl_2_ and 2 mM ATP, as required.

Purified samples of NBD1 were either used immediately in NMR studies or flash frozen and kept at −80 °C until use in ATPase assays. In the latter case, the NBD1 sample contained 2 mM MgCl_2_ and 2 mM ATP prior to freezing. The sample was thawed in an icy-water bath (~7 °C). Our ability to freeze NBD1 samples was determined from ^15^N-^1^H TROSY-HSQC spectra, size exclusion column traces, and ATPase assays that showed identical behaviour for the freshly purified and frozen NBD1. In contrast, purified samples of NBD1-R were used immediately.

Samples of NBD1 and NBD1-R with MgATP or MgAMP-PNP for NMR studies were prepared by adding the MgCl_2_ and ATP or AMP-PNP, at final concentrations of 2 mM, to protein samples in the NBD1 NMR buffer. To prepare NBD1 and NBD1-R proteins for ATP binding studies and ATPase assays, the purified proteins were concentrated to ~250 μM and then applied to the size exclusion column pre-equilibrated in the ATPase reaction buffer (50 mM Tris HCl, 5% (v/v) glycerol, 150 mM NaCl, pH 7.6).

### Purification of R region, R^S903D^, R^4P^

Purification of the R region and R region variants followed the general protocol used for purification of the NBD1 and NBD1-R proteins. All steps were conducted at 4 °C. Cell pellets from 1L of cell culture were resuspended in 15 mL of lysis buffer and incubated on ice for 15-20 min. A small amount of DNase I was added, and the solution was incubated for another 15 min before lysis of the cells by sonication (Misonix Moicroson Ultrasonic Cell Disruptor). The mixture was then subjected to centrifugation (at 15,000 g for 30 min) to separate the soluble and insoluble fractions. The insoluble fraction was resuspended in 15 mL of lysis buffer, incubated for 15-20 min, after which the sonication and centrifugation steps were repeated. The cell lysates from the first and second sonication steps were combined, filtered using 0.45 µm syringe filter (Pall), and applied to a 5mL Ni^2+^-NTA affinity column (GE Healthcare) that was pre-equilibrated with equilibration buffer. The column was washed with 5 column volumes of equilibration buffer and the His_6_-SUMO-R region fusion proteins were eluted in five 5-mL fractions with the Ni^2+^-NTA affinity column elution buffer. Each 5-mL elution fraction was collected separately and immediately diluted threefold with dilution buffer. His_6_-Ulp1 protease, at a final concentration of 3.2 µg/mL, was added to each elution fraction containing the His_6_-SUMO-R region protein (as confirmed by SDS-Page). The resulting mixture was concentrated, so that the total protein concentration was 200-250 µM, and applied to a size exclusion column (Superdex 75, GE Healthcare) in 20 mM Tris HCl, 10% [v/v] glycerol, 150 mM NaCl, 1 mM benzamidine, 1 mM ε-amino n-caproic acid, pH 7.6. The fractions containing the R region (or R^S903D^ or R^4P^) were then applied to a 3-mL reverse Co^2+^-agarose affinity column (Thermo Scientific), which was pre-equilibrated with Co^2+^-column wash buffer, to separate the His_6_-SUMO tag from the R region protein. The column was washed with 5 column volumes of the wash buffer, and the flowthrough and wash fractions were combined and concentrated to 10-15 mL. The resulting R region solution was dialyzed overnight against R region buffer (20 mM Na_3_PO_4_, 150 mM NaCl, 2% [v/v] glycerol, 100 µM benzamidine, and 100 µM ε-amino n-caproic acid, pH 7.3). Note that the R region buffer is identical to the NBD1 buffer lacking β-mercaptoethanol. The concentration of R region samples was determined with a bicinchoninic acid (BCA) assay (Pierce™ BCA Protein Assay; Thermo Scientific). Samples of R region were used immediately after preparation.

### Purification of NBD2

Purification and storage of NBD2 was conducted using a previously established method (35), as described briefly here. All steps were conducted at 4 °C. The cell pellet from 1 L of cell culture was resuspended 25 mL of lysis buffer that contained 2 mM β-mercaptoethanol and had 300 mM NaCl in place of 150 mM NaCl used in the NBD1 and R region protein purifications. The subsequent mixture was incubated on ice for 15-20 min, at which point a small amount of DNase I was added and the mixture was incubated on ice for another 15 min. The cells were then lysed by sonication (Misonix Moicroson Ultrasonic Cell Disruptor), and the soluble and insoluble fractions were subsequently separated by centrifugation at 15,000 g for 30 min. The resulting pellet was resuspended in 15 mL of lysis buffer, and the mixture was incubated on ice for 15-20 min before being subjected to another round of sonication and centrifugation. The cell lysates were combined and filtered using 0.45 µm syringe filter (Pall) and applied to a 10 mL Ni^2+^-NTA affinity column (GE Healthcare) that was pre-equilibrated with 20 mM Tris HCl, 10% (v/v) glycerol, 150 mM NaCl, 5 mM imidazole, pH 7.6. The column was washed with 5 column volumes of equilibration buffer and the His_6_-SUMO-NBD2 protein was eluted with 5 column volumes of the Ni^2+^ column elution buffer used above (*i.e.,* for Ni^2+^-affinity purification of His_6_-SUMO-R). Each 10 mL-fraction was collected separately and immediately diluted threefold with dilution buffer that also contained 2 mM β-mercaptoethanol. Elution fractions containing the fusion protein were pooled and concentrated to 5-6 mL (so that the protein concentration was 200-250 µM) and applied to a size exclusion column (Superdex 200, GE Healthcare) in 20 mM Tris HCl, 10% (v/v) glycerol, 150 mM NaCl, 2 mM β-mercaptoethanol, 1 mM benzamidine, 1 mM ε-amino n-caproic acid, pH 7.6. Note that protein concentrations greater than 250 µM prior to size exclusion chromatography lead to increased dimerization of His_6_-SUMO-NBD2. The fractions containing His_6_-SUMO-NBD2, as confirmed by SDS-PAGE, were pooled and concentrated to 25-30 mL. Approximately 1.8 mg, in total, of His_6_-Ulp1 protease was then added to the purified His_6_-SUMO-NBD2. The NBD2 was then separated from the His_6_-SUMO tag and from His_6_-Ulp1 protease using a reverse Co^2+^-agarose affinity step.

For NMR spectroscopy studies, the mixture containing His_6_-SUMO and NBD2 was applied to a Co^2+^-agarose affinity column (Thermo Scientific) pre-equilibrated with Co^2+^-column buffer used above. The column was then washed with 5 column volumes of wash buffer. The flowthrough and wash fractions contained NBD2. The purified NBD2 was then dialyzed overnight against the NMR NBD2 buffer (20 mM Na_3_PO_4_, 150 mM NaCl, 10% (v/v) glycerol, 1 mM benzamidine, 1 mM ε-amino n-caproic acid, 2mM β-mercaptoethanol, pH 7.3). A high concentration of glycerol was used to ensure sample stability during the NMR experiments. For ATPase activity studies, the His_6_-SUMO and NBD2 mixture was applied to a Co^2+^-agarose affinity column (Thermo Scientific) that was pre-equilibrated with the ATPase reaction buffer plus 1 mM benzamidine and 1 mM ε-amino n-caproic acid. The column was then washed with 5 column volumes of this buffer. The flowthrough and wash fractions containing NBD2 were then pooled and concentrated to ~50 µM NBD2. Aliquots of purified NBD2 of 300 µL were prepared, flash frozen with liquid nitrogen, and stored at −80 °C until use in the assay. Protein concentration samples of NBD2 were determined by an A_280_ measurement using an extinction coefficient (ε) of 18,450 M^-1^cm^-1^.

### NMR Spectroscopy

All experiments were recorded on a 600 MHz Agilent VNMRS or 600 MHz Agilent INOVA spectrometer equipped with actively-shielded z-gradients. The 600 MHz VNMRS spectrometer was equipped with an HCN room temperature, while the 600 MHz INOVA spectrometer was equipped with a H/FCN or HCN cryoprobe, or and H/FCN room temperature probe. All NMR samples contained 1 mM 4,4-dimethyl-4-silapentant-sulfonic acid (DSS) for chemical shift referencing (62). NMR spectral data were processed using NMRPipe/NMRDraw (63) and analyzed with NMRView (64).

#### NMR Assignments of R Region, R^S903D^, and R^4P^

All NMR resonance assignment experiments were collected at 4°C. Backbone ^1^H, ^15^N, ^13^C, ^13^Cα, and side chain ^13^Cβ resonance assignments for R region, R^S903D^, and R^4P^ were obtained from standard triple resonance experiments that correlate Cα and Cβ, or carbonyl C chemical shifts in sequential residues, a triple resonance experiment that correlates the aliphatic carbon chemical shifts in one residue with the ^15^N and ^1^H^N^ chemical shifts in the next residue, as well as an ^15^N-NOESY experiment (65, 66). These experiments were recorded on 250µM protein samples that were uniformly labelled with ^15^N and ^13^C. The R region and R^4P^ data were supplemented with ^15^N-^1^H HSQC spectra on protein samples that contained ^15^N-Leu and were otherwise unlabeled. Resonance assignment experiments were also recorded on NBD1-R^4P^, but in this case experiments that correlate chemical shifts of ^15^N nuclei in sequential residues and are advantageous for disordered proteins (67) were also used in addition to the triple resonance data. Note that resonance assignment experiments of NBD1-R^4P^ were recorded with pulse sequences used for small proteins so that R^4P^ resonances were visible and NBD1 resonances were invisible.

#### Chemical shift differences

The differences in the chemical shifts of resonances between two spectra (*e.g.,* R region vs R^S903D^) were calculated using the equation 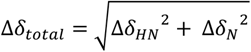, where Δ*δ_total_* is the combined chemical shift difference in Hz, and Δ*δ_HN_* and Δ*δ_N_* are the differences in the ^1^H and ^15^N chemical shifts, respectively.

#### Relaxation Experiments

^15^N R_1_ and ^15^N R_1ρ_ relaxation experiments were performed at 4 °C using previously described pulse sequences (41, 68). ^15^N relaxation experiments were recorded on a sample of 250 µM R region (and R^S903D^ and R^4P^), or a sample of 200 µM NBD1-R (and NBD1-R^S903D^ and NBD1-R^4P^) in the absence and presence of 2 mM MgATP. ^15^N R_1_ rates were obtained from 8 spectra with relaxation delays of 10.1, 50.4, 100.9, 181.6, 262.3, 333.9, 433.8, 554.8 ms. ^15^N R_1ρ_ values were measured from eight different spectra recorded with delays of 10, 20, 30, 40, 50, 80, 90, 100 ms. ^15^N R_2_ values were determined from the R1 and R1ρ values using the equation *R*_2_ = (*R*_1_ − *R*_1*ρ*_*cos*^2^*θ*), where 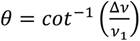 and *ν*_1_ the applied spin-lock rf field and Δ*ν* is the offset Hz. A spin-lock field of 1736 Hz was used. For both the R_1_ and R_1ρ_ data, the Rate Analysis tool in NMRView was used to obtain peak intensities, which were then fit a two-parameter function of the form 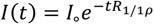 using an in-house Matlab script, with errors in relaxation rates determined by Monte Carlo analysis (69).

#### Interactions of R region and NBD1-R proteins with NBD2

Stock samples of ^15^N-labeled R region (~400 µM) and unlabeled NBD2 (200-250 µM) were prepared and used to generate NMR samples that contained 40 µM R region (or R^S903D^ or R^4P^) and 160 µM NBD2 in a total volume of 550 µL. Matched, control samples were generated by replacing NBD2 with the same volume of the NBD2 NMR buffer. Experiments were also done using 80 µM R region (or R^S903D^ or R^4P^) and 160 µM NBD2. ^1^H-^15^N HSQC spectra were recorded at 4°C. The peak intensity ratios were calculated by dividing intensities of resonances in spectra of ^15^N-labeled R region in the presence of NBD2 by the intensities of peaks in spectra of ^15^N-labeled R region alone. Errors were determined from the noise level in the HSQC spectra.

A similar approach was used to conduct NMR interaction studies between ^15^N-labeled NBD1-R (or NBD1-R^S903D^ or NBD1-R^4P^) and unlabeled NBD2. These NMR interaction experiments were also done in the presence of 2 mM MgCl_2_ and 2 mM adenylyl-imidodiphosphate (AMP-PNP) to promote dimerization of NBD1-R and NBD2. Samples for these binding experiments contained 40 µM NBD1-R proteins and either 80 µM or 160 µM NBD2. Matched controls were made for each separate interaction experiment. Note that ^1^H-^15^N HSQC experiments of NBD1-R proteins without and with NBD2 were recorded at 4°C, so that only R region resonances are observed.

### Nucleotide Binding Studies

Nucleotide binding studies were performed with two types of samples. The initial sample contained 0.75 µM NBD1 or NBD1-R proteins in the ATP-binding buffer that also contained 2 mM MgCl_2._ The titrant solutions contained high concentrations of ATP (0.5 or 2.5 mM), as well as 0.75 µM NBD1 or NBD1-R proteins and 2 mM MgCl_2_, in order to keep the protein and MgCl_2_ concentrations constant during the binding experiment. In contrast, the ATP concentration increased from 0 mM to 490 μM.

Binding of ATP to NBD1 and NBD1-R proteins was assessed by monitoring the change in the intrinsic Trp fluorescence of NBD1 (or NBD1-R). Fluorescence spectra were recorded with a Fluoromax-4 spectrofluorimeter (HORIBA) equipped with a Peltier unit for temperature control. Fluorescence emission spectra from 325-375 nm were acquired at 15 °C using an excitation wavelength of 298 nm, and excitation and emission slit widths of 2 nm 5 nm, respectively. The sample volume remained constant throughout the experiment. Note, excitation of ATP at 298 nm was negligible. The fluorescence intensity at each point in the titration was determined by averaging the fluorescence between 345-350 nm. The change in average fluorescence intensity as a function of ATP concentration was fit to the following equation, assuming a 1:1 complex between the proteins and ATP (70),

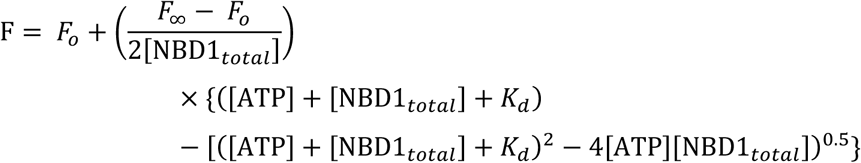

where F is fluorescence at each titration point, F_o_ is the fluorescence signal in the absence of ATP, F∞ is the fluorescence at saturating amounts of ATP, [ATP] is the total concentration of ATP at each titration point, and [NBD1] is the total NBD1 concentration in the sample. The data was fit to the above equation using a plug-in for Microsoft Excel that enables regression analysis of data with user-defined equations (71).

### ATP Hydrolysis Assays

ATPase activity was assessed using the Enzchek™ Phosphate Assay Kit (Invitrogen) (Webb 1992), in which purine nucleoside phosphorylase uses the inorganic phosphate (P*i*) produced by ATP hydrolysis to convert 2-amino-6-mercapto-7-methyl purine riboside (MESG) to ribose-1-phosphate and 2-amino-6-mercapto-7-methylpurine, which absorbs at 360 nm (52). Thus, the amount of inorganic phosphate produced from ATP hydrolysis can be quantified by measuring absorbance at 360 nm (A_360_) over time. As in previous work (35), reaction mixtures containing 10 µM of each protein (*e.g.,* 10 µM NBD1 and 10 µM NBD2), excess MESG (200 µM), 1 U purine nucleoside phosphorylase, and 5 mM MgCl_2_ in the ATPase reaction buffer were prepared. The samples were allowed to equilibrate at 21 °C for 10 minutes to ensure all samples reached the same temperature and that any contaminating inorganic phosphate was consumed. The reaction was started by the addition ATP, at a final concentration of 2 mM, and A_360_ was measured beginning at 2 min after the addition of ATP for 75 min at intervals of 1 min. Control experiments lacked one or both of the NBDs. The total volume of the samples was 1 mL. A standard curve using a phosphate standard (0-150 µM) was used to determine the amount of inorganic phosphate produced at each time point. The initial ATPase rates (nmol P*i*/min) for each sample were determined from the first 20 minutes of the ATPase time course, for which the concentration of P*i* produced *versus* time was linear.

## Acknowledgements

This research was supported by a Natural Sciences and Engineering Research Council of Canada (NSERC) Discovery Grant (RGPIN-2020-05835) to VK. During the course of the study, SCB was supported by a Queen Elizabeth II Graduate Scholarship in Science and Technology, and by a Canadian Graduate Research Scholarship-Master’s Program and an Alexander Graham Bell Canada Graduate Scholarship-Doctoral from NSERC. NMR data was collected at the University of Toronto Mississauga NMR Centre and the authors would like to thank Dmitry Pichugin and Ranjith Muhandiram for training and facilitating acquisition of triple-resonance NMR data. Activity assays were performed using instrumentation in the Core Instrumentation Facility in the Department of Chemical and Physical Sciences at the University of Toronto Mississauga. Mass spectrometry data was acquired and processed by SPARC BioCentre, SickKids Hospital.

## Data Availability

NMR chemical shifts for the R region, RS903D, and R4P are available from the BioMagResBank (https://bmrb.io/) with accession codes XXXX, YYYY, and ZZZZ, respectively.

## Author Contributions

VK conceived of the project, and SESQ, SCB, and VK designed the research. SESQ acquired and analyzed triple resonance experiments on R^4P^ and NBD1-R^4P^. SESQ acquired and analyzed ^15^N R_1_/R_1ρ_ relaxation experiments of R^4P^ and of all NBD1-R proteins in absence and presence of MgATP. SESQ conducted and analyzed all NMR binding experiments between isolated R region proteins and NBD2, and between all NBD1-R proteins and NBD2 in the presence of nucleotide. SESQ performed the fluorescence measurements used to measure ATP binding affinities of NBD1 and all NBD1-R proteins, and conducted the ATPase assays. SCB generated expression plasmids for R region and R region mutants, with assistance from AT. Samples for mass spectrometry were generated by SCB, and the data analyzed by SESQ and SCB. SCB acquired and analyzed NMR triple resonance experiments and ^15^N R1/R1ρ relaxation experiments of R and R^S903D^, and performed NMR interactions experiments between R region and NBD1 as individual polypeptides. SCB and AT performed initial NMR experiments of R region, R^S908D/T911E^, PKA-treated R region. SESQ generated all figures and wrote the paper with VK. SESQ, SCB, AT, and VK edited the paper.

