## Supplementary Figures for "Site-specific phosphorylation affects the structure and interactions of the Ycf1p R region"

**This PDF file includes:**

Supplementary Figures 1 to 22
Supplementary Tables 1 and 2

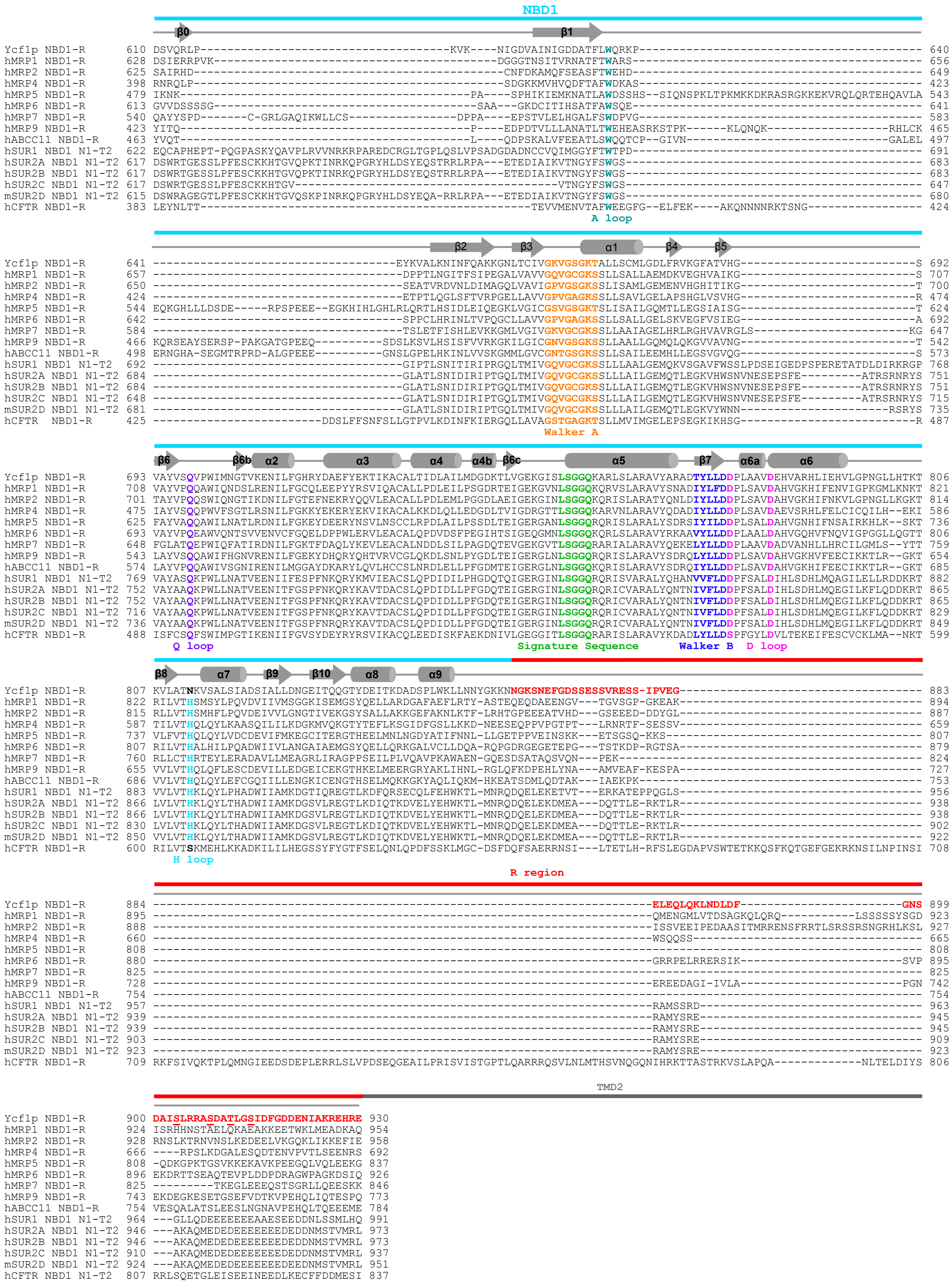
NBD1
β0
β1
A loop
β2
β3
α1
β4
β5
Walker A
β6
β6b
α2
α3
α4
α4b
β6c
α5
β7
α6a
α6
Q loop
Signature Sequence
Walker B
D loop
β8
α7
β9
β10
α8
α9
H loop
R region
TMD2
Ycf1p NBD1-R
hMRP1 NBD1-R
hMRP2 NBD1-R
hMRP4 NBD1-R
hMRP5 NBD1-R
hMRP6 NBD1-R
hMRP7 NBD1-R
hMRP9 NBD1-R
hABCC11 NBD1-R
hSUR1 NBD1 N1-T2
hSUR2A NBD1 N1-T2
hSUR2B NBD1 N1-T2
hSUR2C NBD1 N1-T2
mSUR2D NBD1 N1-T2
hCFTR NBD1-R

19

**Supplementary Fig. 1. Structure-based sequence alignment of ABCC proteins, focussing on NBD1-R.** Structural information of Ycf1p (PDB code 7MPE (1)), rat MRP2 (PDB code 8RQ3 (2)), human MRP3 (PDB code 8HVH (3)), human CFTR (PDB code 6O1V (4)), and human SUR1 (PDB code 6JB1 (5)) was used in the alignment. The secondary structural elements of Ycf1p NBD1-R are displayed and conserved residues in NBD1 are highlighted in various colours. The R region in SUR proteins is denoted as the NBD1-TMD2 (N1-T2) linker (6), hence the notation of NBD1 N1-T2 for SUR1 and isoforms of SUR2. Residues corresponding to NBD1 and R region are indicated by a light blue and red line, respectively, above the schematic representation of the 2º structure elements. The dark grey line indicates the position of TMD2 in the primary structure of ABCC proteins, even though those residues are not shown. Note that β0-strand is seen in Ycf1p structures (1, 7), but necessarily in other ABC protein structures.

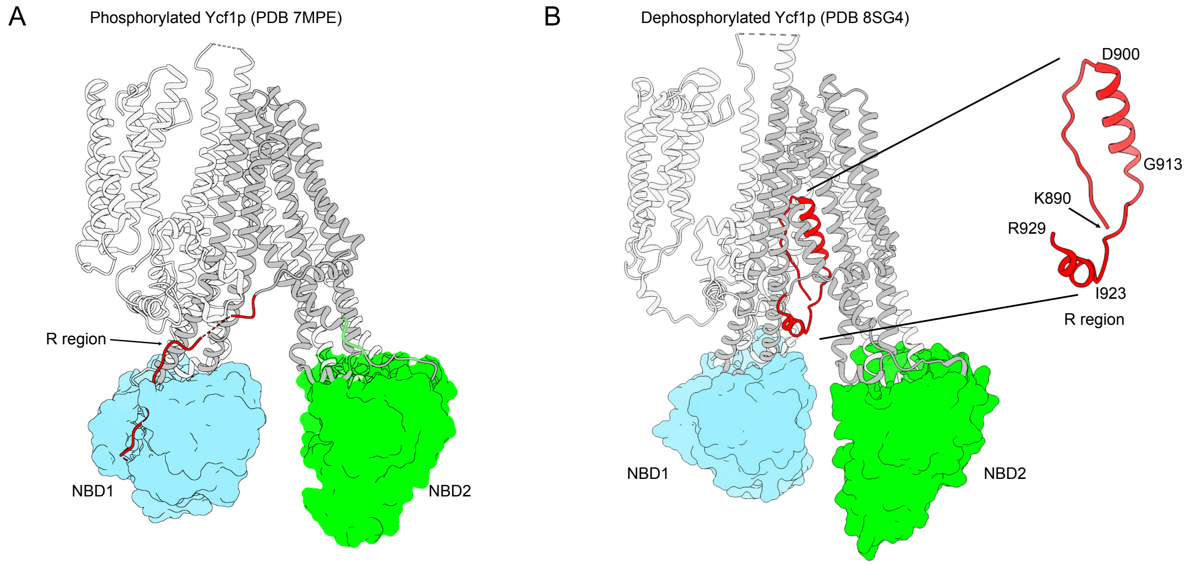


**Supplementary Fig. 2. Cyro-EM structures of Ycf1p show that the conformation and interactions of the R region are sensitive to its phosphorylation state.** (A) The phosphorylated Ycf1p structures (*e.g.,* PDB 7MPE (1)) show an extended conformation of R region that is bound to the peripheral side of NBD1. In this particular structure, R region residues S863-V874 are engaged in NBD1 binding. (B) In the dephosphorylated Ycf1p structure (PDB 8SG4 (8)), R region residues K890-R929 adopt a helix-strand hairpin motif that is lodged in the substrate binding cavity. In both panels A and B, NBD1 and NBD2 are shown in a surface representation coloured blue and green, respectively. Transmembrane domains are shown as schematic ribbon diagrams with TMD0 and TMD1 in light gray, and TMD2 in dark gray. Figures were generated with the structure analysis and visualization program ChimeraX (9).

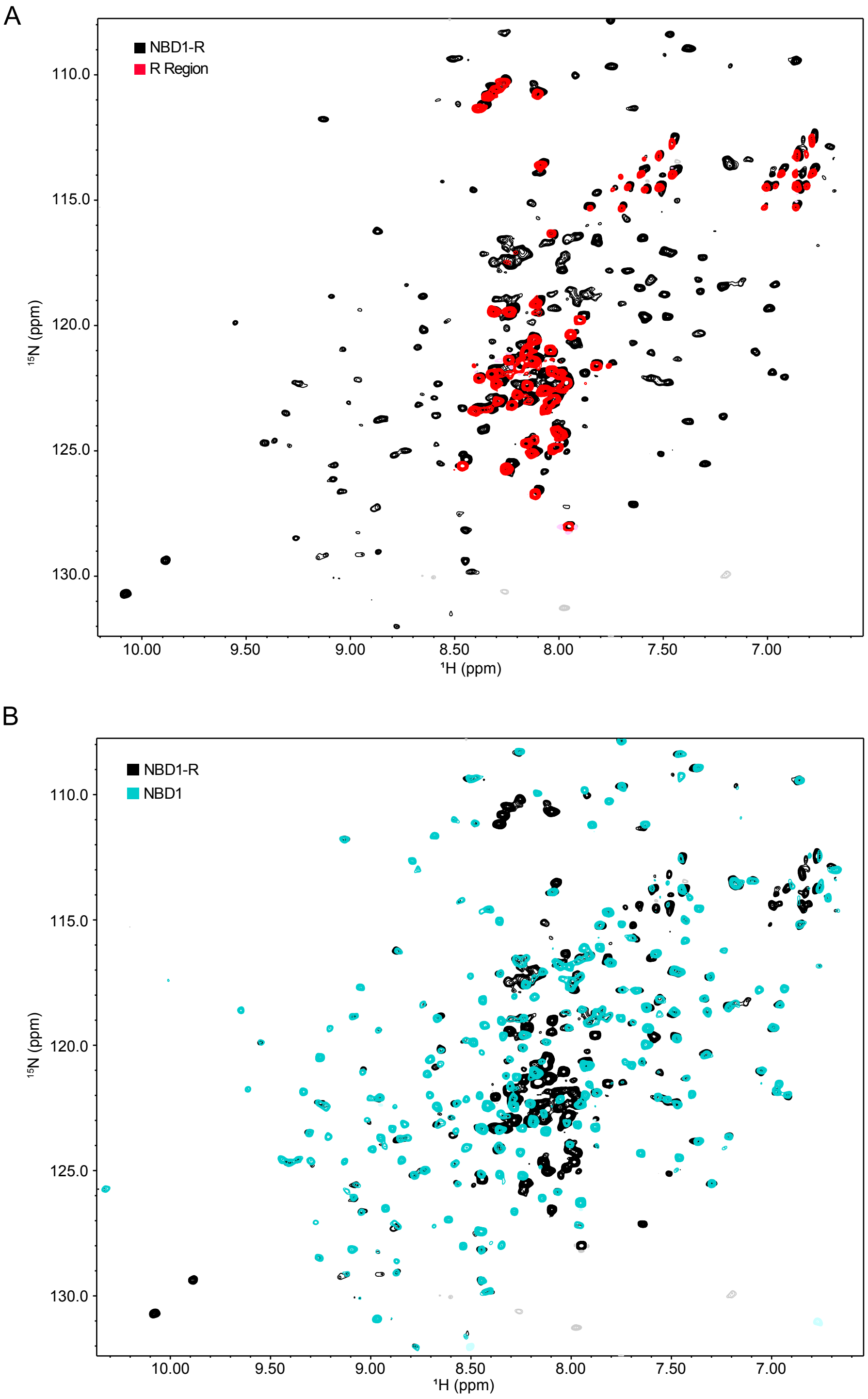


**Supplementary Fig. 3. Comparison of R region, NBD1, and NBD1-R spectra show that R region and NBD1 can be studied as isolated proteins or as part of NBD1-R.** (A) Comparison of $^{1}H$-$^{15}N$ TROSY-HSQC spectra of NBD1-R (200 μM) at 30 ºC and $^{1}H$-$^{15}N$ HSQC spectra of R region (94 μM) at 5 ºC. The spectrum of NBD1-R is shown in black and in the background, while the spectrum of R region is shown in red and in the foreground. (B) Comparison of $^{1}H$-$^{15}N$ TROSY-HSQC spectra of NBD1-R (200 μM) and NBD1 (300 μM) recorded at 30 ºC. The spectrum of NBD1-R is shown in black and in the background, while NBD1 is shown in aqua blue in the foreground. The NBD1 and NBD1-R samples each contain 2 mM MgATP. Note that signals in spectra of isolated R region overlay with R region signals in spectra of NBD1-R lacking MgATP (data not shown).

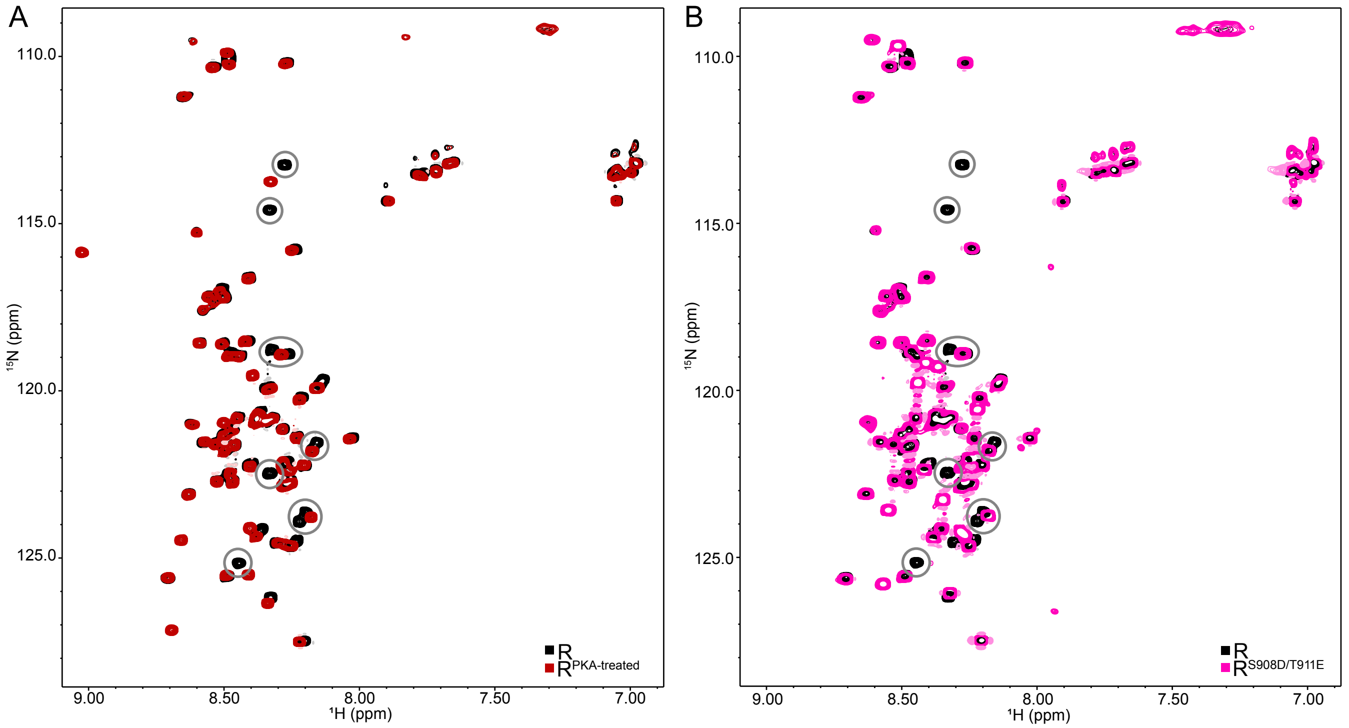


**Supplementary Fig. 4**. **PKA-dependent phosphorylation and phosphomimic mutation of S908 and T911 cause similar changes in R region.** Overlayed $^{1}$H-$^{15}$N HSQC spectra of non-phosphorylated R region (black) and (A) PKA-treated R region (dark red) or (B) the R$^{S908D/T911E}$ mutant (magenta) recorded at 4 ºC. In both panels, the spectrum of the R region is in the background. The spectra of PKA-treated R region and the R$^{S908D/T911E}$ mutant are in the foreground in panels A and B, respectively. R region resonances that change with PKA phosphorylation of S908 and T911 or with introduction of the S908D and T911E phosphomimetic mutations are circled.

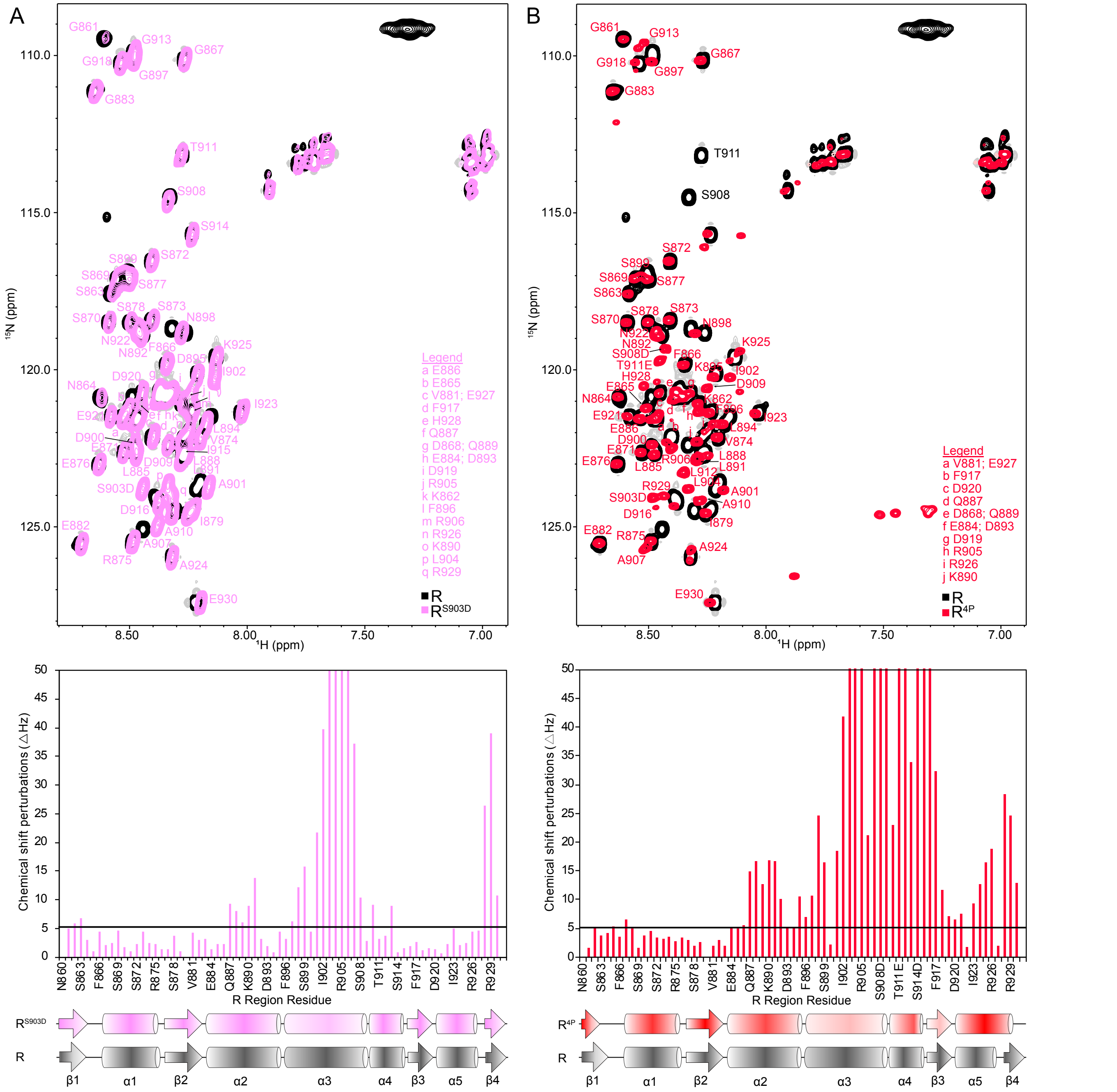


**Supplementary Fig. 5. Phosphorylation and phosphomimetic mutations cause large-scale NMR chemical shift changes in R region resonances.** Overlayed $^{1}$H-$^{15}$N HSQC spectra of non-phosphorylated R region (black) and (A, *top*) the R$^{S903D}$ mutant (pink) or (B, *top*) the R$^{4P}$ mutant(red) recorded at 4 ºC. In both panels, the spectrum of the R region is in black and in the background. The spectra of R$^{S903D}$ and R$^{4P}$ are in pink and red, respectively, and in the foreground in panels A and B. Resonances in the R$^{S903D}$ spectrum (A, top) and in the R$^{4P}$ spectrum (B, top) are labeled with their assignments in pink and red, respectively. The S908 and T911 resonances in the non-phosphorylated R region spectrum in panel B are labeled with their assignments in black. Chemical shift differences of $^{15}$N-$^{1}$H$^{N}$ resonances between spectra of non-phosphorylated R region and R$^{S903D}$ (A, bottom) and between spectra of non-phosphorylated R region and R$^{4P}$ (B, bottom) are plotted as a function of R region residue. The black line indicates the chemical shift perturbation threshold, which is defined as the average of the chemical shift differences between the two spectra plus one standard deviation. The schematic representation of the non-phosphorylated R region SSP data from Fig. 2 is shown below both plots. Also shown, below the corresponding plot, are the schematic representations of the $R^{S903D}$ or $R^{4P}$ SSP data.

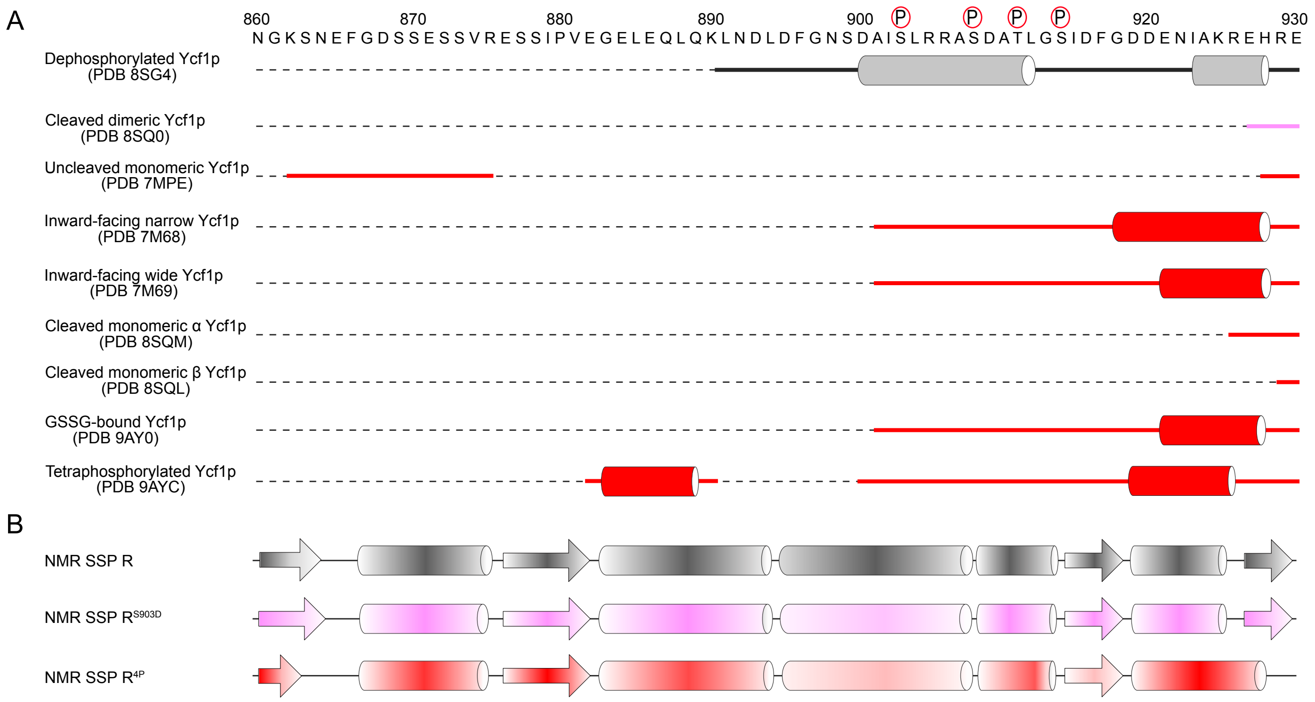


**Supplementary Fig. 6. Variability of R region structure in cryo-EM structures of Ycf1p and comparison to residual secondary structures determined by NMR spectroscopy.** (A) Schematic representation of secondary structures observed for R region in different Ycf1p cryo-EM structures and coloured by phosphorylation state (grey, dephosphorylated; pink, mono-phosphorylated at S903; red, multi-site phosphorylation) (*top*). The resolved portions of the R region are depicted with solid lines representing extended R region segments that lack regular secondary structure and with cylinders representing α-helices. (B) Schematic representation of residual secondary structure of isolated R region proteins from the SSP analysis, from Fig. 2, with cylinders representing α-helices an arrows representing β-strands. The individual secondary structures are coloured as in panel A and shaded according to the SSP value.

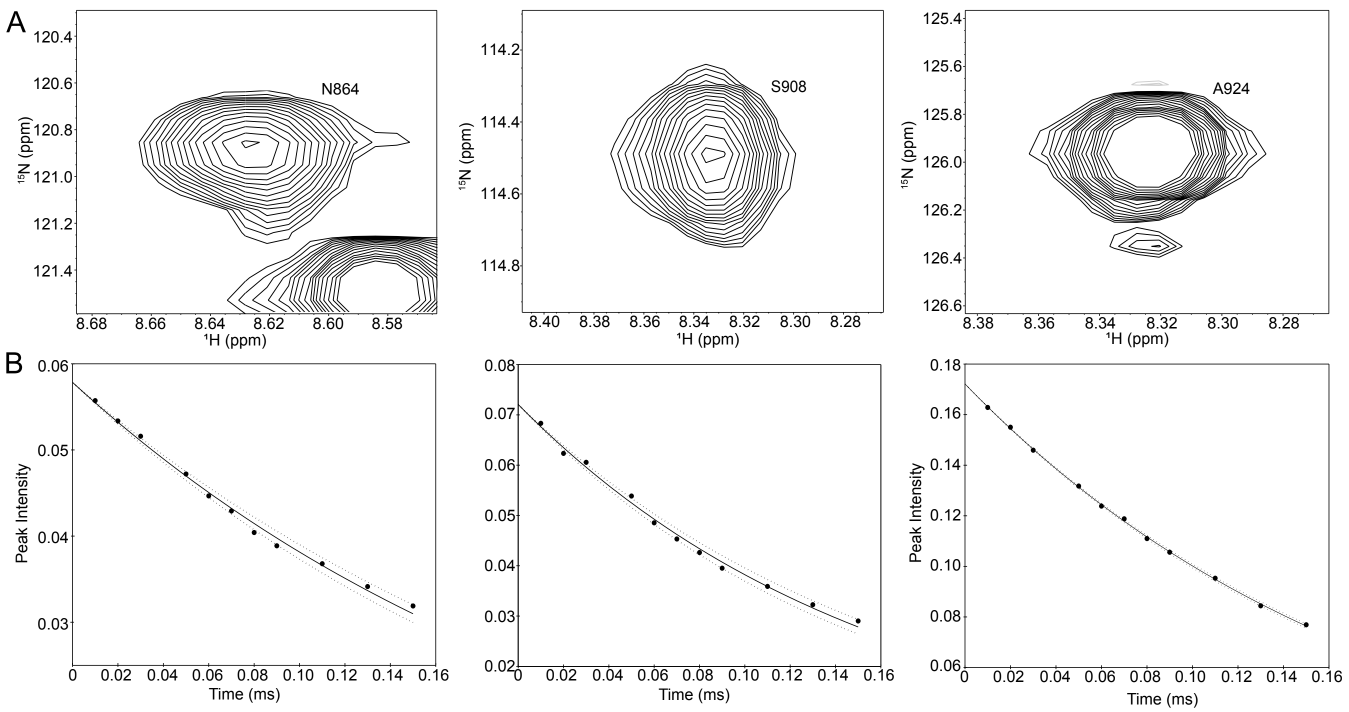


**Supplementary Fig. 7**. **R region $^{15}$N $R_{1\rho}$ relaxation data.** (A) Selected $^{1}$H-$^{15}$N resonances corresponding to R region residues N864, S908, and A924 from $^{15}$N-$^{1}$H correlation spectrum, recorded with a delay of 10 ms, used to the determine the $^{15}$N $R_{1\rho}$ rates. (B) Corresponding decay curves for the resonances displayed in panel A, with peak intensities at each delay time indicated by solid circles. The solid lines indicate the fits of the data to the equation $I(t) = I_{\circ}e^{-tR_{1\rho}}$, whereas the dotted lines indicate the upper and lower bounds as determined by Monte Carlo analysis.

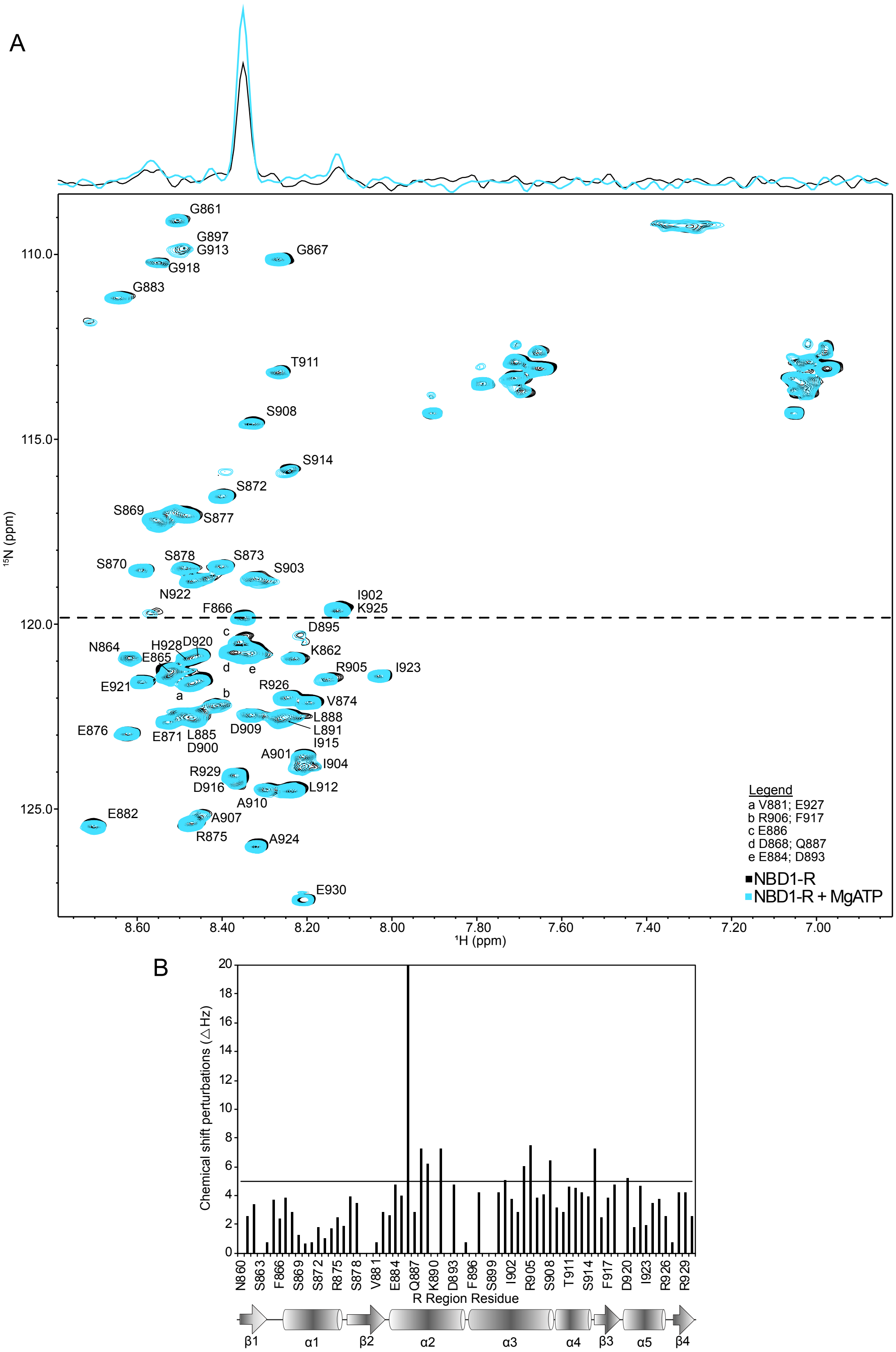


**Supplementary Fig. 8**. **Comparison of $^{1}H$-$^{15}N$ HSQC spectra of NBD1-R (200 µM) with and without 2 mM MgATP at 4 ºC.** (A) The apo NBD1-R spectrum is in black in the background and the MgATP-bound NBD1-R spectrum is in sky blue and in the foreground. The $^{1}H$ trace through the approximate centre of the $^{15}N$-$^{1}H$ resonance for F866, as depicted by the dashed line, is shown on top of the spectrum. (B) Chemical shift differences of $^{15}N$-$^{1}H^{N}$ resonances between spectra of NBD1-R in the absence and presence of 2 mM MgATP are plotted as a function of R region residue. The black line indicates the chemical shift perturbation threshold, which is defined as the average of the chemical shift differences between the two spectra plus one standard deviation. The schematic representation of non-phosphorylated R region SSP data, from Fig. 2, is shown below the plot. The data shown are representation of two experiments on separate preparations of NBD1-R.

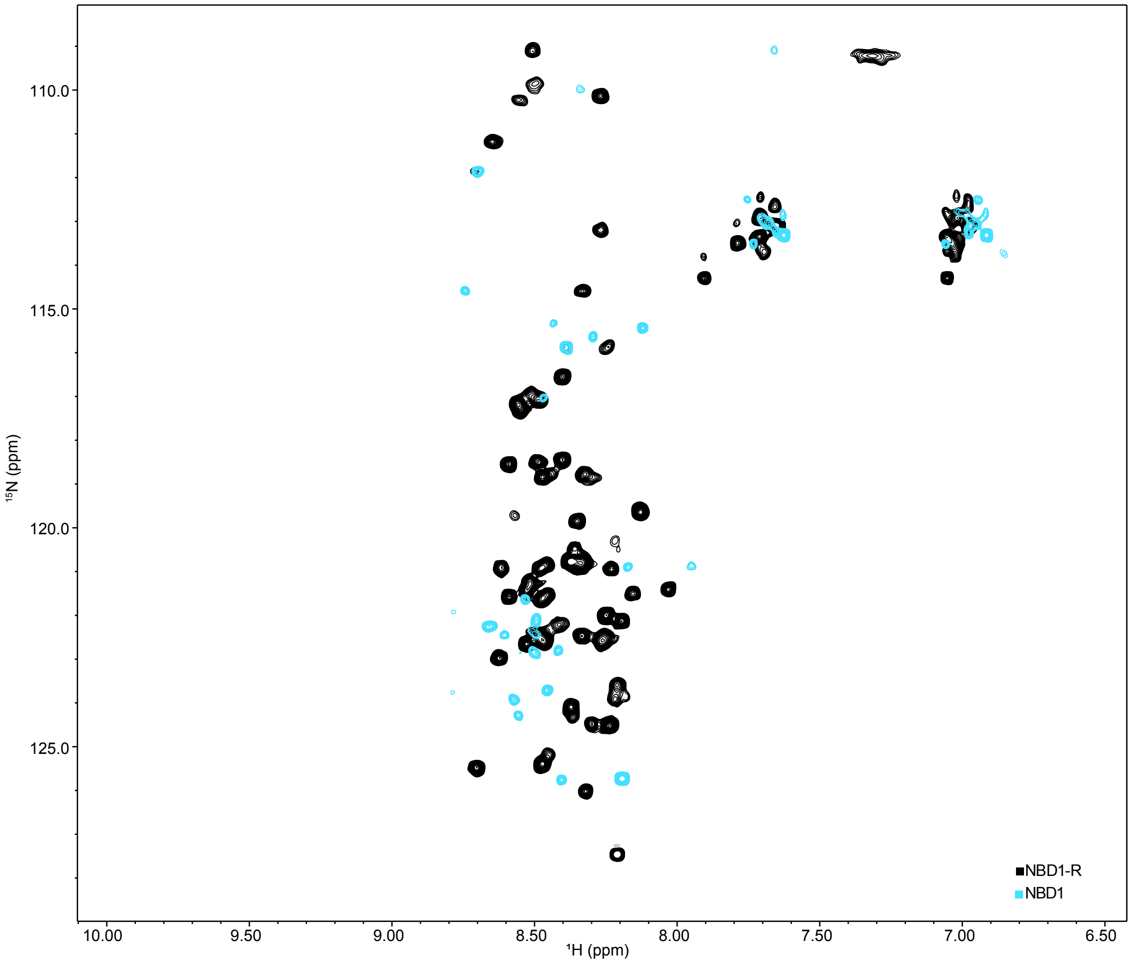


**Supplementary Fig. 9. NBD1 signals are silenced in $^{1}$H-$^{15}$N HSQC spectra at low temperatures**. Overlay of $^{1}$H-$^{15}$N HSQC spectra of 200 μM NBD1-R (in black and in the background) and 200 μM NBD1 (in cyan and in the foreground). Both spectra are collected at 4 ºC and with the same spectral parameters. Most NBD1 resonances are not visible in $^{1}$H-$^{15}$N HSQC spectra.

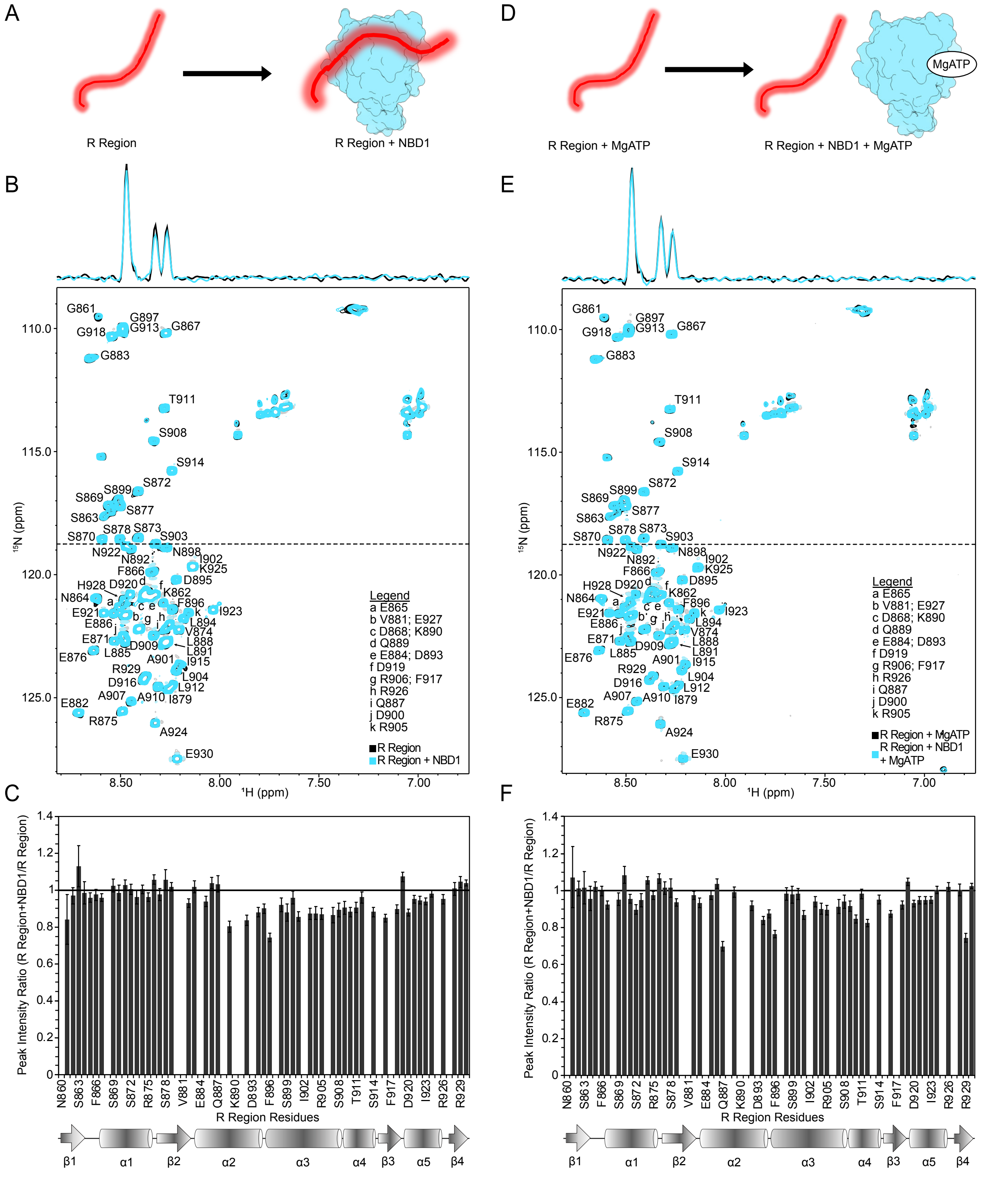


**Supplementary Fig. 10. Interactions of R region with NBD1 as separate polypeptides in the absence and presence of 2 mM MgATP.** (A) Schematic representation of the R region and NBD1 samples used in the NMR interaction experiments. Experiments are performed such that R region was $^{15}$N-labelled and NBD1 was unlabelled. (B) Overlay of the $^{1}$H-$^{15}$N HSQC spectra of R region (18 μM), recorded at 4 ºC, in the absence (in black and in the background) and presence (in blue and in the foreground) of 180 μM NBD1. The $^{1}$H trace through the approximate centre of the $^{15}$N-$^{1}$H resonance for

S903, as depicted by the dashed line, is shown on top of the spectrum. (C) Plot of the peak intensity ratios for R region with and without NBD1 as a function of R region residues. Peak intensity ratios above 1 indicate increased intensities of the corresponding resonances upon addition of NBD1, while ratios below 1 indicate decreased NMR intensities. Schematic representation of the secondary structure of R region from Fig. 2 is shown below the plot. (D) Schematic representation of the R region and NBD1 samples used in the NMR interaction experiments in the presence of 2 mM MgATP. Experiments are performed such that R region was $^{15}$N-labelled and NBD1 was unlabelled. (E) Overlay of the $^{1}$H-$^{15}$N HSQC spectra of R region (18 µM), recorded at 4 ºC, in the absence (in black and in the background) and presence (in blue and in the foreground) of 180 µM NBD1. The $^{1}$H trace through the approximate centre of the $^{15}$N-$^{1}$H resonance for S903, as depicted by the dashed line, is shown on top of the spectrum. All samples contained 2 mM MgATP. (F) Plot of the peak intensity ratios for R region with and without NBD1 as a function of R region residues. Peak intensity ratios above 1 indicate increased intensities of the corresponding resonances upon addition of NBD1, while ratios below 1 indicate decreased NMR intensities. Schematic representation of the secondary structure of R region from Fig. 2 is shown below the plot. The data shown are representation of two experiments on separate preparations of NBD1-R$^{S903D}$ and NBD1-R$^{4P}$.

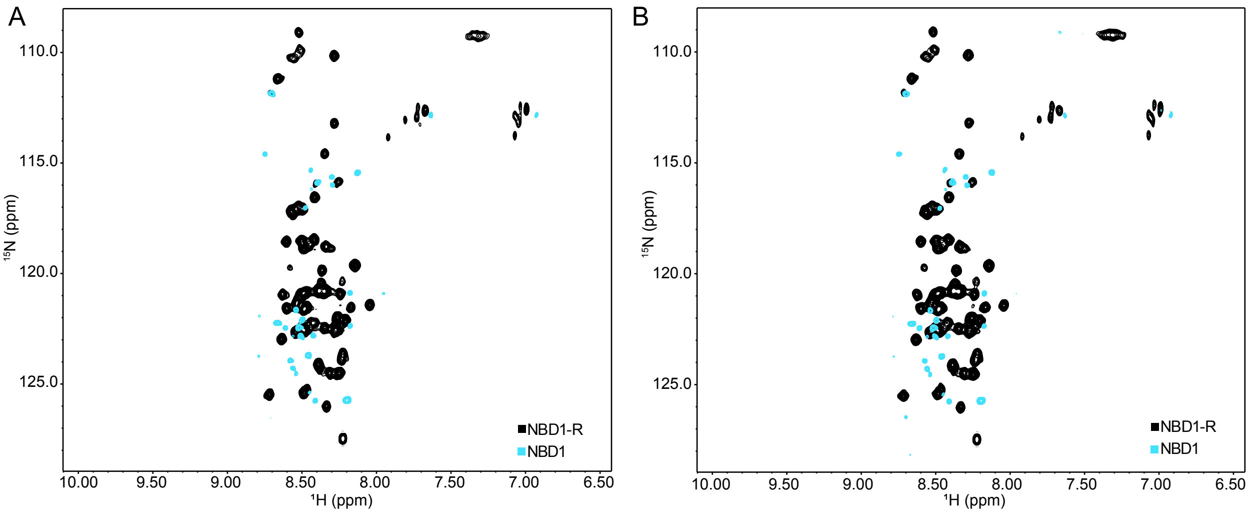


**Supplementary Fig. 11. NBD1 signals are silenced in non-TROSY-based experiments for measuring $^{15}N$ $R_{1\rho}$ and $^{15}N$ $R_1$ rates recorded low temperatures**. Overlay of the non-TROSY $^{15}N$-$^{1}H$ correlation spectra for obtaining (A) $^{15}N$ $R_{1\rho}$ and (B) $^{15}N$ $R_1$ of 200 μM NBD1-R (in black and in the background) and 200 μM NBD1 (in cyan and in the foreground). The $R_{1\rho}$ and $R_1$experiments were recorded at 4 ºC. The $^{15}N$-$^{1}H$ correlation spectra shown were obtained with delay times of 10 ms for the $R_{1\rho}$ experiment and 10.1 ms for the $R_1$, the shortest delays used for obtaining $^{15}N$ $R_{1\rho}$ and $R_1$ rates for the R region residues in NBD1-R and in the isolated R region. Given the low intensity of the NBD1 peaks in the spectra shown above, it is likely that NBD1 signals would be absent in spectra collected at the longer delay times used in the $^{15}N$ $R_{1\rho}$ and $R_1$ experiments.

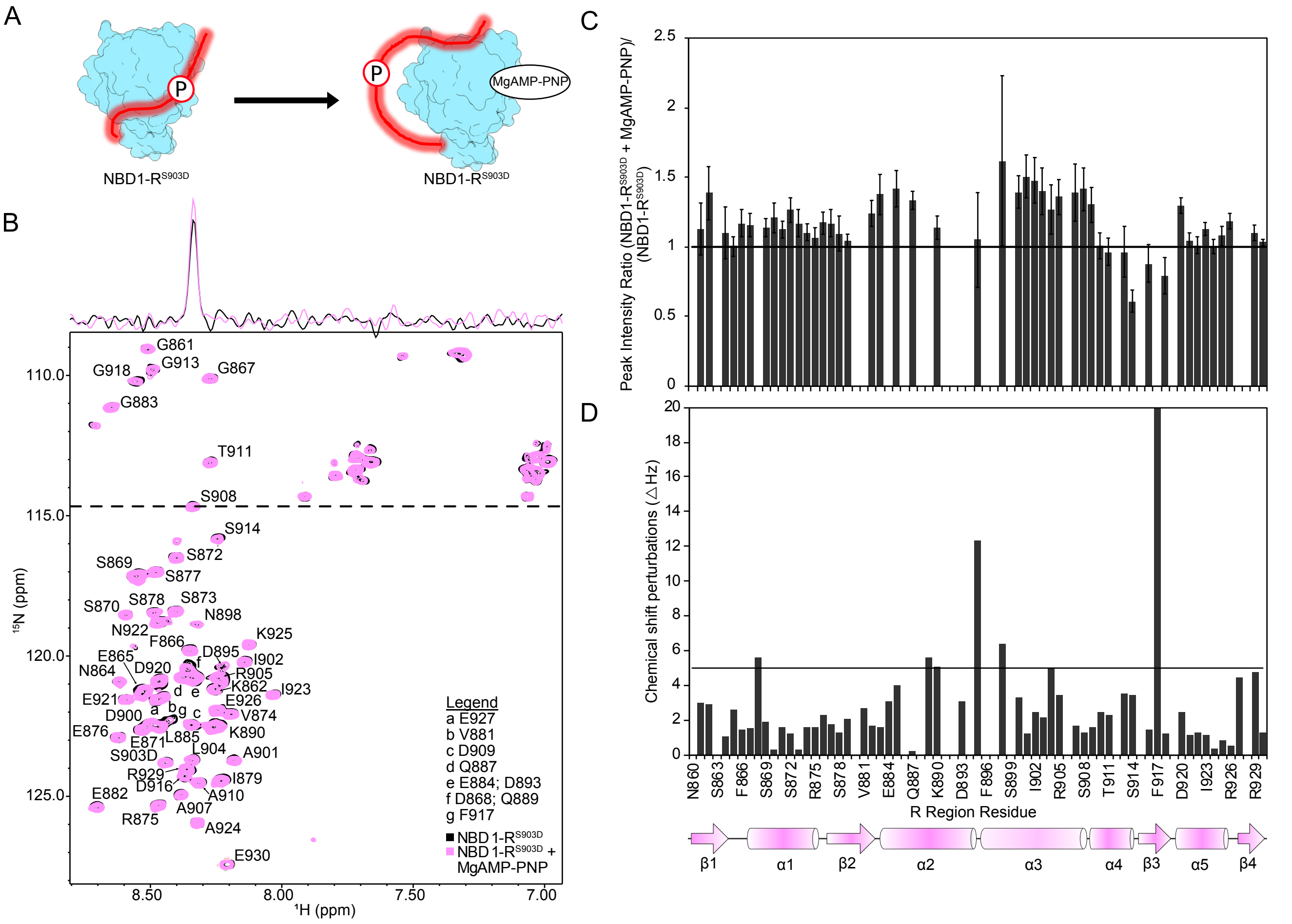


**Supplementary Fig. 12. Interactions of $R^{S903D}$ with NBD1 change when NBD1 binds nucleotide.** (A) Schematic representation of the NBD1-$R^{S903D}$ samples used in the NMR interaction experiments. Experiments are performed such that NBD1 resonances are silenced and only $R^{S903D}$ peaks are visible, represented by the red glow around $R^{S903D}$. (B) Overlay of the $^{1}$H-$^{15}$N HSQC spectra of NBD1-$R^{S903D}$ (40 μM), recorded at 4 ºC, in the absence (in black and in the background) and presence (in pink and in the foreground) of 2 mM MgATP. The $^{1}$H trace through the approximate centre of the $^{15}$N-$^{1}$H$^{N}$ resonance for S908, as depicted by the dashed line, is shown on top of the spectrum. (C) Plot of the peak intensity ratios for NBD1-$R^{S903D}$ with and without MgATP as a function of R region residues. Peak intensity ratios above 1 indicate increased intensities of the corresponding resonances upon addition of MgATP, while ratios below 1 indicate decreased NMR intensities. (D) Chemical shift differences of $^{15}$N-$^{1}$H$^{N}$ resonances between spectra of NBD1-$R^{S903D}$ in the absence and presence of 2 mM MgATP are plotted as a function of R region residue. The black line indicates the chemical shift perturbation threshold, which is defined as the average of the chemical shift differences between the two spectra plus one standard deviation. Schematic representation of the secondary structure of $R^{S903D}$ from Fig. 2 is shown below the plot. The data shown are representation of two experiments on separate preparations of NBD1-$R^{S903D}$.

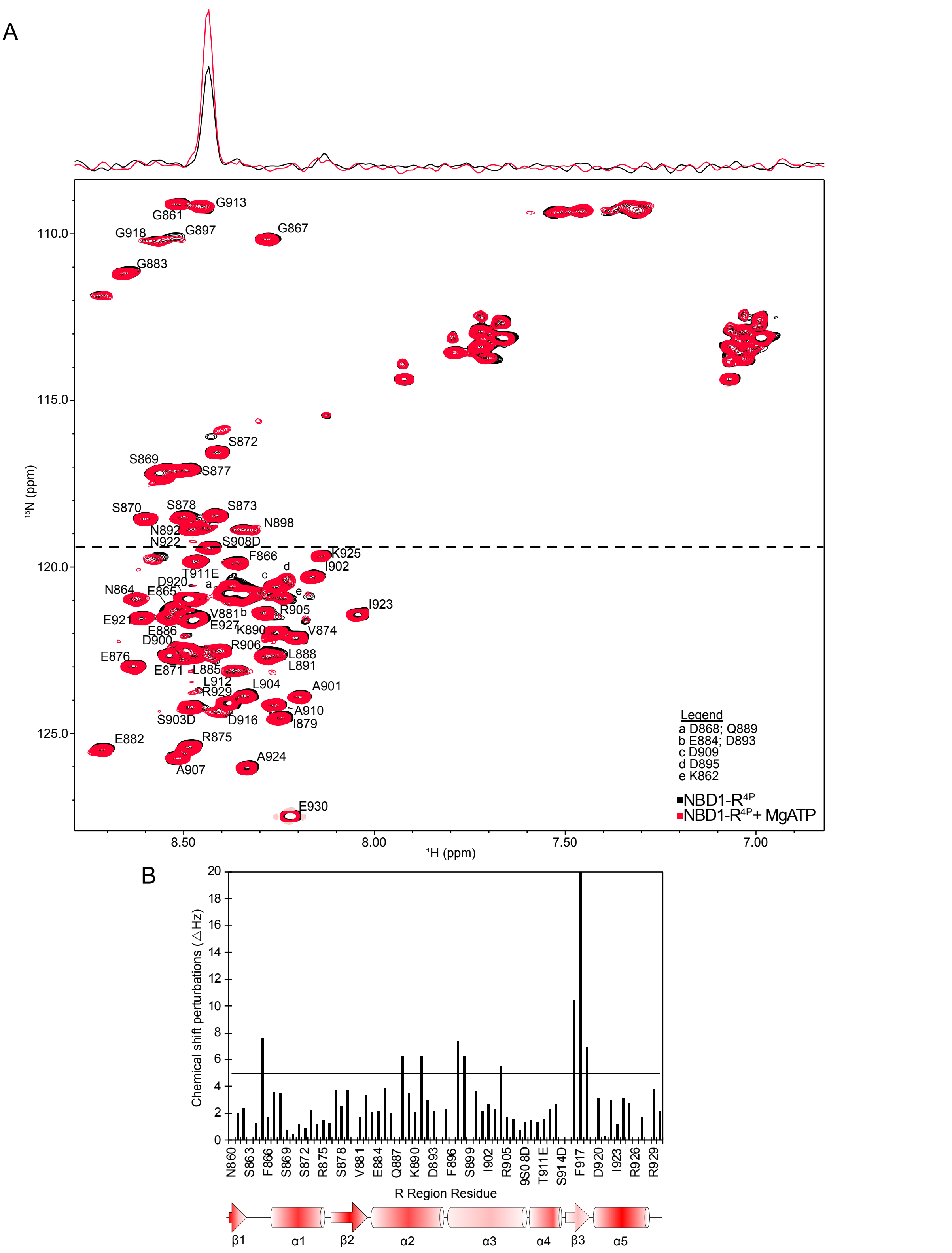


**Supplementary Fig. 13**. Comparison of $^1$H-$^{15}$N HSQC spectra of NBD1-R$^{4P}$ (200 µM) with and without 2 mM MgATP at 4 ºC. (A) The spectrum of NBD1-R$^{4P}$ without MgATP is in black and in the background, while the spectrum of NBD1-R$^{4P}$ with MgATP is in red and in the foreground. The $^1$H trace through the approximate centre of the $^{15}$N-$^1$H resonance for the phosphomimetic S908D mutated residue, as depicted by the dashed line, is shown on top of the spectrum. (B) Chemical shift differences of $^{15}$N-$^1$H$^N$ resonances between spectra of NBD1-R$^{4P}$ in the absence and presence of 2 mM MgATP are plotted as a function of R region residue. The black line indicates the chemical shift perturbation threshold, which is defined as the average of the chemical shift differences between the two spectra plus one standard deviation. Schematic representation of the secondary structure of $R^{4P}$ from Fig. 2 is shown below the plot. The data shown are representation of two experiments on separate preparations of NBD1-$R^{4P}$.

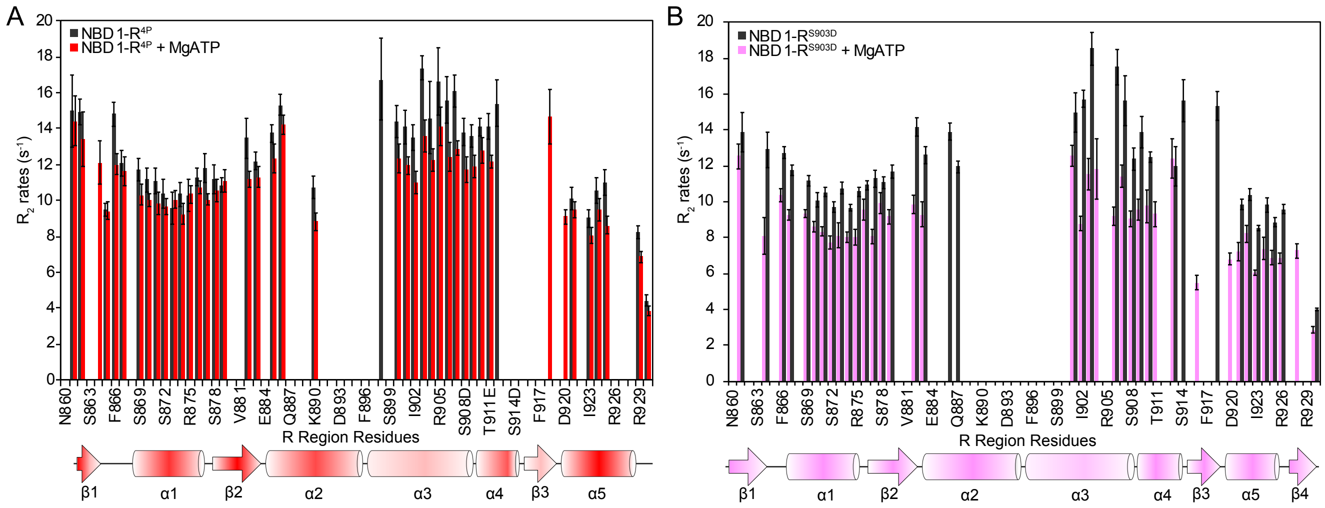


**Supplementary Fig. 14. Dynamics of R residues in NBD1-R$^{S903D}$ and NBD1-R$^{4P}$ changes in the presence of nucleotide.** (A) $^{15}$N $R_2$ relaxation rates for 200 μM NBD1-R$^{4P}$ in the absence (dark gray) and presence (pink) of 2 mM MgATP. Schematic representation of R$^{4P}$ SSP values (Fig. 2) is shown below the plot. For NBD1-R$^{S903D}$ and NBD1-R$^{4P}$, overlapped resonances or resonances with low signal:noise ratios were omitted from the analysis. (B) $^{15}$N $R_2$ relaxation rates for 200 μM NBD1-R$^{S903D}$ in the absence (dark gray) and presence (pink) of 2 mM MgATP. Schematic representation of R$^{S903D}$ SSP values (Fig. 2) is shown below the plot.

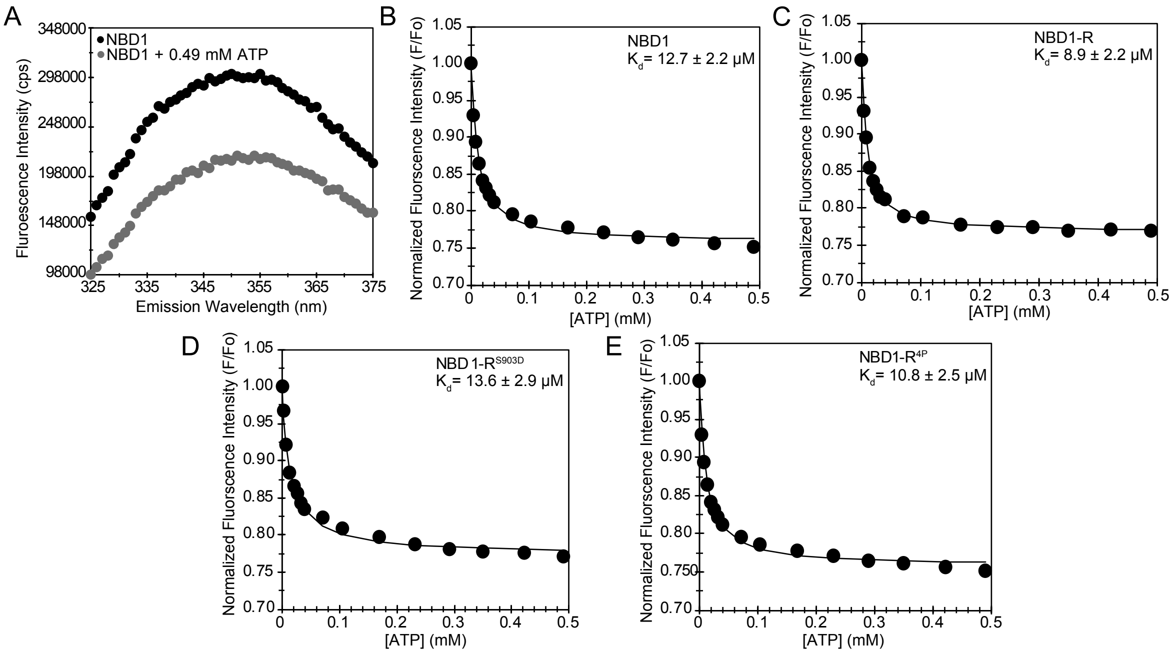


**Supplementary Fig. 15. Intrinsic Trp fluorescence of NBD1 and NBD1-R proteins change upon nucleotide binding and can be used to determine the associated binding affinities.** (A) Representative Trp fluorescence emission spectra of NBD1 with 0 mM (black) and 0.49 mM MgATP (grey). Each emission spectrum is corrected for the fluorescence of the buffer. (B-E) Representative fluorescence binding curves for the interactions of MgATP with (B) NBD1, (C) NBD1-R, (D) NBD1-$R^{S903D}$, and (E) NBD1-$R^{4P}$. The average $K_d$ values for the NBD1/MgATP and NBD1-R/MgATP interactions were determined from 6 trials using purified proteins from each of two separate preparations. The average $K_d$ value for the NBD1-$R^{S903D}$/MgATP interaction was determined from 9 trials using purified NBD1-$R^{S903D}$ from three separate preparations. The average $K_d$ value was determined from 11 trials using purified NBD1-$R^{4P}$ from four separate preparations.

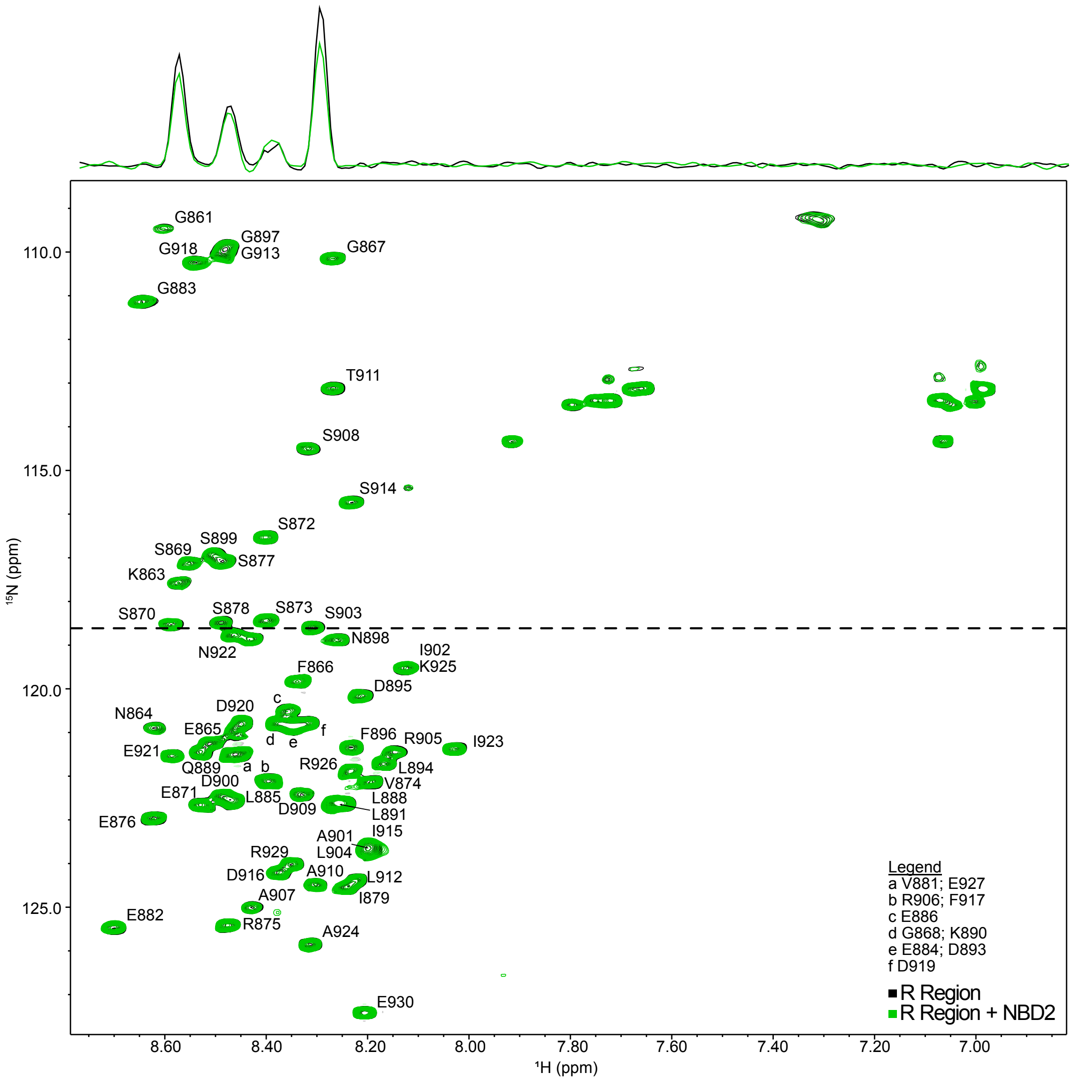


**Supplementary Fig. 16.** Comparison of $^{1}$H-$^{15}$N HSQC spectra of R region (40 μM) in the presence and absence of NBD2 (160 μM) at 4 ºC. The spectrum of R region in absence of NBD2 is in black and in the background, whereas that of R region with NBD2 is in green and in the foreground. The $^{1}$H trace through the approximate centre of the $^{15}$N-$^{1}$H resonance for S903, as depicted by the dashed line, is shown on top of the spectrum. The data shown are representation of two experiments on separate preparations of R region and NBD2.

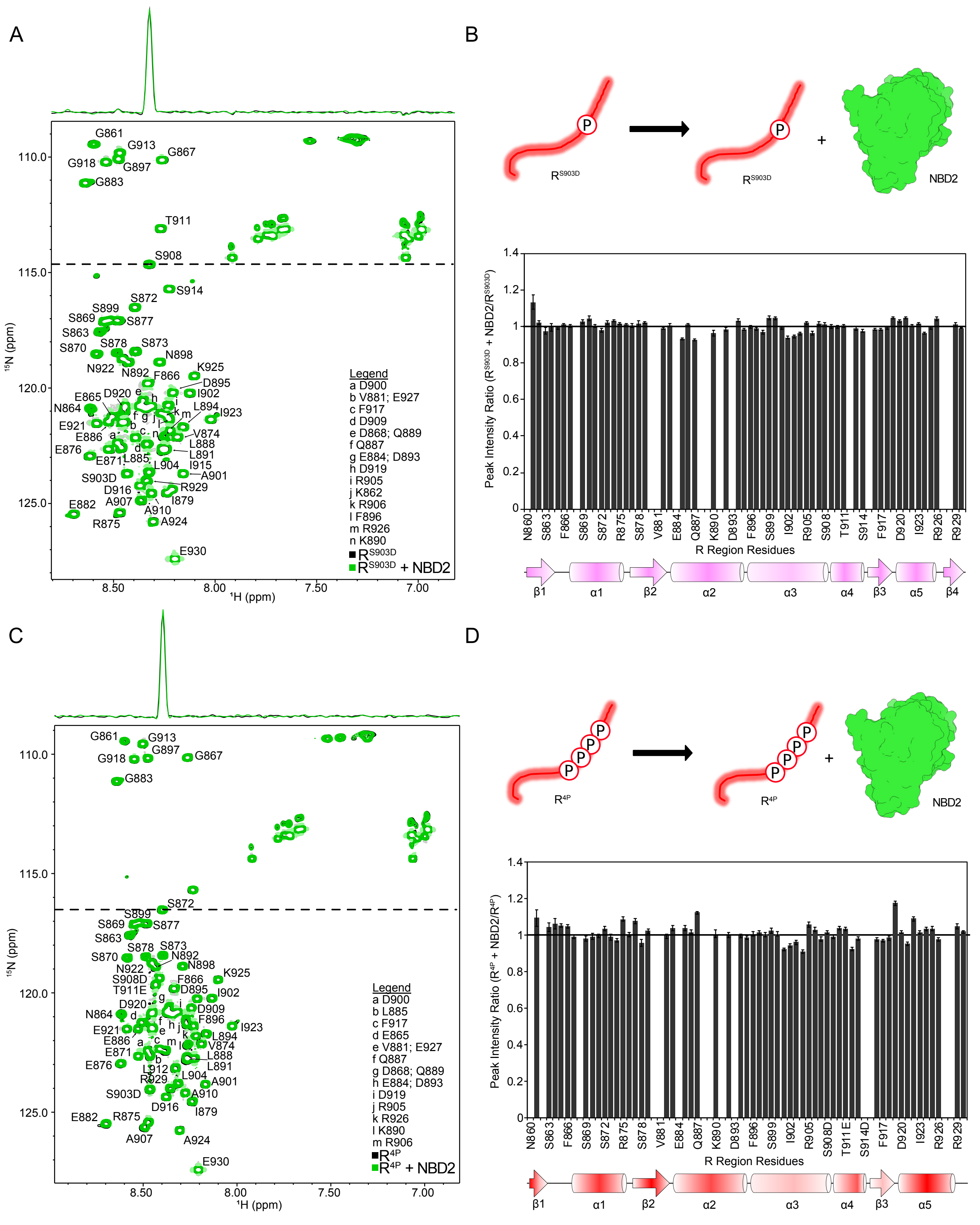


**Supplementary Fig. 17. The $R^{S903D}$ and $R^{4P}$ proteins do not interact with NBD2.** (A) Overlay of the $^{1}H$-$^{15}N$ HSQC spectra, recorded at 4 ºC, of 40 μM $R^{S903D}$ in the absence (in black and in the background) and presence (in green and in the foreground) of 160 μM NBD2. The $^{1}H$ trace through the approximate centre of the $^{15}$N-$^{1}$H resonance for S908, as depicted by the dashed line, is shown on top of the spectrum. (B) Schematic representation of the samples used in the NMR interaction experiments in panel A (*top*). Plot of the peak intensity ratios for $R^{S903D}$ with and without NBD2 as a function of R region residues (*bottom*). The fact that most peak intensity ratios are ~1 indicates that $R^{S903D}$ does not bind NBD2 Schematic representation of the secondary structure of $R^{S903D}$ from Fig. 2 is shown below the plot. (C) Overlap of the $^{1}$H-$^{15}$N HSQC spectra, recorded at 4 ºC, of 40 μM $R^{4P}$ in the absence (in black and in the background) and presence of 160 μM NBD2 (in green and in the foreground). The $^{1}$H trace through the approximate centre of the $^{15}$N-$^{1}$H resonance for S872, as depicted by the dashed line, is shown on top of the spectrum. (D) Schematic representation of the protein samples used in the NMR interaction experiments in panel C (*top*). Plot of the peak intensity ratios for $R^{4P}$ with and without NBD2 as a function of R region residues (*bottom*), demonstrating that $R^{4P}$ does not bind NBD2. Schematic representation of the secondary structure of $R^{4P}$ from Fig. 2 is shown below the plot. Data shown in panels A-D are representative of experiments repeated from two separate preparations of all proteins.

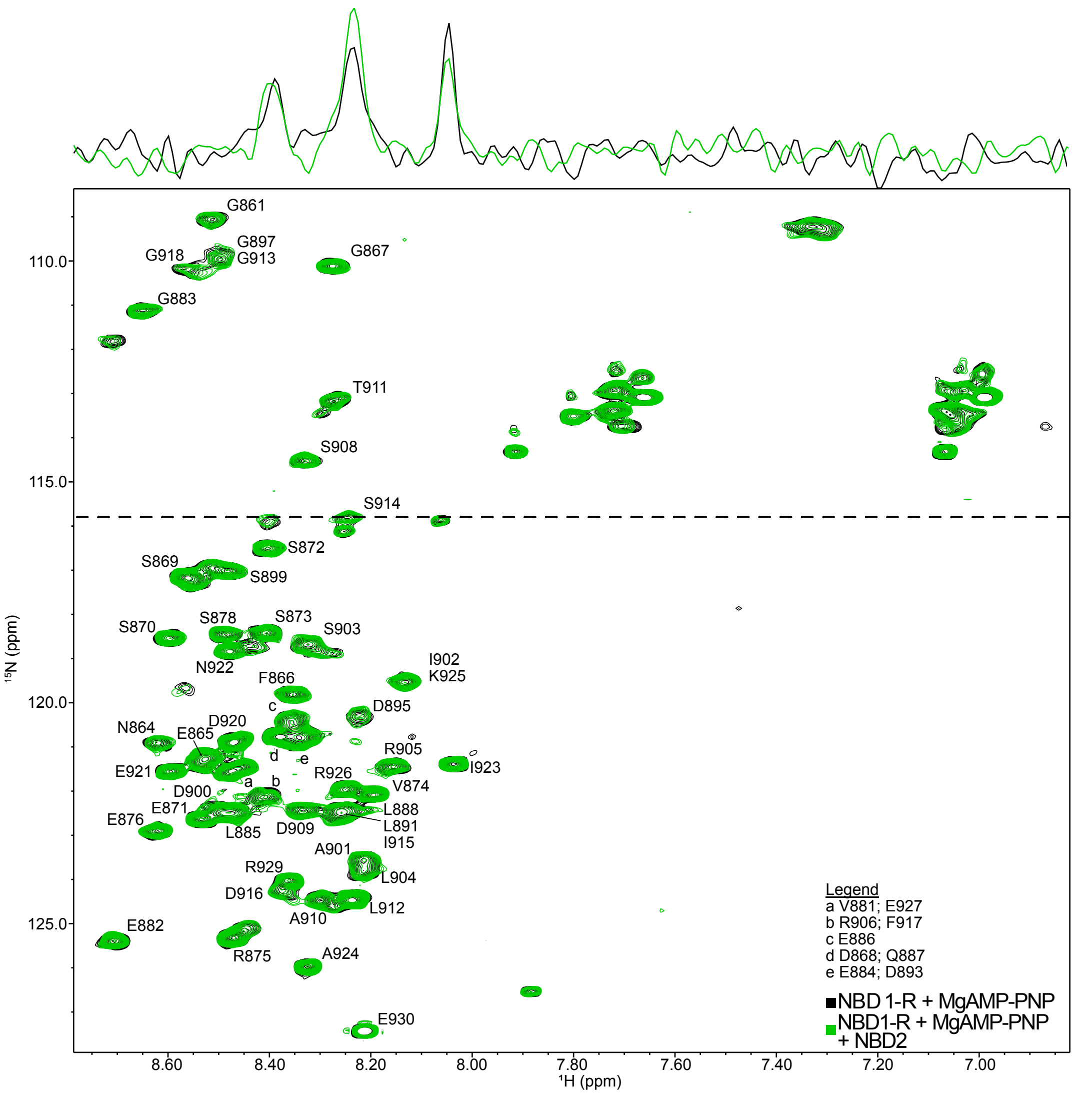


**Supplementary Fig. 18. R interactions when NBD1-R dimerizes with NBD2 in the presence of nucleotide.** Overlay of $^{1}H$-$^{15}N$ HSQC spectra, recorded at 4 ºC of 40 µM NBD1-R in the absence (black, background) and presence of 160 µM NBD2) with 2 mM MgAMP-PNP (green, foreground). The $^{1}H$ trace through the approximate centre of the $^{15}N$-$^{1}H$ resonance for S914, as depicted by the dashed line, is shown on top of the spectrum. The data shown are representation of two experiments on separate preparations of NBD1-R and NBD2.

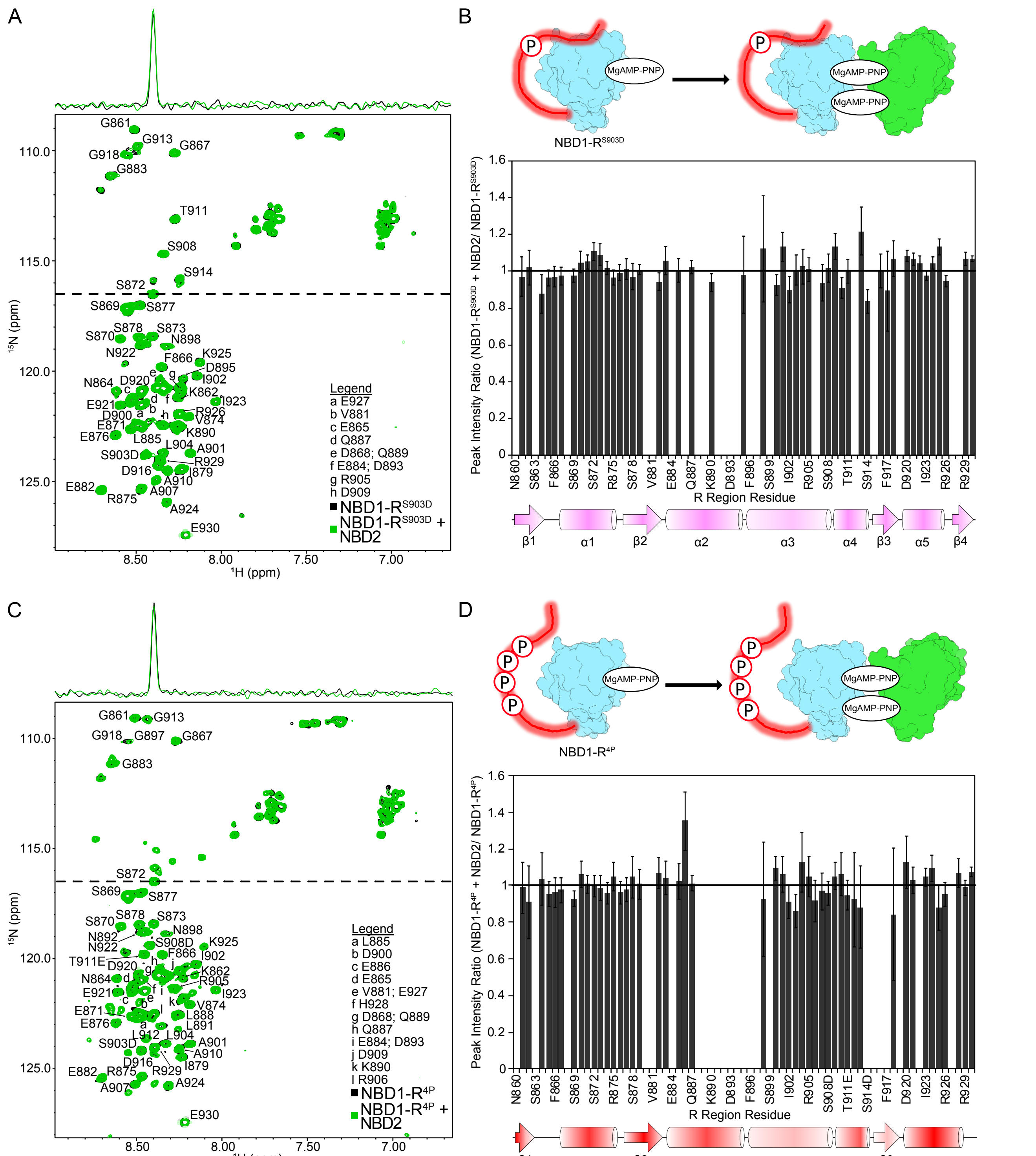


**Supplementary Fig. 19. R$^{S903D}$ and R$^{4P}$ do not interact with NBD2 in the context of NBD1-R$^{S903D}$ and NBD1-R$^{4P}$, respectively.** (A) Overlay of the $^{1}$H-$^{15}$N HSQC spectra, recorded at 4 ºC, of 40 μM MgAMP-PNP-bound NBD1-R$^{S903D}$ in the absence (black, background) and presence (green, foreground) of 160 μM NBD2. A total of 2 mM MgAMP-PNP was used in each sample. The $^{1}$H trace shows the $^{1}$H-$^{15}$N signal corresponding to the dotted line on the spectrum. (B) Schematic representation of the proteins and nucleotide used in the NMR interaction experiments in A (top). Experiments are performed such that NBD1 resonances are silenced and only R$^{S903D}$ peaks are visible, represented by the red glow around R$^{S903D}$. Plot of the peak intensity ratios for R$^{S903D}$ in MgAMP-PNP-bound NBD1-R$^{S903D}$ with and without NBD2 as a function of R region residues (*bottom*). The fact that most peak intensity ratios are ~1 indicates that $R^{S903D}$ does not bind NBD2. Schematic representation of the secondary structure of $R^{S903D}$ from Fig. 2 is shown below the plot. (C) Overlay of the $^{1}H$-$^{15}N$ HSQC spectra, recorded at 4 ºC, of 40 μM MgAMP-PNP-bound NBD1-$R^{4P}$ in the absence (black, background) and presence (green, foreground) of 160 μM NBD2. The $^{1}H$ trace shows the $^{1}H$-$^{15}N$ signal corresponding to the dotted line on the spectrum. (D) Schematic representation of the proteins and nucleotide used in the NMR interaction experiments in A (top). Experiments are performed such that NBD1 resonances are silenced and only $R^{4P}$ peaks are visible, represented by the red glow around R. Plot of the peak intensity ratios for $R^{4P}$ in MgAMP-PNP-bound NBD1-$R^{4P}$ with and without NBD2 as a function of R region residues (*bottom*). The fact that most peak intensity ratios are ~1 indicates that $R^{4P}$ does not bind NBD2. Schematic representation of the secondary structure of $R^{4P}$ from Fig. 2 is shown below the plot. The data shown in panels A-D are representative of experiments repeated from two separate preparations of all proteins.

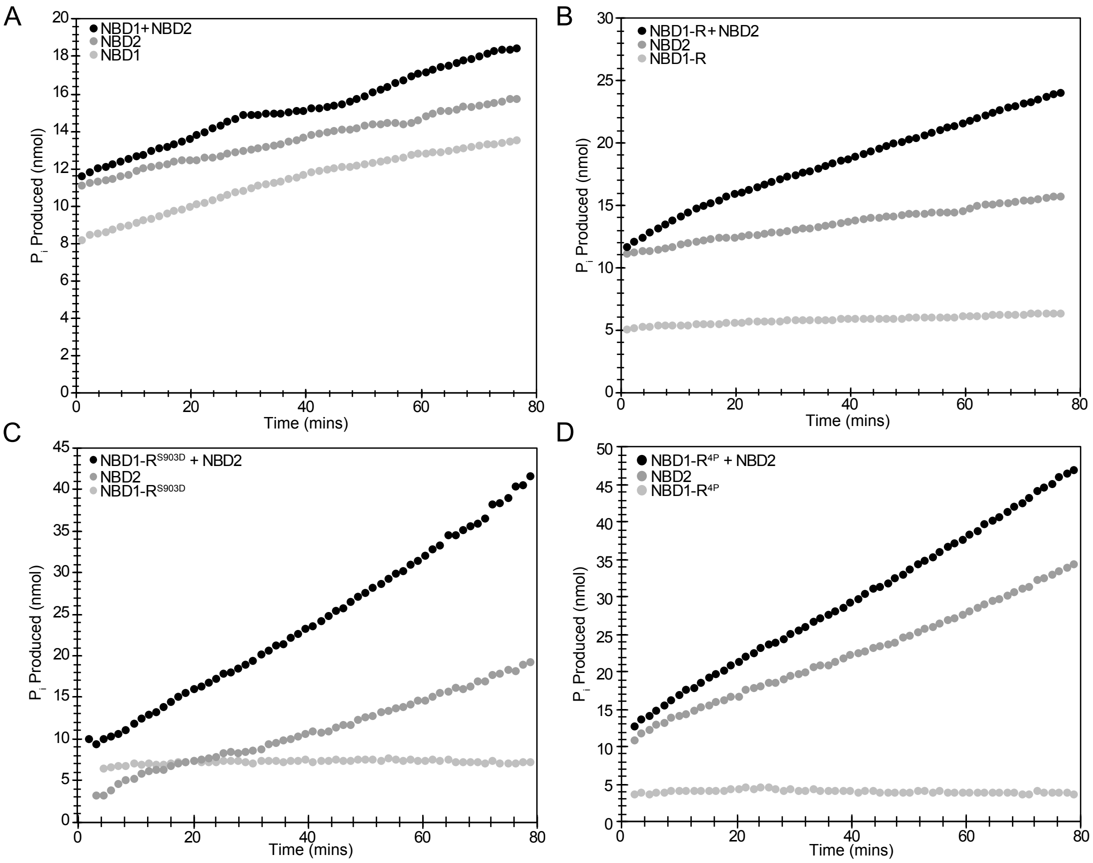


E

| | ATP hydrolysis rate (nmol $P_i$/min) | Number of trials |
|---|---|---|
| NBD1 | 0.22 ± 0.11 | 8 |
| NBD1-R | 0.05 ± 0.03 | 8 |
| NBD1-R$^{S903D}$ | 0.10 ± 0.05 | 9 |
| NBD1-R$^{4P}$ | 0.06 ± 0.07 | 10 |
| NBD2 | 0.17 ± 0.10 | 23 |
| NBD1 + NBD2 | 0.27 ± 0.14 | 8 |
| NBD1-R + NBD2 | 0.22 ± 0.06 | 8 |
| NBD1-R$^{S903D}$ + NBD2 | 0.42 ± 0.12 | 6 |
| NBD1-R$^{4P}$ + NBD2 | 0.46 ± 0.09 | 6 |

**Supplementary Fig. 20. ATPase activity of NBD1-NBD2 heterodimers increases with phosphorylation.** Representation plots showing the $P_i$ production as a function of time by 10 μM NBD2 in the presence of (A) 10 μM NBD1, (B) 10 μM NBD1-R, (C) 10 μM NBD1-R$^{S903D}$, and (D) 10 μM NBD1-R$^{4P}$. Data for the experiments with NBD1 proteins or NBD2 alone are shown in filled light grey and dark grey circles, respectively. $P_i$ production curves for experiments in which both NBD1 proteins and NBD2 are present are shown in filled black circles. Note that experiments shown in each panel were conducted simultaneously. (E) Table of ATPase rates. Note that ATPase rates for NBD1 alone and NBD1/NBD2 were published previously (10), and are shown here for clarity. Data for isolated NBD2 were also shown previously. However, the NBD2 alone data here include additional trials needed as controls for ATPase experiments with NBD2 and NBD1-R proteins. Data were collected with multiple preparations of proteins.

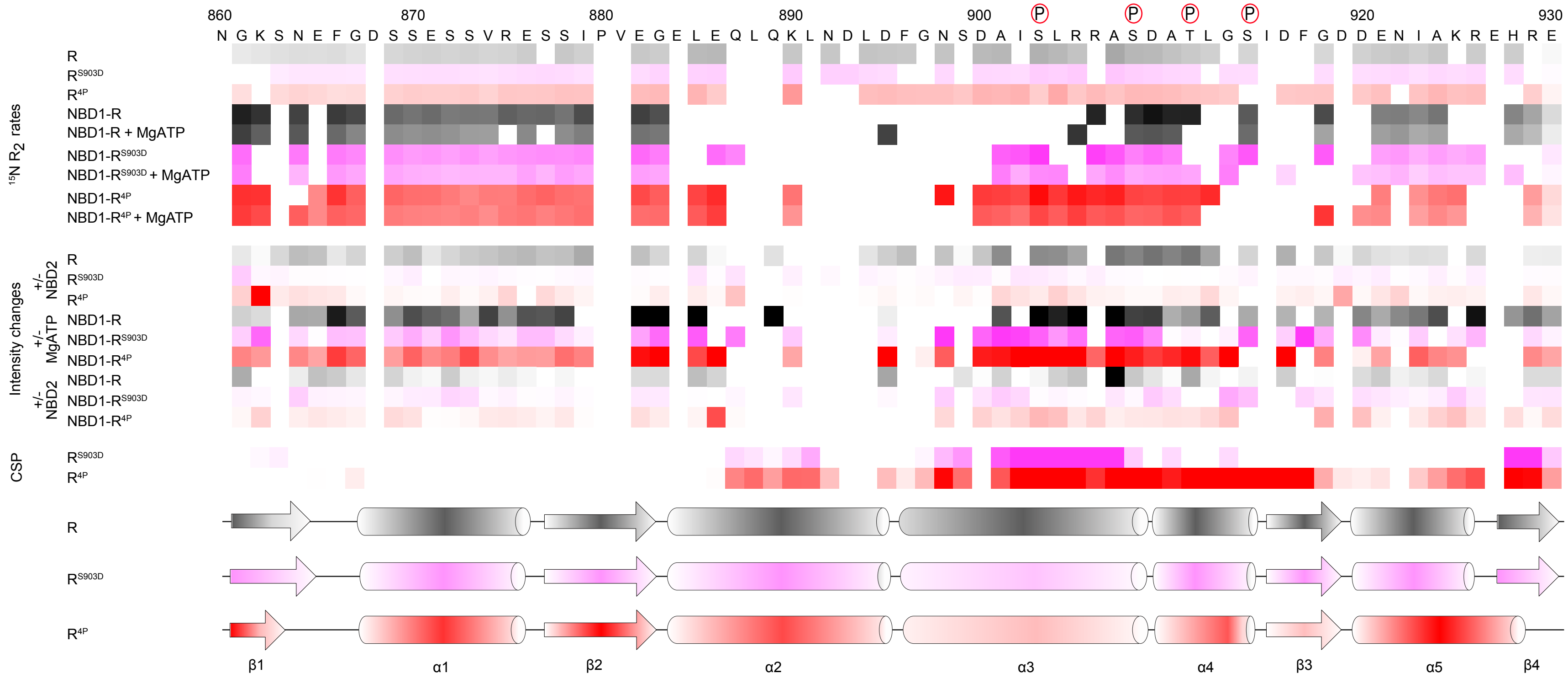


**Supplementary Fig. 21. Summary of changes in the molecular features Ycf1p R region with phosphorylation**. The top panel depicts the representation of $^{15}N$ $R_2$ rates are shown for R region proteins as well as for NBD1-R region proteins without and with MgATP. The middle panel depicts changes in intensities of R region peaks, in the context of the R region alone or as part of NBD1-R proteins, upon addition of NBD2 or MgAMP-PNP alone, or MgAMP-PNP and NBD2. The third set of data show chemical shift perturbations (CSPs) between non-phosphorylated R region and $R^{S903D}$ or $R^{4P}$. Schematic representations of the secondary structure propensities of R region, $R^{S903D}$, and $R^{4P}$, from Fig. 2, are shown below. Data for R and NBD1-R are shown in shades of grey to black, whereas data for RS903 and NBD1-RS903 are shown in shades of pink. Data for $R^{4P}$ and NBD1-$R^{4P}$ are shown in shades of red. The degree of shading indicates the value of the $^{15}N$ $R_2$ rates or CSPs, or the degree of intensity change.
