## Supplementary Table 1 for "Site-specific phosphorylation affects the structure and interactions of the Ycf1p R region"

**Supplementary Table 1. Mass spectrometry of tryptic peptides of non-phosphorylated R region**

| Entry # | Starting Residue | Ending Residue | Peptide sequence (Lowercase letters highlight modified amino acids)* | Probability | Modifications (mass difference)* | Observed | Actual Mass | Charge | Delta Da | Delta PPM | Retention Time | Total Ion Current |
|---|---|---|---|---|---|---|---|---|---|---|---|---|
| 1 | 863 | 875 | (K)snefgdssessvr(E) | 100% | | 709.2769 | 1,416.54 | 2 | 0.004503 | 3.176 | 1,270.78 | 3.34E+08 |
| 2 | 863 | 875 | (K)snefgdssessvr(E) | 100% | | 709.2756 | 1,416.54 | 2 | 0.001983 | 1.399 | 1,311.79 | 3049380 |
| 3 | 863 | 875 | (K)snefgdssessvr(E) | 100% | | 709.2769 | 1,416.54 | 2 | 0.004503 | 3.176 | 1,278.96 | 2.75E+08 |
| 4 | 863 | 875 | (K)snefgdssessvr(E) | 100% | | 709.2693 | 1,416.52 | 2 | -0.01062 | -7.49 | 1,401.56 | 3174730 |
| 5 | 863 | 875 | (K)snefgdssessvr(E) | 100% | | 709.2764 | 1,416.54 | 2 | 0.003583 | 2.527 | 1,327.93 | 2095070 |
| 6 | 863 | 875 | (K)snefgdssessvr(E) | 100% | | 709.2769 | 1,416.54 | 2 | 0.004503 | 3.176 | 1,287.02 | 3.55E+07 |
| 7 | 863 | 875 | (K)snefgdssessvr(E) | 100% | | 709.2769 | 1,416.54 | 2 | 0.004503 | 3.176 | 1,295.28 | 9680010 |
| 8 | 863 | 875 | (K)snefgdssessvr(E) | 100% | | 473.1861 | 1,416.54 | 3 | 0.001807 | 1.274 | 1,275.67 | 1947540 |
| 9 | 863 | 875 | (K)snefgdssessvr(E) | 100% | | 709.2756 | 1,416.54 | 2 | 0.002043 | 1.441 | 1,303.52 | 5066680 |
| 10 | 863 | 875 | (K)snefgdssessvr(E) | 100% | | 709.2725 | 1,416.53 | 2 | -0.004297 | -3.032 | 1,352.31 | 1364840 |
| 11 | 863 | 875 | (K)snefgdssessvr(E) | 100% | | 709.2693 | 1,416.52 | 2 | -0.01062 | -7.49 | 1,393.50 | 2034900 |
| 12 | 863 | 875 | (K)snefgdssessvr(E) | 100% | | 709.2747 | 1,416.53 | 2 | 0.0001025 | 0.07234 | 1,262.44 | 690661 |
| 13 | 863 | 875 | (K)snefgdssessvr(E) | 100% | | 709.275 | 1,416.54 | 2 | 0.0007825 | 0.552 | 1,319.87 | 2203860 |
| 14 | 863 | 875 | (K)snefgdssessvr(E) | 100% | | 709.2764 | 1,416.54 | 2 | 0.003583 | 2.527 | 1,336.01 | 8831330 |
| 15 | 863 | 875 | (K)snefgdssessvr(E) | 100% | | 709.2725 | 1,416.53 | 2 | -0.004297 | -3.032 | 1,360.43 | 669473 |
| 16 | 863 | 875 | (K)snefgdssessvr(E) | 100% | | 709.2725 | 1,416.53 | 2 | -0.004297 | -3.032 | 1,344.16 | 3746620 |
| 17 | 863 | 875 | (K)snefgdssessvr(E) | 100% | | 709.2725 | 1,416.53 | 2 | -0.004297 | -3.032 | 1,368.69 | 573447 |
| 18 | 863 | 875 | (K)snefgdssessvr(E) | 100% | | 709.2693 | 1,416.52 | 2 | -0.01062 | -7.49 | 1,409.69 | 1151920 |
| 19 | 863 | 875 | (K)snefgdssessvr(E) | 100% | | 709.2725 | 1,416.53 | 2 | -0.004297 | -3.032 | 1,376.98 | 476126 |
| 20 | 863 | 875 | (K)snefgdssessvr(E) | 100% | | 709.2725 | 1,416.53 | 2 | -0.004297 | -3.032 | 1,385.27 | 373915 |
| 21 | 863 | 875 | (K)snefgdssessvr(E) | 100% | | 473.1861 | 1,416.54 | 3 | 0.001807 | 1.274 | 1,267.28 | 300643 |
| 22 | 863 | 875 | (K)snefgdssessvr(E) | 100% | | 473.1861 | 1,416.54 | 3 | 0.001807 | 1.274 | 1,283.93 | 332888 |
| 23 | 863 | 875 | (K)snefgdssessvr(E) | 100% | | 709.2731 | 1,416.53 | 2 | -0.003017 | -2.129 | 1,511.11 | 377065 |
| 24 | 875 | 890 | (V)ressipvegeleqlqk(L) | 100% | | 621.9706 | 1,862.89 | 3 | 0.001931 | 1.036 | 2,091.35 | 607520 |
| 25 | 875 | 890 | (V)ressipvegeleqlqk(L) | 100% | | 621.9706 | 1,862.89 | 3 | 0.001931 | 1.036 | 2,083.15 | 551074 |
| 26 | 875 | 890 | (V)ressipvegeleqlqk(L) | 100% | | 621.9706 | 1,862.89 | 3 | 0.001931 | 1.036 | 2,075.18 | 343834 |
| 27 | 876 | 890 | (R)essipvegeleqlqk(L) | 100% | | 568.608 | 1,702.80 | 3 | 0.003462 | 2.032 | 2,598.50 | 1.81E+07 |
| 28 | 876 | 890 | (R)essipvegeleqlqk(L) | 100% | | 568.608 | 1,702.80 | 3 | 0.003462 | 2.032 | 2,582.18 | 1.08E+07 |
| 29 | 876 | 890 | (R)essipvegeleqlqk(L) | 100% | | 568.608 | 1,702.80 | 3 | 0.003462 | 2.032 | 2,590.38 | 2.29E+07 |
| 30 | 876 | 890 | (R)essipvegeleqlqk(L) | 100% | | 852.4101 | 1,702.81 | 2 | 0.006848 | 4.019 | 2,592.80 | 8.29E+07 |
| 31 | 876 | 890 | (R)essipvegeleqlqk(L) | 100% | | 568.608 | 1,702.80 | 3 | 0.003462 | 2.032 | 2,606.83 | 4313500 |
| 32 | 876 | 890 | (R)essipvegeleqlqk(L) | 100% | | 852.4101 | 1,702.81 | 2 | 0.006848 | 4.019 | 2,568.20 | 2453130 |
| 33 | 876 | 890 | (R)essipvegeleqlqk(L) | 100% | | 852.4101 | 1,702.81 | 2 | 0.006848 | 4.019 | 2,584.69 | 5.32E+07 |
| 34 | 876 | 890 | (R)essipvegeleqlqk(L) | 100% | | 852.4101 | 1,702.81 | 2 | 0.006848 | 4.019 | 2,600.95 | 5.52E+07 |
| 35 | 876 | 890 | (R)essipvegeleqlqk(L) | 100% | | 852.4101 | 1,702.81 | 2 | 0.006848 | 4.019 | 2,576.41 | 1.20E+07 |
| 36 | 876 | 890 | (R)essipvegeleqlqk(L) | 100% | | 568.608 | 1,702.80 | 3 | 0.003462 | 2.032 | 2,573.88 | 1934200 |
| 37 | 876 | 890 | (R)essipvegeleqlqk(L) | 100% | | 568.608 | 1,702.80 | 3 | 0.003462 | 2.032 | 2,614.91 | 767828 |
| 38 | 876 | 890 | (R)essipvegeleqlqk(L) | 100% | | 852.4101 | 1,702.81 | 2 | 0.006848 | 4.019 | 2,609.06 | 9576840 |
| 39 | 876 | 890 | (R)essipvegeleqlqk(L) | 100% | | 852.4101 | 1,702.81 | 2 | 0.006848 | 4.019 | 2,617.25 | 2186330 |
| 40 | 876 | 890 | (R)essipvegeleqlqk(L) | 100% | | 852.4101 | 1,702.81 | 2 | 0.006848 | 4.019 | 2,625.36 | 986233 |
| 41 | 876 | 890 | (R)essipvegeleqlqk(L) | 100% | | 852.4078 | 1,702.80 | 2 | 0.002328 | 1.366 | 2,633.54 | 526804 |
| 42 | 876 | 890 | (R)essipvegeleqlqk(L) | 100% | | 852.407 | 1,702.80 | 2 | 0.0007275 | 0.427 | 2,641.66 | 368172 |
| 43 | 876 | 890 | (R)essipvegeleqlqk(L) | 100% | | 568.608 | 1,702.80 | 3 | 0.003462 | 2.032 | 2,622.98 | 242271 |
| 44 | 876 | 890 | (R)essipvegeleqlqk(L) | 100% | | 852.4056 | 1,702.80 | 2 | -0.002152 | -1.263 | 2,658.21 | 317088 |
| 45 | 876 | 890 | (R)essipvegeleqlqk(L) | 100% | | 568.608 | 1,702.80 | 3 | 0.003462 | 2.032 | 2,565.81 | 209326 |
| 46 | 880 | 890 | (I)pvegeleqlqk(L) | 100% | | 642.3166 | 1,282.62 | 2 | -0.001212 | -0.9445 | 2,592.16 | 747707 |
| 47 | 891 | 905 | (K)lndldfgnsdaislr(R) | 100% | | 835.3816 | 1,668.75 | 2 | 0.002077 | 1.244 | 2,837.75 | 2004460 |
| 48 | 891 | 905 | (K)lndldfgnsdaislr(R) | 100% | | 835.3816 | 1,668.75 | 2 | 0.002077 | 1.244 | 2,829.70 | 2211000 |
| 49 | 891 | 905 | (K)lndldfgnsdaislr(R) | 100% | | 835.3816 | 1,668.75 | 2 | 0.002077 | 1.244 | 2,845.88 | 348479 |
| 50 | 891 | 905 | (K)lndldfgnsdaislr(R) | 100% | | 835.3816 | 1,668.75 | 2 | 0.002077 | 1.244 | 2,821.54 | 313809 |
| 51 | 891 | 905 | (K)lndldfgnsdaislr(R) | 100% | | 835.3744 | 1,668.73 | 2 | -0.01232 | -7.38 | 3,056.56 | 273631 |
| 52 | 891 | 906 | (K)lndldfgnsdaislrr(A) | 100% | | 610.6232 | 1,828.85 | 3 | 0.01187 | 6.488 | 2,267.64 | 2.30E+07 |
| 53 | 891 | 906 | (K)lndldfgnsdaislrr(A) | 100% | | 610.6212 | 1,828.84 | 3 | 0.005961 | 3.258 | 2,259.44 | 9843840 |
| 54 | 891 | 906 | (K)lndldfgnsdaislrr(A) | 100% | | 610.6204 | 1,828.84 | 3 | 0.003441 | 1.881 | 2,284.25 | 2276780 |
| 55 | 891 | 906 | (K)lndldfgnsdaislrr(A) | 100% | | 610.6224 | 1,828.85 | 3 | 0.009561 | 5.225 | 2,275.92 | 1.11E+07 |
| 56 | 891 | 906 | (K)lndldfgnsdaislrr(A) | 100% | | 610.6201 | 1,828.84 | 3 | 0.002541 | 1.389 | 2,292.54 | 749249 |
| 57 | 891 | 906 | (K)lndldfgnsdaislrr(A) | 100% | | 610.6163 | 1,828.83 | 3 | -0.008739 | -4.776 | 2,451.09 | 800247 |
| 58 | 891 | 906 | (K)lndldfgnsdaislrr(A) | 100% | | 610.6199 | 1,828.84 | 3 | 0.002061 | 1.127 | 2,308.84 | 440344 |
| 59 | 891 | 906 | (K)lndldfgnsdaislrr(A) | 100% | | 610.6195 | 1,828.84 | 3 | 0.0008614 | 0.4707 | 2,251.10 | 439204 |
| 60 | 891 | 906 | (K)lndldfgnsdaislrr(A) | 100% | | 610.6202 | 1,828.84 | 3 | 0.003081 | 1.684 | 2,442.95 | 665268 |
| 61 | 891 | 906 | (K)lndldfgnsdaislrr(A) | 100% | | 610.6199 | 1,828.84 | 3 | 0.002151 | 1.176 | 2,300.71 | 465803 |
| 62 | 896 | 905 | (D)fgnsdaislr(R) | 97% | | 547.2571 | 1,092.50 | 2 | 0.0003876 | 0.3545 | 1,734.03 | 245010 |
| 63 | 906 | 925 | (R)rasdatlgsidfgddeniak(R) | 100% | | 707.3131 | 2,118.92 | 3 | 0.004576 | 2.159 | 2,222.98 | 9672930 |
| 64 | 906 | 925 | (R)rasdatlgsidfgddeniak(R) | 100% | | 707.3118 | 2,118.91 | 3 | 0.0007363 | 0.3473 | 2,214.73 | 493944 |
| 65 | 906 | 926 | (R)rasdatlgSidfgddeniakr(E) | 100% | | 570.5104 | 2,278.01 | 4 | 0.007705 | 3.381 | 1,910.68 | 2331470 |
| 66 | 906 | 926 | (R)rasdatlgSidfgddeniakr(E) | 100% | | 570.5104 | 2,278.01 | 4 | 0.007705 | 3.381 | 1,918.75 | 2561940 |
| 67 | 906 | 926 | (R)rasdatlgsidfgddeniakr(E) | 100% | | 570.7596 | 2,279.01 | 4 | 0.00731 | 3.206 | 1,918.75 | 2561940 |
| 68 | 906 | 926 | (R)rasdatlgSidfgddeniakr(E) | 100% | | 570.5104 | 2,278.01 | 4 | 0.007705 | 3.381 | 1,926.89 | 874131 |
| 69 | 906 | 926 | (R)rasdatlgsidfgddeniakr(E) | 100% | | 570.7583 | 2,279.00 | 4 | 0.00211 | 0.9255 | 1,910.68 | 2331470 |
| 70 | 906 | 926 | (R)rasdatlgSidfgddeniakr(E) | 100% | | 760.3422 | 2,278.00 | 3 | -0.0002987 | -0.1311 | 1,914.62 | 930782 |
| 71 | 906 | 926 | (R)rasdatlgsidfgddeniakr(E) | 100% | | 760.675 | 2,279.00 | 3 | 0.001186 | 0.5203 | 1,914.62 | 930782 |
| 72 | 906 | 926 | (R)rasdatlgsidfgddeniakr(E) | 100% | | 570.7587 | 2,279.01 | 4 | 0.00359 | 1.575 | 1,926.89 | 874131 |
| 73 | 906 | 926 | (R)rasdatlgSidfgddeniakr(E) | 100% | | 760.3422 | 2,278.00 | 3 | -0.0002987 | -0.1311 | 1,922.75 | 534913 |

Additional Notes:
*R region was 15N-labeled and thus, all N aroms are 15N.
The 15N-modification is not listed above for ease of reading.

**Supplementary Table 1. Mass spectrometry of tryptic peptides of non-phosphorylated R region**

| **Entry #** | **Starting Residue** | **Ending Residue** | **Peptide sequence** (Lowercase letters highlight modified amino acids)* | **Probability** | **Modifications** (mass difference)* | **Observed** | **Actual Mass** | **Charge** | **Delta Da** | **Delta PPM** | **Retention Time** | **Total Ion Current** |
|---|---|---|---|---|---|---|---|---|---|---|---|---|
| 74 | 906 | 926 | (R)rasdatlgSidfgddeniakr(E) | 100% | | 570.5104 | 2,278.01 | 4 | 0.007705 | 3.381 | 1,902.62 | 391538 |
| 75 | 906 | 926 | (R)rasdatlgsidfgddeniakr(E) | 100% | | 760.6747 | 2,279.00 | 3 | 0.0004062 | 0.1782 | 1,922.75 | 534913 |
| 76 | 906 | 926 | (R)rasdatlgsidfgddeniakr(E) | 100% | | 570.7578 | 2,279.00 | 4 | 0.0001102 | 0.04835 | 1,902.62 | 391538 |
| 77 | 906 | 926 | (R)rasdatlgsidfgddeniakr(E) | 100% | | 760.6754 | 2,279.00 | 3 | 0.002386 | 1.047 | 1,906.57 | 403726 |
| 78 | 906 | 926 | (R)rasdatlgSidfgddeniakr(E) | 100% | | 760.3422 | 2,278.00 | 3 | -0.0002987 | -0.1311 | 1,906.57 | 403726 |
| 79 | 907 | 925 | (R)asdatlgsidfgddeniak(R) | 100% | | 653.9497 | 1,958.83 | 3 | 0.003766 | 1.922 | 2,682.87 | 6494260 |
| 80 | 907 | 925 | (R)asdatlgSidfgddeniak(R) | 100% | | 653.6177 | 1,957.83 | 3 | 0.004801 | 2.451 | 2,691.08 | 824286 |
| 81 | 907 | 925 | (R)asdatlgsidfgddeniak(R) | 100% | | 653.9493 | 1,958.83 | 3 | 0.002686 | 1.371 | 2,691.08 | 824286 |
| 82 | 907 | 925 | (R)asdatlgsidfgddeniak(R) | 100% | | 980.4227 | 1,958.83 | 2 | 0.007282 | 3.716 | 2,680.08 | 570478 |
| 83 | 907 | 925 | (R)asdatlgsidfgddeniak(R) | 100% | | 653.9498 | 1,958.83 | 3 | 0.003976 | 2.029 | 2,674.60 | 7112370 |
| 84 | 907 | 925 | (R)asdatlgSidfgddeniak(R) | 100% | | 653.6177 | 1,957.83 | 3 | 0.004801 | 2.451 | 2,682.87 | 6494260 |
| 85 | 907 | 925 | (R)asdatlgSidfgddeniak(R) | 100% | | 653.6177 | 1,957.83 | 3 | 0.004801 | 2.451 | 2,666.30 | 194919 |
| 86 | 907 | 925 | (R)asdatlgsidfgddeniak(R) | 100% | | 980.4227 | 1,958.83 | 2 | 0.007282 | 3.716 | 2,672.01 | 213940 |
| 87 | 907 | 925 | (R)asdatlgsidfgddeniak(R) | 100% | | 653.949 | 1,958.83 | 3 | 0.001786 | 0.9115 | 2,699.39 | 173682 |
| 88 | 907 | 925 | (R)asdaTlgsidfgddeniak(R) | 100% | | 653.6177 | 1,957.83 | 3 | 0.004801 | 2.451 | 2,699.39 | 173682 |
| 89 | 907 | 925 | (R)asdatlgSidfgddeniak(R) | 100% | | 653.6177 | 1,957.83 | 3 | 0.004801 | 2.451 | 2,674.60 | 7112370 |
| 90 | 907 | 925 | (R)asdatlgsidfgddeniak(R) | 100% | | 653.9492 | 1,958.83 | 3 | 0.002266 | 1.156 | 2,666.30 | 194919 |
| 91 | 907 | 925 | (R)asdatlgsidfgddeniak(R) | 100% | | 980.4227 | 1,958.83 | 2 | 0.007282 | 3.716 | 2,688.29 | 76,930.30 |
| 92 | 907 | 926 | (R)asdatlgsidfgddeniakr(E) | 100% | | 707.3121 | 2,118.91 | 3 | 0.001726 | 0.8143 | 2,263.73 | 1779470 |
| 93 | 907 | 926 | (R)asdatlgsidfgddeniakr(E) | 100% | | 707.3166 | 2,118.93 | 3 | 0.01523 | 7.182 | 2,247.39 | 3.07E+07 |
| 94 | 907 | 926 | (R)asdatlgsidfgddeniakr(E) | 100% | | 707.3171 | 2,118.93 | 3 | 0.01673 | 7.89 | 2,239.32 | 6.27E+07 |
| 95 | 907 | 926 | (R)asdatlgsidfgddeniakr(E) | 100% | | 707.3132 | 2,118.92 | 3 | 0.005026 | 2.371 | 2,255.66 | 7004330 |
| 96 | 907 | 926 | (R)asdatlgsidfgddeniakr(E) | 100% | | 707.3119 | 2,118.91 | 3 | 0.001126 | 0.5313 | 2,271.97 | 645415 |
| 97 | 907 | 926 | (R)asdatlgsidfgddeniakr(E) | 100% | | 707.3127 | 2,118.92 | 3 | 0.003646 | 1.72 | 2,296.55 | 926424 |
| 98 | 907 | 926 | (R)asdatlgsidfgddeniakr(E) | 100% | | 707.3167 | 2,118.93 | 3 | 0.01553 | 7.324 | 2,231.07 | 4.73E+07 |
| 99 | 907 | 926 | (R)asdatlgsidfgddeniakr(E) | 100% | | 707.3131 | 2,118.92 | 3 | 0.004576 | 2.159 | 2,288.50 | 1005150 |
| 100 | 907 | 926 | (R)asdatlgsidfgddeniakr(E) | 100% | | 707.3118 | 2,118.91 | 3 | 0.0007363 | 0.3473 | 2,280.25 | 487755 |
| 101 | 907 | 926 | (R)asdatlgsidfgddeniakr(E) | 100% | | 707.3126 | 2,118.92 | 3 | 0.003106 | 1.465 | 2,304.75 | 416425 |
| 102 | 907 | 929 | (R)asdatlgSidfgddeniakrehr(E) | 100% | | 638.0313 | 2,548.10 | 4 | 0.001505 | 0.5905 | 1,769.58 | 439859 |
| 103 | 907 | 929 | (R)asdaTlgsidfgddeniakrehr(E) | 100% | | 638.0313 | 2,548.10 | 4 | 0.001505 | 0.5905 | 1,761.25 | 552157 |
| 104 | 907 | 929 | (R)asdatlgsidfgddeniakrehr(E) | 100% | | 638.2806 | 2,549.09 | 4 | 0.00183 | 0.7177 | 1,769.58 | 439859 |
| 105 | 907 | 929 | (R)asdaTlgsidfgddeniakrehr(E) | 100% | | 510.6272 | 2,548.10 | 5 | 0.005029 | 1.973 | 1,769.33 | 397260 |
| 106 | 907 | 929 | (R)asdatlgsidfgddeniakrehr(E) | 100% | | 638.2804 | 2,549.09 | 4 | 0.0008701 | 0.3412 | 1,761.25 | 552157 |
| 107 | 907 | 929 | (R)asdatlgsidfgddeniakrehr(E) | 100% | | 510.8267 | 2,549.10 | 5 | 0.005294 | 2.076 | 1,769.33 | 397260 |
| 108 | 907 | 930 | (R)asdatlgSidfgddeniakrehre(Q) | 100% | | 536.6343 | 2,678.14 | 5 | 0.0008941 | 0.3337 | 1,763.48 | 937283 |
| 109 | 907 | 930 | (R)asdatlgsidfgddeniakrehre(Q) | 100% | | 670.7904 | 2,679.13 | 4 | 0.001235 | 0.4608 | 1,756.52 | 1064750 |
| 110 | 907 | 930 | (R)asdatlgSidfgddeniakrehre(Q) | 97% | | 536.6343 | 2,678.14 | 5 | 0.0008941 | 0.3337 | 1,755.29 | 877056 |

Additional Notes:
*R region was 15N-labeled and thus, all N aroms are 15N.
The 15N-modification is not listed above for ease of reading.
