## Supplementary Table 2 for "Site-specific phosphorylation affects the structure and interactions of the Ycf1p R region"

**Supplementary Table 2. Mass spectrometry of tryptic peptides of PKA-treated Ycf1p R region**

| Entry # | Starting Residue | Ending Residue | **Peptide sequence** (Lowercase letters highlight modified amino acids)* | Probability | **Modifications** (mass difference)* | Observed | Actual Mass | Charge | Delta Da | Delta PPM | Retention Time | Total Ion Current |
|---|---|---|---|---|---|---|---|---|---|---|---|---|
| 1 | 861 | 875 | (N)gksnefgdssessvr(E) | 100% | | 535.8887 | 1,604.64 | 3 | 0.002091 | 1.303 | | 375114 |
| 2 | 863 | 875 | (K)snefgdssessvr(E) | 100% | | 709.2764 | 1,416.54 | 2 | 0.003523 | 2.485 | | 6692120 |
| 3 | 863 | 875 | (K)snefgdssessvr(E) | 100% | | 709.275 | 1,416.54 | 2 | 0.0007025 | 0.4956 | | 534494 |
| 4 | 863 | 875 | (K)snefgdssessvr(E) | 100% | | 709.2758 | 1,416.54 | 2 | 0.002303 | 1.624 | | 5257150 |
| 5 | 863 | 875 | (K)snefgdssessvr(E) | 100% | | 709.2769 | 1,416.54 | 2 | 0.004503 | 3.176 | | 2.35E+08 |
| 6 | 863 | 875 | (K)snefgdssessvr(E) | 100% | | 709.2743 | 1,416.53 | 2 | -0.0006175 | -0.4356 | | 1039220 |
| 7 | 863 | 875 | (K)snefgdssessvr(E) | 100% | | 709.2769 | 1,416.54 | 2 | 0.004503 | 3.176 | | 3.98E+08 |
| 8 | 863 | 875 | (K)snefgdssessvr(E) | 100% | | 709.2769 | 1,416.54 | 2 | 0.004503 | 3.176 | | 2.70E+07 |
| 9 | 863 | 875 | (K)snefgdssessvr(E) | 100% | | 709.2764 | 1,416.54 | 2 | 0.003523 | 2.485 | | 2574670 |
| 10 | 863 | 875 | (K)snefgdssessvr(E) | 100% | | 709.2722 | 1,416.53 | 2 | -0.004897 | -3.455 | | 2453050 |
| 11 | 863 | 875 | (K)snefgdssessvr(E) | 100% | | 709.2701 | 1,416.53 | 2 | -0.009017 | -6.361 | | 1032760 |
| 12 | 863 | 875 | (K)snefgdssessvr(E) | 100% | | 709.2764 | 1,416.54 | 2 | 0.003523 | 2.485 | | 3754130 |
| 13 | 863 | 875 | (K)snefgdssessvr(E) | 100% | | 709.2751 | 1,416.54 | 2 | 0.0009825 | 0.6931 | | 3426610 |
| 14 | 863 | 875 | (K)snefgdssessvr(E) | 100% | | 709.273 | 1,416.53 | 2 | -0.003317 | -2.34 | | 551870 |
| 15 | 863 | 875 | (K)snefgdssessvr(E) | 100% | | 709.2769 | 1,416.54 | 2 | 0.004503 | 3.176 | | 1.05E+07 |
| 16 | 863 | 875 | (K)snefgdssessvr(E) | 100% | | 709.2748 | 1,416.54 | 2 | 0.0003825 | 0.2699 | | 713939 |
| 17 | 863 | 875 | (K)snefgdssessvr(E) | 100% | | 709.2754 | 1,416.54 | 2 | 0.001683 | 1.187 | | 576731 |
| 18 | 863 | 875 | (K)snefgdssessvr(E) | 100% | | 473.1859 | 1,416.54 | 3 | 0.001147 | 0.8088 | | 339772 |
| 19 | 863 | 875 | (K)snefgdssessvr(E) | 100% | | 709.2701 | 1,416.53 | 2 | -0.009017 | -6.361 | | 1314300 |
| 20 | 863 | 875 | (K)snefgdssessvr(E) | 100% | | 709.2701 | 1,416.53 | 2 | -0.009017 | -6.361 | | 405800 |
| 21 | 863 | 875 | (K)snefgdssessvr(E) | 100% | | 473.1859 | 1,416.54 | 3 | 0.001147 | 0.8088 | | 3099890 |
| 22 | 863 | 875 | (K)snefgdssessvr(E) | 100% | | 709.2745 | 1,416.53 | 2 | -0.0002175 | -0.1534 | | 650402 |
| 23 | 863 | 875 | (K)snefgdssessvr(E) | 100% | | 709.2745 | 1,416.53 | 2 | -0.0001375 | -0.09696 | | 474693 |
| 24 | 863 | 875 | (K)snefgdssessvr(E) | 100% | | 473.1859 | 1,416.54 | 3 | 0.001147 | 0.8088 | | 422218 |
| 25 | 865 | 875 | (N)efgdssessvr(E) | 100% | | 607.2417 | 1,212.47 | 2 | 0.0001876 | 0.1546 | 1,254.25 | 329394 |
| 26 | 876 | 890 | (R)essipvegeleqlqk(L) | 100% | | 568.6074 | 1,702.80 | 3 | 0.001572 | 0.9224 | | 2004710 |
| 27 | 876 | 890 | (R)essipvegeleqlqk(L) | 100% | | 568.6074 | 1,702.80 | 3 | 0.001572 | 0.9224 | | 2.00E+07 |
| 28 | 876 | 890 | (R)essipvegeleqlqk(L) | 100% | | 568.6074 | 1,702.80 | 3 | 0.001572 | 0.9224 | | 9991940 |
| 29 | 876 | 890 | (R)essipvegeleqlqk(L) | 100% | | 568.6074 | 1,702.80 | 3 | 0.001572 | 0.9224 | | 5448060 |
| 30 | 876 | 890 | (R)essipvegeleqlqk(L) | 100% | | 568.6074 | 1,702.80 | 3 | 0.001572 | 0.9224 | | 1.56E+07 |
| 31 | 876 | 890 | (R)essipvegeleqlqk(L) | 100% | | 852.4104 | 1,702.81 | 2 | 0.007448 | 4.371 | 2,580.43 | 6.92E+07 |
| 32 | 876 | 890 | (R)essipvegeleqlqk(L) | 100% | | 852.4104 | 1,702.81 | 2 | 0.007448 | 4.371 | | 5.19E+07 |
| 33 | 876 | 890 | (R)essipvegeleqlqk(L) | 100% | | 852.4104 | 1,702.81 | 2 | 0.007448 | 4.371 | | 9.09E+07 |
| 34 | 876 | 890 | (R)essipvegeleqlqk(L) | 100% | | 852.4104 | 1,702.81 | 2 | 0.007448 | 4.371 | 2,605.09 | 2.11E+07 |
| 35 | 876 | 890 | (R)essipvegeleqlqk(L) | 100% | | 568.6074 | 1,702.80 | 3 | 0.001572 | 0.9224 | | 1189780 |
| 36 | 876 | 890 | (R)essipvegeleqlqk(L) | 100% | | 852.4104 | 1,702.81 | 2 | 0.007448 | 4.371 | 2,564.17 | 1305370 |
| 37 | 876 | 890 | (R)essipvegeleqlqk(L) | 100% | | 852.4104 | 1,702.81 | 2 | 0.007448 | 4.371 | 2,572.36 | 1.24E+07 |
| 38 | 876 | 890 | (R)essipvegeleqlqk(L) | 100% | | 852.4104 | 1,702.81 | 2 | 0.007448 | 4.371 | | 1842100 |
| 39 | 876 | 890 | (R)essipvegeleqlqk(L) | 100% | | 852.4104 | 1,702.81 | 2 | 0.007448 | 4.371 | | 4303980 |
| 40 | 876 | 890 | (R)essipvegeleqlqk(L) | 100% | | 852.4047 | 1,702.79 | 2 | -0.004032 | -2.367 | 2,661.97 | 764325 |
| 41 | 876 | 890 | (R)essipvegeleqlqk(L) | 100% | | 852.4104 | 1,702.81 | 2 | 0.007448 | 4.371 | 2,629.54 | 805060 |
| 42 | 876 | 890 | (R)essipvegeleqlqk(L) | 100% | | 852.4056 | 1,702.80 | 2 | -0.002152 | -1.263 | | 603848 |
| 43 | 876 | 890 | (R)essipvegeleqlqk(L) | 100% | | 568.6074 | 1,702.80 | 3 | 0.001572 | 0.9224 | | 380203 |
| 44 | 876 | 890 | (R)essipvegeleqlqk(L) | 100% | | 852.4076 | 1,702.80 | 2 | 0.001948 | 1.143 | | 531136 |
| 45 | 876 | 890 | (R)essipvegeleqlqk(L) | 100% | | 852.4068 | 1,702.80 | 2 | 0.0002475 | 0.1453 | | 348109 |
| 46 | 876 | 890 | (R)essipvegeleqlqk(L) | 100% | | 568.6074 | 1,702.80 | 3 | 0.001572 | 0.9224 | | 254557 |
| 47 | 880 | 890 | (I)pvegeleqlqk(L) | 100% | | 642.3188 | 1,282.62 | 2 | 0.003188 | 2.483 | | 638175 |
| 48 | 880 | 890 | (I)pvegeleqlqk(L) | 100% | | 642.316 | 1,282.62 | 2 | -0.002432 | -1.895 | | 543723 |
| 49 | 881 | 890 | (P)vegeleqlqk(L) | 100% | | 593.2922 | 1,184.57 | 2 | -0.0001973 | -0.1665 | | 263601 |
| 50 | 891 | 905 | (K)lndldfgnsdaislr(R) | 100% | | 835.3829 | 1,668.75 | 2 | 0.004677 | 2.801 | | 1882160 |
| 51 | 891 | 905 | (K)lndldfgnsdaislr(R) | 100% | | 835.3829 | 1,668.75 | 2 | 0.004677 | 2.801 | | 3811060 |
| 52 | 891 | 905 | (K)lndldfgnsdaislr(R) | 100% | | 835.375 | 1,668.74 | 2 | -0.01112 | -6.661 | | 1087450 |
| 53 | 891 | 905 | (K)lndldfgnsdaislr(R) | 100% | | 835.3829 | 1,668.75 | 2 | 0.004677 | 2.801 | | 5833670 |
| 54 | 891 | 905 | (K)lndldfgnsdaislr(R) | 100% | | 835.3815 | 1,668.75 | 2 | 0.001877 | 1.124 | | 442265 |
| 55 | 891 | 905 | (K)lndldfgnsdaislr(R) | 100% | | 835.3829 | 1,668.75 | 2 | 0.004677 | 2.801 | | 585063 |
| 56 | 891 | 905 | (K)lndldfgnsdaislr(R) | 100% | | 835.3829 | 1,668.75 | 2 | 0.004677 | 2.801 | | 215036 |
| 57 | 891 | 905 | (K)lndldfgnsdaislr(R) | 100% | | 557.257 | 1,668.75 | 3 | 0.002601 | 1.558 | | 259993 |
| 58 | 891 | 905 | (K)lndldfgnsdaislr(R) | 100% | | 835.3829 | 1,668.75 | 2 | 0.004677 | 2.801 | | 161615 |
| 59 | 891 | 905 | (K)lndldfgnsdaislr(R) | 100% | | 557.257 | 1,668.75 | 3 | 0.002601 | 1.558 | | 181580 |

Additional Notes:
*R region was 15N-labeled and thus, all N aroms are 15N.
The 15N-modification is not listed above for ease of reading.
*Phosphorylated residues are bolded and underlined.

**Supplementary Table 2. Mass spectrometry of tryptic peptides of PKA-treated Ycf1p R region**

| Entry # | Starting Residue | Ending Residue | **Peptide sequence** (Lowercase letters highlight modified amino acids)* | Probability | **Modifications** (mass difference)* | Observed | Actual Mass | Charge | Delta Da | Delta PPM | Retention Time | Total Ion Current |
|---|---|---|---|---|---|---|---|---|---|---|---|---|
| 60 | 891 | 905 | (K)lndldfgnsdaislr(R) | 100% | | 835.375 | 1,668.74 | 2 | -0.01112 | -6.661 | | 380304 |
| 61 | 891 | 905 | (K)lndldfgnsdaislr(R) | 100% | | 835.3815 | 1,668.75 | 2 | 0.001877 | 1.124 | | 184432 |
| 62 | 891 | 905 | (K)lndldfgnsdaislr(R) | 100% | | 835.375 | 1,668.74 | 2 | -0.01112 | -6.661 | | 190283 |
| 63 | 891 | 905 | (K)lndldfgnsdaislr(R) | 100% | | 835.3815 | 1,668.75 | 2 | 0.001877 | 1.124 | 2,889.98 | 139962 |
| 64 | 891 | 905 | (K)lndldfgnsdaislr(R) | 100% | | 835.3829 | 1,668.75 | 2 | 0.004677 | 2.801 | | 185610 |
| 65 | 891 | 906 | (K)lndldfgnsdaislrr(A) | 100% | | 610.6202 | 1,828.84 | 3 | 0.003081 | 1.684 | | 4488880 |
| 66 | 891 | 906 | (K)lndldfgnsdaislrr(A) | 100% | | 610.6201 | 1,828.84 | 3 | 0.002661 | 1.454 | | 4484540 |
| 67 | 891 | 906 | (K)lndldfgnsdaislrr(A) | 100% | | 610.6197 | 1,828.84 | 3 | 0.001461 | 0.7986 | | 1722020 |
| 68 | 891 | 906 | (K)lndldfgnsdaislrr(A) | 100% | | 610.6196 | 1,828.84 | 3 | 0.001071 | 0.5855 | | 489100 |
| 69 | 891 | 906 | (K)lndldfgnSdaislrr(A) | 100% | | 610.2885 | 1,827.84 | 3 | 0.004746 | 2.595 | | 4484540 |
| 70 | 891 | 906 | (K)lndldfgnsdaislrr(A) | 100% | | 610.6201 | 1,828.84 | 3 | 0.002541 | 1.389 | | 490114 |
| 71 | 891 | 906 | (K)lndldfgnSdaislrr(A) | 100% | | 610.2885 | 1,827.84 | 3 | 0.004746 | 2.595 | 2,253.46 | 540078 |
| 72 | 891 | 906 | (K)lndldfgnSdaislrr(A) | 100% | | 610.2885 | 1,827.84 | 3 | 0.004746 | 2.595 | | 1722020 |
| 73 | 891 | 906 | (K)lndldfgnSdaislrr(A) | 100% | | 610.2885 | 1,827.84 | 3 | 0.004746 | 2.595 | | 4488880 |
| 74 | 891 | 906 | (K)lndldfgnSdaislrr(A) | 100% | | 610.2885 | 1,827.84 | 3 | 0.004746 | 2.595 | | 489100 |
| 75 | 896 | 905 | (D)fgnsdaislr(R) | 100% | | 547.2577 | 1,092.50 | 2 | 0.001588 | 1.452 | 1,736.41 | 510486 |
| 76 | 905 | 925 | (L)rraSda**t**lgsidfgddeniak(R) | 100% | Phospho (+81) | 787.0007 | 2,357.98 | 3 | 0.009083 | 3.85 | | 1085390 |
| 77 | 906 | 925 | (R)ra**s**datlgsidfgddeniak(R) | 100% | Phospho (+81) | 733.9686 | 2,198.88 | 3 | 0.004808 | 2.185 | 2,655.40 | 1028460 |
| 78 | 906 | 925 | (R)ra**s**datlgsidfgddeniak(R) | 100% | Phospho (+81) | 733.9677 | 2,198.88 | 3 | 0.002198 | 0.999 | | 1166980 |
| 79 | 906 | 925 | (R)ra**s**datlgsidfgddeniak(R) | 100% | Phospho (+81) | 733.9675 | 2,198.88 | 3 | 0.001598 | 0.7263 | | 2080180 |
| 80 | 906 | 925 | (R)ra**s**datlgSidfgddeniak(R) | 100% | Phospho (+81) | 733.6373 | 2,197.89 | 3 | 0.007943 | 3.612 | | 1470070 |
| 81 | 906 | 925 | (R)ra**s**datlgSidfgddeniak(R) | 100% | Phospho (+81) | 733.6373 | 2,197.89 | 3 | 0.007943 | 3.612 | | 2080180 |
| 82 | 906 | 925 | (R)ra**s**datlgSidfgddeniak(R) | 100% | Phospho (+81) | 733.6373 | 2,197.89 | 3 | 0.007943 | 3.612 | | 1166980 |
| 83 | 906 | 925 | (R)ra**s**datlgSidfgddeniak(R) | 100% | Phospho (+81) | 733.6373 | 2,197.89 | 3 | 0.007943 | 3.612 | | 687417 |
| 84 | 906 | 925 | (R)ra**s**datlgSidfgddeniak(R) | 100% | Phospho (+81) | 733.6373 | 2,197.89 | 3 | 0.007943 | 3.612 | | 949641 |
| 85 | 906 | 925 | (R)ra**s**datlgSidfgddeniak(R) | 100% | Phospho (+81) | 733.6373 | 2,197.89 | 3 | 0.007943 | 3.612 | | 1569660 |
| 86 | 906 | 925 | (R)ra**s**datlgsidfgddeniak(R) | 100% | Phospho (+81) | 733.9678 | 2,198.88 | 3 | 0.002498 | 1.135 | | 1470070 |
| 87 | 906 | 925 | (R)ra**s**datlgSidfgddeniak(R) | 100% | Phospho (+81) | 733.6373 | 2,197.89 | 3 | 0.007943 | 3.612 | | 1126870 |
| 88 | 906 | 925 | (R)ra**s**datlgsidfgddeniak(R) | 100% | Phospho (+81) | 733.968 | 2,198.88 | 3 | 0.002978 | 1.354 | | 357632 |
| 89 | 906 | 925 | (R)rasda**t**lgsidfgddeniak(R) | 100% | Phospho (+81) | 733.9679 | 2,198.88 | 3 | 0.002798 | 1.272 | | 365235 |
| 90 | 906 | 925 | (R)ra**s**datlgSidfgddeniak(R) | 100% | Phospho (+81) | 733.6373 | 2,197.89 | 3 | 0.007943 | 3.612 | | 743417 |
| 91 | 906 | 925 | (R)ra**s**datlgSidfgddeniak(R) | 100% | Phospho (+81) | 733.6373 | 2,197.89 | 3 | 0.007943 | 3.612 | | 357632 |
| 92 | 906 | 925 | (R)ra**s**datlgsidfgddeniak(R) | 100% | Phospho (+81) | 733.9677 | 2,198.88 | 3 | 0.002048 | 0.9308 | | 1126870 |
| 93 | 906 | 925 | (R)ra**s**datlgSidfgddeniak(R) | 100% | Phospho (+81) | 733.6373 | 2,197.89 | 3 | 0.007943 | 3.612 | | 2.17E+07 |
| 94 | 906 | 925 | (R)ra**s**datlgSidfgddeniak(R) | 100% | Phospho (+81) | 733.6373 | 2,197.89 | 3 | 0.007943 | 3.612 | | 4702950 |
| 95 | 906 | 925 | (R)ra**s**datlgsidfgddeniak(R) | 100% | Phospho (+81) | 733.968 | 2,198.88 | 3 | 0.003098 | 1.408 | | 4702950 |
| 96 | 906 | 925 | (R)ra**s**datlgsidfgddeniak(R) | 100% | Phospho (+81) | 733.9689 | 2,198.89 | 3 | 0.005888 | 2.676 | | 6184340 |
| 97 | 906 | 925 | (R)ra**s**datlgSidfgddeniak(R) | 100% | Phospho (+81) | 733.6373 | 2,197.89 | 3 | 0.007943 | 3.612 | | 613968 |
| 98 | 906 | 925 | (R)ra**s**datlgsidfgddeniak(R) | 100% | Phospho (+81) | 733.9686 | 2,198.88 | 3 | 0.004808 | 2.185 | 2,647.26 | 1827130 |
| 99 | 906 | 925 | (R)rasda**t**lgSidfgddeniak(R) | 100% | Phospho (+81) | 733.6373 | 2,197.89 | 3 | 0.007943 | 3.612 | | 365235 |
| 100 | 906 | 925 | (R)ra**s**datlgsidfgddeniak(R) | 100% | Phospho (+81) | 733.9689 | 2,198.89 | 3 | 0.005888 | 2.676 | | 3170290 |
| 101 | 906 | 925 | (R)ra**s**datlgsidfgddeniak(R) | 100% | Phospho (+81) | 733.9686 | 2,198.88 | 3 | 0.004808 | 2.185 | | 1.67E+07 |
| 102 | 906 | 925 | (R)ra**s**datlgSidfgddeniak(R) | 100% | Phospho (+81) | 733.6373 | 2,197.89 | 3 | 0.007943 | 3.612 | | 6184340 |
| 103 | 906 | 925 | (R)ra**s**datlgSidfgddeniak(R) | 100% | Phospho (+81) | 733.6373 | 2,197.89 | 3 | 0.007943 | 3.612 | | 3170290 |
| 104 | 906 | 925 | (R)ra**s**datlgsidfgddeniak(R) | 100% | Phospho (+81) | 733.968 | 2,198.88 | 3 | 0.002978 | 1.354 | | 743417 |
| 105 | 906 | 925 | (R)rasdatlgSidfgddeniak(R) | 100% | | 706.9794 | 2,117.92 | 3 | 0.0007514 | 0.3546 | 2,224.69 | 233021 |
| 106 | 906 | 925 | (R)ra**s**datlgsidfgddeniak(R) | 100% | Phospho (+81) | 733.9701 | 2,198.89 | 3 | 0.009398 | 4.272 | | 2.17E+07 |
| 107 | 906 | 925 | (R)ra**s**datlgSidfgddeniak(R) | 100% | Phospho (+81) | 733.6373 | 2,197.89 | 3 | 0.007943 | 3.612 | | 225374 |
| 108 | 906 | 925 | (R)rasda**t**lgSidfgddeniak(R) | 100% | Phospho (+81) | 733.6373 | 2,197.89 | 3 | 0.007943 | 3.612 | | 202137 |
| 109 | 906 | 925 | (R)ra**s**datlgsidfgddeniak(R) | 100% | Phospho (+81) | 733.9684 | 2,198.88 | 3 | 0.004418 | 2.008 | 2,557.13 | 772099 |
| 110 | 906 | 925 | (R)ra**s**datlgsidfgddeniak(R) | 100% | Phospho (+81) | 733.9686 | 2,198.88 | 3 | 0.004898 | 2.226 | | 613968 |
| 111 | 906 | 925 | (R)ra**s**datlgSidfgddeniak(R) | 100% | Phospho (+81) | 733.6373 | 2,197.89 | 3 | 0.007943 | 3.612 | | 1.67E+07 |
| 112 | 906 | 925 | (R)ra**s**datlgsidfgddeniak(R) | 100% | Phospho (+81) | 733.9683 | 2,198.88 | 3 | 0.003878 | 1.763 | 2,524.50 | 247439 |
| 113 | 906 | 925 | (R)rasda**t**lgsidfgddeniak(R) | 100% | Phospho (+81) | 733.9679 | 2,198.88 | 3 | 0.002798 | 1.272 | 2,540.72 | 226097 |
| 114 | 906 | 925 | (R)rasdatlgsidfgddeniak(R) | 100% | | 707.3119 | 2,118.91 | 3 | 0.001126 | 0.5313 | 2,224.69 | 233021 |
| 115 | 906 | 925 | (R)rasda**t**lgSidfgddeniak(R) | 100% | Phospho (+81) | 733.6373 | 2,197.89 | 3 | 0.007943 | 3.612 | | 261005 |
| 116 | 906 | 925 | (R)ra**s**datlgSidfgddeniak(R) | 100% | Phospho (+81) | 733.6373 | 2,197.89 | 3 | 0.007943 | 3.612 | | 184442 |
| 117 | 906 | 925 | (R)ra**s**datlgsidfgddeniak(R) | 100% | Phospho (+81) | 733.9684 | 2,198.88 | 3 | 0.004298 | 1.954 | 2,508.15 | 158016 |
| 118 | 906 | 925 | (R)ra**s**datlgSidfgddeniak(R) | 100% | Phospho (+81) | 733.6373 | 2,197.89 | 3 | 0.007943 | 3.612 | | 356811 |

Additional Notes:
*R region was 15N-labeled and thus, all N aroms are 15N.
The 15N-modification is not listed above for ease of reading.
*Phosphorylated residues are bolded and underlined.

**Supplementary Table 2. Mass spectrometry of tryptic peptides of PKA-treated Ycf1p R region**

| Entry # | Starting Residue | Ending Residue | Peptide sequence (Lowercase letters highlight modified amino acids)* | Probability | Modifications (mass difference)* | Observed | Actual Mass | Charge | Delta Da | Delta PPM | Retention Time | Total Ion Current |
|---|---|---|---|---|---|---|---|---|---|---|---|---|
| 119 | 906 | 925 | (R)ra**s**datlgsidfgddeniak(R) | 100% | Phospho (+81) | 733.968 | 2,198.88 | 3 | 0.002978 | 1.354 | 2,532.57 | 214862 |
| 120 | 906 | 925 | (R)rasda**t**lgsidfgddeniak(R) | 100% | Phospho (+81) | 733.9683 | 2,198.88 | 3 | 0.003878 | 1.763 | 2,516.38 | 195614 |
| 121 | 906 | 925 | (R)rasda**t**lgSidfgddeniak(R) | 100% | Phospho (+81) | 733.6373 | 2,197.89 | 3 | 0.007943 | 3.612 | | 205366 |
| 122 | 906 | 925 | (R)ra**s**datlgsidfgddeniak(R) | 100% | Phospho (+81) | 733.9682 | 2,198.88 | 3 | 0.003698 | 1.681 | | 356811 |
| 123 | 906 | 925 | (R)rasda**t**lgsidfgddeniak(R) | 100% | Phospho (+81) | 733.9675 | 2,198.88 | 3 | 0.001598 | 0.7263 | 2,687.91 | 260750 |
| 124 | 906 | 926 | (R)ra**s**datlgsidfgddeniakr(E) | 100% | Phospho (+81) | 787.3317 | 2,358.97 | 3 | 0.004868 | 2.063 | | 1147420 |
| 125 | 906 | 926 | (R)ra**s**datlgsidfgddeniakr(E) | 100% | Phospho (+81) | 590.7498 | 2,358.97 | 4 | 0.001782 | 0.7549 | | 645858 |
| 126 | 906 | 926 | (R)rasda**t**lgsidfgddeniakr(E) | 100% | Phospho (+81) | 787.331 | 2,358.97 | 3 | 0.002858 | 1.211 | | 431061 |
| 127 | 906 | 926 | (R)rasda**t**lgSidfgddeniakr(E) | 100% | Phospho (+81), | 590.5009 | 2,357.97 | 4 | 0.003217 | 1.364 | | 450579 |
| 128 | 906 | 926 | (R)ra**s**datlgSidfgddeniakr(E) | 100% | Phospho (+81) | 590.5009 | 2,357.97 | 4 | 0.003217 | 1.364 | | 645858 |
| 129 | 906 | 926 | (R)ra**s**datlgSidfgddeniakr(E) | 100% | Phospho (+81) | 787.0007 | 2,357.98 | 3 | 0.009083 | 3.85 | | 1147420 |
| 130 | 906 | 926 | (R)ra**s**datlgSidfgddeniakr(E) | 100% | Phospho (+81) | 787.0007 | 2,357.98 | 3 | 0.009083 | 3.85 | | 1806830 |
| 131 | 906 | 926 | (R)ra**s**datlgsidfgddeniakr(E) | 100% | Phospho (+81) | 787.3314 | 2,358.97 | 3 | 0.003938 | 1.668 | | 1806830 |
| 132 | 906 | 926 | (R)rasda**t**lgSidfgddeniakr(E) | 100% | Phospho (+81) | 787.0007 | 2,357.98 | 3 | 0.009083 | 3.85 | | 431061 |
| 133 | 906 | 926 | (R)ra**s**datlgSidfgddeniakr(E) | 100% | Phospho (+81) | 787.0007 | 2,357.98 | 3 | 0.009083 | 3.85 | | 1.26E+07 |
| 134 | 906 | 926 | (R)ra**s**datlgsidfgddeniakr(E) | 100% | Phospho (+81) | 787.3319 | 2,358.97 | 3 | 0.005558 | 2.355 | | 6469750 |
| 135 | 906 | 926 | (R)ra**s**datlgsidfgddeniakr(E) | 100% | Phospho (+81) | 787.3314 | 2,358.97 | 3 | 0.003938 | 1.668 | | 4487530 |
| 136 | 906 | 926 | (R)ra**s**datlgsidfgddeniakr(E) | 100% | Phospho (+81) | 787.332 | 2,358.97 | 3 | 0.005948 | 2.52 | | 1.23E+07 |
| 137 | 906 | 926 | (R)ra**s**datlgsidfgddeniakr(E) | 100% | Phospho (+81) | 787.332 | 2,358.97 | 3 | 0.005948 | 2.52 | | 1.26E+07 |
| 138 | 906 | 926 | (R)ra**s**datlgsidfgddeniakr(E) | 100% | Phospho (+81) | 787.3314 | 2,358.97 | 3 | 0.004058 | 1.719 | | 3600890 |
| 139 | 906 | 926 | (R)rasda**t**lgsidfgddeniakr(E) | 100% | Phospho (+81) | 590.7499 | 2,358.97 | 4 | 0.002342 | 0.9922 | | 450579 |
| 140 | 906 | 926 | (R)ra**s**datlgSidfgddeniakr(E) | 100% | Phospho (+81) | 787.0007 | 2,357.98 | 3 | 0.009083 | 3.85 | | 1.23E+07 |
| 141 | 906 | 926 | (R)ra**s**datlgSidfgddeniakr(E) | 100% | Phospho (+81) | 787.0007 | 2,357.98 | 3 | 0.009083 | 3.85 | | 6469750 |
| 142 | 906 | 926 | (R)rasda**t**lgSidfgddeniakr(E) | 100% | Phospho (+81) | 787.0007 | 2,357.98 | 3 | 0.009083 | 3.85 | | 730436 |
| 143 | 906 | 926 | (R)ra**s**datlgSidfgddeniakr(E) | 100% | Phospho (+81) | 787.0007 | 2,357.98 | 3 | 0.009083 | 3.85 | | 3600890 |
| 144 | 906 | 926 | (R)ra**s**datlgSidfgddeniakr(E) | 100% | Phospho (+81) | 787.0007 | 2,357.98 | 3 | 0.009083 | 3.85 | | 4487530 |
| 145 | 906 | 926 | (R)rasda**t**lgsidfgddeniakr(E) | 100% | Phospho (+81) | 787.33 | 2,358.97 | 3 | -0.0001424 | -0.06034 | | 730436 |
| 146 | 906 | 926 | (R)ra**s**datlgSidfgddeniakr(E) | 100% | Phospho (+81) | 590.5009 | 2,357.97 | 4 | 0.003217 | 1.364 | | 242401 |
| 147 | 906 | 926 | (R)rasdaTlg**s**idfgddeniakr(E) | 100% | Phospho (+81) | 787.0007 | 2,357.98 | 3 | 0.009083 | 3.85 | | 340843 |
| 148 | 906 | 926 | (R)rasda**t**lgSidfgddeniakr(E) | 100% | Phospho (+81) | 787.0007 | 2,357.98 | 3 | 0.009083 | 3.85 | | 125217 |
| 149 | 906 | 926 | (R)ra**s**datlgSidfgddeniakr(E) | 100% | Phospho (+81) | 787.0007 | 2,357.98 | 3 | 0.009083 | 3.85 | | 290840 |
| 150 | 906 | 926 | (R)ra**s**datlgsidfgddeniakr(E) | 100% | Phospho (+81) | 787.3304 | 2,358.97 | 3 | 0.001178 | 0.499 | | 290840 |
| 151 | 906 | 926 | (R)ra**s**datlgsidfgddeniakr(E) | 100% | Phospho (+81) | 590.75 | 2,358.97 | 4 | 0.002582 | 1.094 | 2,183.03 | 267674 |
| 152 | 906 | 926 | (R)rasda**t**lgsidfgddeniakr(E) | 100% | Phospho (+81) | 787.3309 | 2,358.97 | 3 | 0.002558 | 1.084 | | 340843 |
| 153 | 906 | 926 | (R)ra**s**datlgSidfgddeniakr(E) | 100% | Phospho (+81) | 787.0007 | 2,357.98 | 3 | 0.009083 | 3.85 | | 2127560 |
| 154 | 906 | 926 | (R)rasda**t**lgsidfgddeniakr(E) | 100% | Phospho (+81) | 787.3303 | 2,358.97 | 3 | 0.0006376 | 0.2702 | | 125217 |
| 155 | 906 | 926 | (R)ra**s**datlgsidfgddeniakr(E) | 100% | Phospho (+81) | 787.331 | 2,358.97 | 3 | 0.002858 | 1.211 | | 2127560 |
| 156 | 906 | 926 | (R)rasda**t**lgsidfgddeniakr(E) | 100% | Phospho (+81) | 787.3311 | 2,358.97 | 3 | 0.003008 | 1.274 | 2,163.78 | 1328110 |
| 157 | 907 | 925 | (R)asdatlgsidfgddeniak(R) | 100% | | 653.9494 | 1,958.83 | 3 | 0.002866 | 1.463 | 2,681.32 | 113664 |
| 158 | 907 | 925 | (R)asdatlgsidfgddeniak(R) | 100% | | 653.9494 | 1,958.83 | 3 | 0.002866 | 1.463 | 2,673.20 | 93,753.00 |
| 159 | 907 | 926 | (R)asdatlgsidfgddeniakr(E) | 100% | | 707.3121 | 2,118.91 | 3 | 0.001726 | 0.8143 | | 1167480 |
| 160 | 907 | 926 | (R)asdaTlgsidfgddeniakr(E) | 100% | | 706.9794 | 2,117.92 | 3 | 0.0007514 | 0.3546 | | 1167480 |
| 161 | 907 | 926 | (R)asdatlgsidfgddeniakr(E) | 100% | | 707.3124 | 2,118.92 | 3 | 0.002626 | 1.239 | | 447605 |
| 162 | 907 | 926 | (R)asdaTlgsidfgddeniakr(E) | 100% | | 706.9794 | 2,117.92 | 3 | 0.0007514 | 0.3546 | | 447605 |
| 163 | 907 | 926 | (R)asdaTlgsidfgddeniakr(E) | 100% | | 706.9794 | 2,117.92 | 3 | 0.0007514 | 0.3546 | | 597673 |
| 164 | 907 | 926 | (R)asdatlgsidfgddeniakr(E) | 100% | | 707.3121 | 2,118.91 | 3 | 0.001816 | 0.8568 | 2,232.80 | 689329 |

Additional Notes:
*R region was 15N-labeled and thus, all N aroms are 15N.
The 15N-modification is not listed above for ease of reading.
*Phosphorylated residues are bolded and underlined.
